# Specification of distinct cell types in a sensory-adhesive organ for metamorphosis in the *Ciona* larva

**DOI:** 10.1101/2023.05.02.539060

**Authors:** Christopher J. Johnson, Florian Razy-Krajka, Fan Zeng, Katarzyna M. Piekarz, Shweta Biliya, Ute Rothbächer, Alberto Stolfi

**Author notes:** Equal contributions (Christopher J. Johnson, Florian Razy-Krajka).

## Abstract

The papillae of tunicate larvae contribute sensory, adhesive, and metamorphosis-regulating functions that are crucial for the biphasic lifestyle of these marine, non-vertebrate chordates. We have identified additional molecular markers for at least five distinct cell types in the papillae of the model tunicate *Ciona,* allowing us to further study the development of these organs. Using tissue-specific CRISPR/Cas9-mediated mutagenesis and other molecular perturbations, we reveal the roles of key transcription factors and signaling pathways that are important for patterning the papilla territory into a highly organized array of different cell types and shapes. We further test the contributions of different transcription factors and cell types to the production of the adhesive glue that allows for larval attachment during settlement, and to the processes of tail retraction and body rotation during metamorphosis. With this study, we continue working towards connecting gene regulation to cellular functions that control the developmental transition between the motile larva and sessile adult of *Ciona*.

## Introduction

Tunicates, the sister group to the vertebrates, comprise a diverse group of marine non-vertebrate chordates (Fodor et al., 2021; Lemaire, 2011). Most tunicate species are classified in the order Ascidiacea, commonly known as ascidians (Satoh, 2013), although phylogenetic evidence suggests this is not a monophyletic group within Tunicata (DeBiasse et al., 2020; Delsuc et al., 2018; Kocot et al., 2018). The majority of ascidians have a biphasic life cycle that alternates between a swimming larva and a sessile adult. The larva functions exclusively to disperse the species, not feeding until it has found a suitable location on which to settle and trigger metamorphosis (Karaiskou et al., 2015).

Recent work has started to reveal the cellular and molecular basis of larval settlement and metamorphosis. Key to the process of settlement and metamorphosis are the papillae, which comprise a set of three anterior sensory/adhesive organs in the laboratory model species of the genus *Ciona* and a majority of other ascidian genera as well (**Figure 1**)(Caicci et al., 2010; Torrence and Cloney, 1983; Turon, 1991; Zeng et al., 2019b). The papillae are composed of a few different cell types that have been characterized by both electron and fluorescence microscopy (Dolcemascolo et al., 2009; Pennati et al., 2009; Pennati et al., 2007; Zeng et al., 2019b). Several cells appear to secrete the “glue” or bioadhesive material required for the attachment of the larva to the substrate, termed “collocytes” (Zeng et al., 2019a; Zeng et al., 2019b). Other cells are clearly neuronal (four ciliated neurons per papilla)(Zeng et al., 2019b) and are required to trigger the onset of metamorphosis (Sakamoto et al., 2022), which was also recently shown to depend on mechanical stimulation of the papillae (Wakai et al., 2021). Finally, at the very center of each papilla are four “Axial Columnar Cells” (ACCs), which have been suggested to possess chemosensory and contractile properties (Poncelet and Shimeld, 2020; Poncelet et al., 2022; Turon, 1991). Although they have been called papilla “sensory cells” or “neurons”, they are not innervated and have little structural and molecular overlap with the other two cell types. Furthermore, single-cell RNA sequencing revealed that they do not express genes typically associated with neuronal function (Sharma et al., 2019).

**Figure 1.**
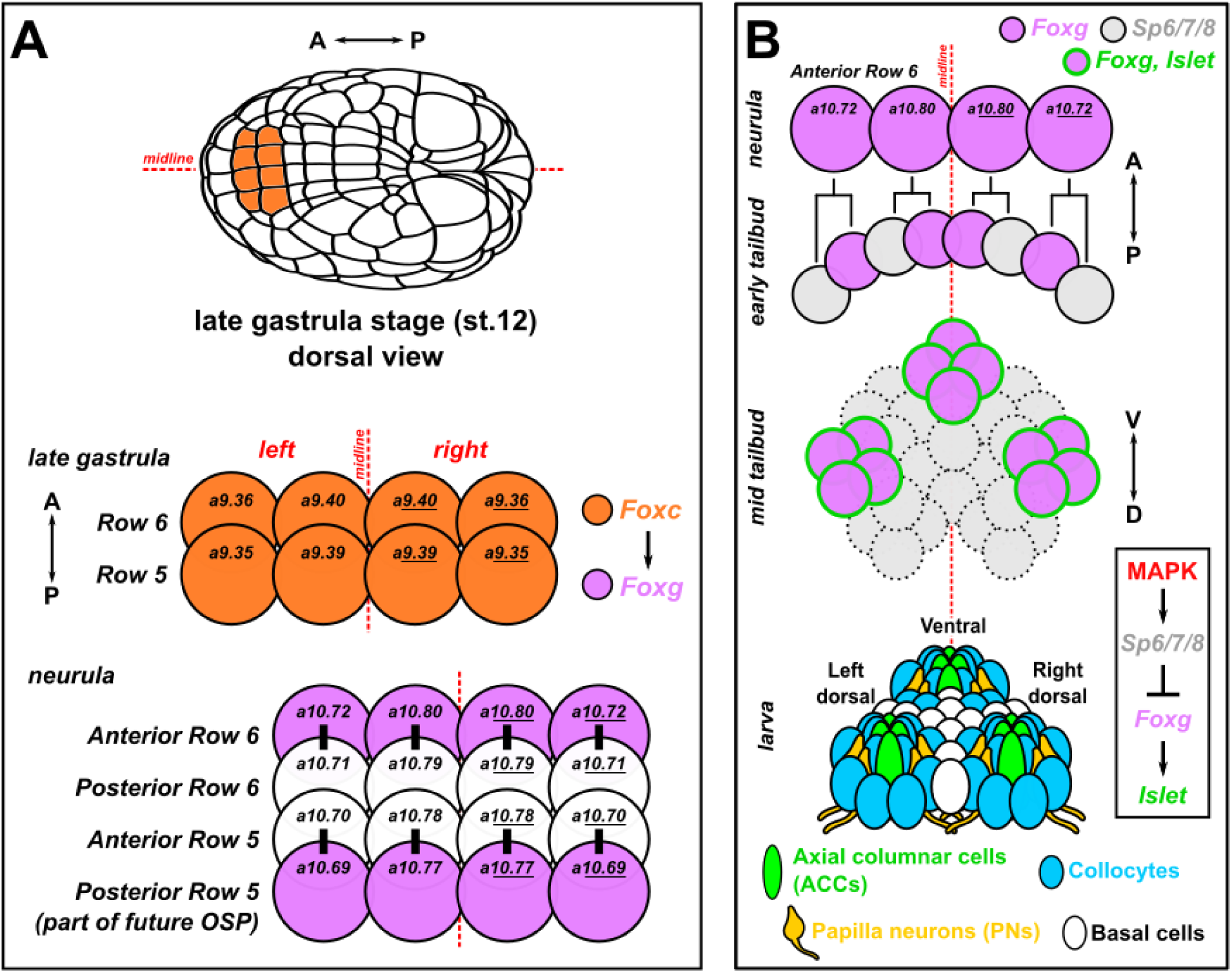
Development of the papillae of Ciona. A) Diagram showing the early cell lineages that give rise to the papillae. The papillae invariantly derive from Foxc+ cells in the anterior neural plate, more specifically the anterior daughter cells of “Row 6” of the neural plate, which activate *Foxg* downstream of Foxc. *Foxg* is also activated in the posterior daughter cells of “Row 5”, which go on to give rise to part of the oral siphon primordium (OSP). Numbers in each cell indicate their invariant identity according to the Conklin cell lineage nomenclature. Black bars indicate sibling cells born from the same mother cell. B) Diagram of what is currently known about the later lineage and fates of the *Foxg+* “Anterior Row 6” cells shown in panel A. As the cells divide mediolaterally, some cells upregulate *Sp6/7/8* and downregulate *Foxg* (grey cells). Those cells that maintain *Foxg* expression turn on *Islet* and coalesce as three clusters of cells (pink with green outline): one medial, more ventral cluster, and two left/right, more dorsal clusters. Later these three clusters organize the territory into the three protruding papillae of the larva, which contains several cell types described in detail by transmission electron microscopy (Zeng et al. 2019b). Dashed cell outlines indicate uncertain number/provenance of cells. A-P: anterior-posterior. D-V: dorsal-ventral. Lineages and gene networks are based mostly on Liu and Satou 2019, Nicol and Meinertzhagen 1988, Wagner and Levine 2012, and Wagner et al. 2014.

In *Ciona,* previous work had established that the three papillae likely arise from three clusters of *Foxg+/Islet+* cells arranged roughly as a triangle-two dorsal clusters (left and right) and single ventral cluster (Liu and Satou, 2019; Wagner et al., 2014). Although *Foxg* is initially activated in an entire row of cells at the very anterior of the neural plate, Sp6/7/8 (also known as Zfp220 or Buttonhead) is required to refine this swath of expression down to three “spots” of *Foxg,* which required for expression of *Islet* in these cell clusters (**Figure 1**)(Liu and Satou, 2019). MEK/ERK (e.g. MAPK) signaling also appears to play an important role in this refinement, as treatment with the MEK inhibitor U0126 results in a “U”-shaped band of *Islet* expression instead of three discrete foci (**Figure 1**)(Wagner et al., 2014). Similarly, BMP inhibition also causes a similar “U-shape” swath of *Foxg/Islet* expression, resulting in a single protrusion instead of the normal three, termed the “*cyrano*” phenotype (Liu et al., 2023; Roure et al., 2022). However, it has not been shown how these early specification events connect to the final cell type diversity and arrangement of the papillae.

Here we describe novel genetic markers and reporter constructs that allowed us to visualize each of the different cell type of the papillae, and follow their development upon various molecular perturbations targeting specific transcription factors or signaling pathways. We show that different transcription factors contribute to the specification of the different cell types, and that cell-cell signaling in the FGF/MAPK and Delta/Notch pathways are crucial for patterning and arranging these cells in the three papillae. Altering papilla development in different ways contributes to different processes of post-settlement larval body plan rearrangements, revealing the complex molecular and cellular underpinning of tunicate larval metamorphosis.

## Methods

### Ciona handling

*Ciona robusta (intestinalis Type A)* were shipped from San Diego (M-REP), while *Ciona intestinalis (Type B)* were shipped from Roscoff Biological Station, France. Eggs were fertilized *in vitro,* dechorionated, and electroporated following established protocols (Christiaen et al., 2009a, b; Kari et al., 2016). Unc-76 tags were used as a default for fluorescent proteins (FPs) for optimal cell labeling as previously described (Stolfi and Levine, 2011), which excludes the FPs from the nucleus and ensures transport down axons. Typically 40-100 μg of untagged or Unc-76-tagged FP plasmids and 10-35 μg of histone (H2B) fusion FP plasmids was used per 700 μl of electroporation solution. For CRISPR, typically 35-40 μg of Cas9 plasmid and 25-40 μg of each gRNA plasmid was used per 700 μl of electroporation solution, except when validating sgRNAs (see further below). Precise electroporation mixes for given perturbation experiments and controls are specified in the **Supplemental Sequence File.** *C. robusta* embryos were raised at 20°C and *C. intestinalis* embryos were raised at 18°C, unless otherwise specified. For U0126 treatment, U0126 stock solution resuspended in DMSO was diluted to 10 µM final concentration in artificial seawater prior to transferring embryos at 7.5 hpf. Negative control embryos were transferred to seawater with the equivalent volume of DMSO vehicle.

### Fixation, staining, and imaging

Embryos and larvae were fixed for fluorescent protein imaging in MEM-FA fixation solution (3.7% formaldehyde, 0.1 M MOPS pH 7.4, 0.5 M NaCl, 1 mM EGTA, 2 mM MgSO4, 0.1% Triton-X100), rinsed in 1X PBS, 0.4% Triton-X100, 50 mM NH4Cl and 1X PBS, 0.1% Triton-X100. For mRNA *in situ* hybridization, embryos/larvae were fixed in MEM-PFA fixation solution (4% paraformaldehyde, 0.1 M MOPS pH 7.4, 0.5 M NaCl, 1 mM EGTA, 2 mM MgSO4, 0.05% Tween-20) and *in situ* hybridization was carried out as previously described (Ikuta and Saiga, 2007; Stolfi et al., 2011). All probe template sequences are shown in the **Supplemental Sequence File**. Immunolabeling of β-galactosidase and mCherry (alone or in conjunction with mRNA *in situ* hybridization) was carried out as previously described (Beh et al., 2007), on embryos/larvae using mouse anti-β-gal (Promega catalog number Z3781, 1:1000) and rabbit anti-mCherry (BioVision, accession number ACY24904, 1:500) primary antibodies. Specimens were imaged on Leica DM IL LED or DMI8 inverted epifluorescence microscopes, with maximum Z projection processing and cell measurements performed in LAS X.

PNA staining was carried out on 4% PFA fixed larvae, using Tris-buffered saline (pH 8.0) supplemented with 5 mM CaCl2 and 0.1% Triton X-100 (TBS-T). Unspecific background was blocked by 3% BSA in TBS-T for 2 hours at room temperature. Biotinylated Peanut Agglutinin (PNA; B-1075, Vector Laboratory) was diluted in BSA-TBS-T to a final concentration of 25 μg/ml and applied to the specimen overnight at 4 °C. After several washes in TBS-T over 2 hours, larvae were incubated for 1 hour in fluorescent streptavidin (SA-5006, Vector Laboratory) diluted 1:300 in BSA-TBS-T at room temperature. PNA stainings were imaged using a Leica SP5 II confocal scanning microscope. Stacks were acquired sequentially and z-projected. Images were analyzed with ImageJ (Version 1.52 h).

### CRISPR/Cas9 sgRNA design and validation

The Cas9 (Stolfi et al., 2014) and Cas9::Geminin-Nterminus (Song et al., 2022) protein-coding sequences have been described before. Single-chain guide RNAs (sgRNAs) were designed using the CRISPOR website (Haeussler et al., 2016)(crispor.tefor.net). Those sgRNAs with high Doench ‘16 score, high MIT specificity score, and not spanning known SNPs were selected for testing. Validation of sgRNAs was performed by co-electroporation 25 μg of *Eef1a>Cas9* or *Eef1a>Cas9::Geminin-Nterminus* and 75 μg of the sgRNA plasmid, per 700 μl of total electroporation volume. Genomic DNA was extracted from pooled larvae electroporated with a single sgRNA, using the QIAamp DNA micro kit (Qiagen). PCR products spanning each sgRNA target site were amplified from the corresponding genomic DNA, with primers designed so that the amplicon was to be 150-450 bp in size. Amplicons were purified by QIAquick PCR purification kit (Qiagen) and submitted for Amplicon-EZ Illumina-based sequencing by Azenta/Genewiz (New Jersey, USA), which returned mutagenesis rates and indel plots.

### RNA sequencing and analysis

Single-cell RNA sequencing (scRNAseq) data from Cao et al. 2019 were re-analyzed in Seurat (Satija et al., 2015). Combined larva stage data was clustered and plotted using 30 dimensions (**Supplemental Figure 1A**). Clusters 3 and 33 were determined to contain papilla cell types and were re-clustered separately, also using 30 dimensions (**Supplemental Figure 1B**). Differential gene expression plots (**Supplemental Figure 1C**) and tables (**Supplemental Table 1**) were explored to find candidate papilla cell type markers, to be confirmed by *in situ* hybridization (**Supplemental Figure 1D**) and/or reporter plasmids. All code and Seurat files can be downloaded from: https://osf.io/sc7pr/

Bulk RNA integrity numbers were determined using the Agilent Bioanalyzer RNA 6000 Nano kit and used as a QC measure. All samples with RINs over 7 were used for library preparation. mRNA was enriched using the NEBNext Poly(A) mRNA isolation module and Illumina compatible libraries were prepared using the NEBNext Ultra II RNA directional library preparation kit. QC on the libraries was performed on the Agilent Bioanalyzer 2100 and concentrations were determined fluorometrically. The libraries were then pooled and sequenced on the NovaSeq 6000 with an SP Flow Cell to get PE100bp reads.

The RNA-seq raw files were analyzed in Galaxy hub (usegalaxy.org)(Afgan et al., 2022). Firstly, the raw fastq files were inspected using FastQC Read Quality Reports (Galaxy Version 0.73+galaxy0) and MultiQC (Galaxy Version 1.11+galaxy0). The reads were then filtered and trimmed with Cutadapt (Galaxy Version 4.0+galaxy0). The minimum read length was set to 20 and the reads that did not meet the quality cutoff of 20 were discarded. Then, FastQC and MultiQC were used again to assess the resulting files after filtering and trimming. Next, the technical replicates were combined and used as the input to the mapping tool (RNA STAR, Galaxy Version 2.7.8a+galaxy0, length of the SA pre-indexing string of 12), together with the custom *Ciona* reference genome sequence and gene model files (KY21, both obtained from the Ghost Database; http://ghost.zool.kyoto-u.ac.jp/download_ht.html)(Satou et al., 2022). The counts were generated using featureCounts (Galaxy Version 2.0.1+galaxy2; minimum mapping quality per gene was set to 10). Lastly, the differential gene expression analysis (**Supplemental Table 2**) was performed with DESeq2 (Galaxy Version 2.11.40.7+galaxy1). KY21 gene models were linked to KH gene models using the Ciona Gene Model Converter application https://github.com/katarzynampiekarz/ciona_gene_model_converter (Piekarz and Stolfi, *under review).* Raw sequencing reads available from the SRA database under accession PRJNA949791. Analysis code and files can be found at: https://osf.io/wzrdk/

### Quantification of ACC length in *Villin* CRISPR larvae

Larvae subjected to papilla-specific knockout of *Villin* (using *Foxc>Cas9,* see **Supplemental Sequence File** for detailed electroporation recipe) and negative control larvae were fixed at 17 hpf, 20°C and mounted as above. *CryBG>Unc-76::GFP+* cells were imaged on a Leica DMI8 inverted epifluorescence microscope and the greatest distance between the apical and basal extremities of each GFP+ papilla was measured in LAS X, based on visible GFP fluorescence at a given focal plane.

## Results

### Identification of novel markers and reporters for specific cell types in the papillae

We searched *Ciona robusta* (i.e. *intestinalis* Type A) whole-larva single-cell RNA sequencing (scRNAseq) data (Cao et al., 2019) for evidence of the cell types described by transmission electron microscopy (TEM) of the papillae (Zeng et al., 2019b). While a cell cluster annotated as “Palp Sensory Cells” (PSCs) appeared enriched for known markers of ACCs like *CryBG (KH.S605.3)* and *KH.C3.516* (Sharma et al., 2019; Shimeld et al., 2005), genes expressed in other papilla cell types were also enriched in this cluster as well, including *Sp6/7/8 (KH.C13.22)* (Liu and Satou, 2019; Wagner et al., 2014) and *Pou4 (KH.C2.42)*(Roure et al., 2022; Sakamoto et al., 2022). Re-analysis and reclustering of these data revealed novel potential markers for these different cell types within the cluster (**Supplemental Figure 1A-C, Supplemental Table 1**). We performed *in situ* mRNA hybridization for several of these PSC candidate markers in *C. robusta* larvae (**Supplemental Figure 1D**). As we had hoped, they appeared to label different cells in the papilla territory. Some appeared to label cells in the center of each papilla, while others were expressed in cells surrounding or on the outermost edges of each papilla. These vastly different expression patterns supported the idea of mixed cell identities in the PSC scRNAseq cluster.

To further confirm the expression patterns of these and other candidate markers, we made reporter plasmids from their upstream *cis*-regulatory sequences and electroporated these into *Ciona* embryos. None of the selected genes showed any appreciable homology to genes of known function in other organisms, but we reasoned that they might serve as useful markers for specific papilla cell types. First, a *KH.L96.43* reporter (“*L96.43>GFP*”) was expressed in cells surrounding and in between the three papillae (**Figure 2A**). Co-electroporation with the papilla-specific *Foxg>mCherry* reporter (Cao et al., 2019) showed clear, mutually-exclusive expression between the two reporters. We propose that L96.43 marks a population of “peri-papillary” and/or “inter-papillary” cells previously identified as “basal cells” that are part of the larger papilla region but excluded from the three protruding, *Foxg+* papillae *sensu stricto* (Zeng et al., 2019b).

**Figure 2.**
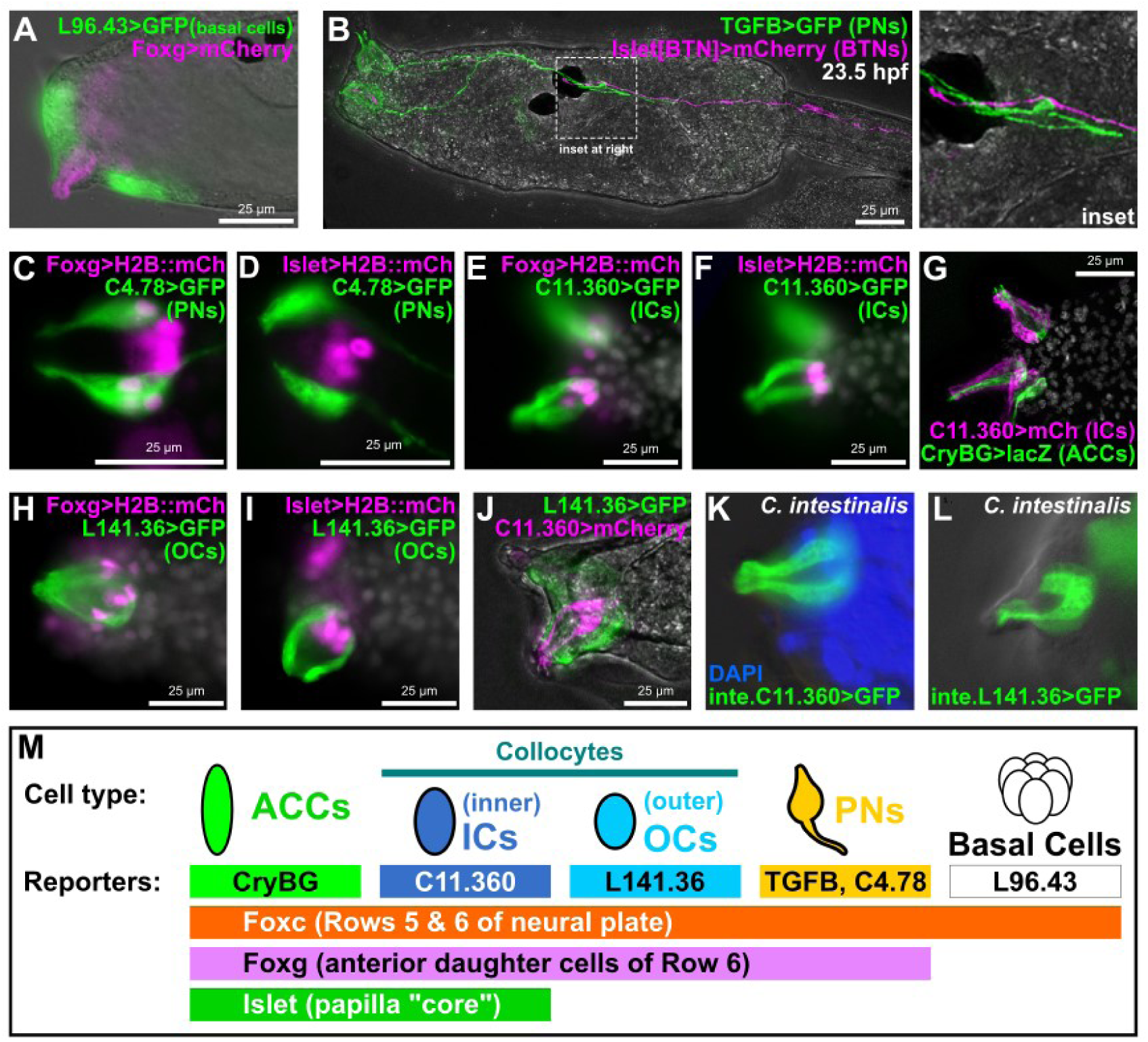
Novel genetic markers label distinct cell types of the papillae. A) GFP reporter plasmid (green) constructed using the *cis*-regulatory sequences from the *KH.L96.43* gene labels Basal Cells in between and surrounding the protruding papillae labeled by *Foxg* reporter plasmid (pink). B) *TGFB>GFP* reporter (green) labels Papilla Neurons (PNs), the axons of which make contacts with Bipolar Tail Neuron (BTN) axons labeled by a BTN-specific *Islet* reporter (pink), at 23.5 hours post-fertilization (hpf). C) A *KH.C4.78* reporter (*C4.78>GFP)* also labels PNs, which are also labeled by *Foxg>H2B::mCherry (mCh)* reporter (pink nuclei). D) Lack of overlap between expression of *C4.78>GFP* (green) and a papilla-specific *Islet* reporter plasmid (pink nuclei) showing that PNs do not arise from Islet+ cells. E,F) Co-electroporation of *C11.360>GFP* (green) with H2B::mCherry reporter plasmids (pink nuclei) indicates these cells come from Foxg-expressing cells that also express Islet. G) *C11.360>mCherry* reporter (pink) labels centrally-located “inner” collocytes (ICs) adjacent to Axial Columnar Cells (ACCs) labeled by *CryBG>LacZ* reporter (green). H,I) *L141.36>GFP* reporter (green) labels “outer” collocytes (OCs) that arise from *Foxg+* cells (pink nuclei) but do not express *Islet* (pink nuclei). J) ICs and OCs are distinct cells as there is no overlap between *C11.360* (green) and *L141.36* (pink) reporter plasmid expression. K) *Ciona intestinalis* (Type B) larva ICs labeled with a reporter plasmid made from the corresponding *cis*-regulatory sequence of the *C. intestinalis Chr11.1038* gene, orthologous to *C. robusta KH.C11.360.* L) *C. intestinalis* larva OCs labeled by a *Chr7.130* reporter, corresponding to the *C. robusta* ortholog *KH.L141.36.* M) Summary of the main marker genes and corresponding reporter plasmids used in this study to label different subsets of papilla progenitors and their derivative cell types. All GFP and mCherry reporters fused to the Unc-76 tag, unless specified (see methods and supplement for details). Weaker *Foxg -2863/-3* promoter used in panel B, all other *Foxg* reporters used the improved *Foxg -2863/+54* sequence instead. All *Islet* reporters shown correspond to the *Islet intron 1 + bpFOG>H2B::mCherry* plasmid. White channel shows either DAPI (nuclei) and/or larva outline in brightfield, depending on the panel. All *C. robusta* raised at 20°C to 18 hpf except: panel B (23.5 hpf); panels C-F (17 hpf); panels H-J (20 hpf). *C. intestinalis* raised at 18°C to 20-22 hpf.

Next, we further confirmed that the Papilla Neurons (PNs) are distinct from the ACCs (Zeng et al., 2019b). Previously identified as a potential PN marker by *in situ* hybridization (Razy-Krajka et al., 2014), a *TGFB* reporter clearly labeled PNs (**Figure 2B, Supplemental Figure 2A**), which are distinguished as the only papilla cell types bearing an axon. However, co-electroporation of TGFB reporter with an ACC-specific *CryBG* reporter (Shimeld et al., 2005) resulted in “cross-talk”, or cross-plasmid transvection (**Supplemental Figure 2B**). Indeed, other PN-specific reporters tested did not cross-talk with *CryBG*, including the previously published *Gnrh1* (Kusakabe et al., 2012), and the novel marker *KH.C4.78 (“C4.78>GFP”)*(**Supplemental Figure 2C-E**). Interestingly, PN axons continued to extend posteriorly during the swimming phase to contact the anterior axon branches of the Bipolar Tail Neurons (**Figure 2B**), which project their posterior axon branches to the very tip of the tail (Imai and Meinertzhagen, 2007). This hints at a potential mechanism for transducing sensory information from the papillae to the tail tip where tail retraction initiates, especially during later time points when larvae are competent to settle (Matsunobu and Sasakura, 2015).

Double electroporation with *KH.C4.78* and *Foxg* reporters (**Figure 2C, Supplemental Figure 2C**) revealed that, unlike the basal cells, PNs are specified from *Foxg+* cells in the papillae. However, co-electoporation with a papilla-specific *Islet* reporter plasmid also revealed that PNs are adjacent to but distinct from the central Islet+ “core” of each papilla (**Figure 2D**). In contrast, a *KH.C11.360* reporter (“*C11.360>GFP/mCherry”*) labeled cells that were both Foxg+ and Islet+, but were clearly not the ACCs (**Figure 2E-G**). The C11.360+ cells were adjacent to the ACCs but lacked the thin protrusions into the hyaline cap that are typical of the ACCs, and also lacked axons typical of the PNs. Therefore, these cells appear to be collocytes, proposed to be adhesive-secreting cells responsible for attachment to the substrate during larval settlement (Zeng et al., 2019b).

Previous characterization of the papillae by TEM described 12 collocytes in each papilla (Zeng et al., 2019b), yet the C11.360 reporter appeared to only label at most four cells per papilla. This suggested the existence of cryptic collocyte subtypes. In fact, those same TEM images showed certain qualitative differences in cytoplasmic contents between peripheral collocytes and the more central collocytes (Zeng et al., 2019b). Indeed, we identified another reporter, that of the gene *KH.L141.36 (“L141.36>GFP”),* that labeled *Foxg+* but *Islet-*negative cells that are at the periphery of each papilla but that are not PNs (**Figure 2H,I**). Co-electroporation of *L141.36* and *C11.360* reporters labelled mutually-exclusive groups of cells (**Figure 2J**). We propose that these respective reporters delineate more peripheral, or “outer” collocytes (OCs) vs. more central, or “inner” collocytes (ICs). Interestingly, *KH.L141.36* reporter expression was only visible at 20 hpf, not at 17 hpf like most of the other reporters described.

When using these *C. robusta* reporter plasmids to electroporate the closely related *C. intestinalis* (i.e. Type B) sourced from Roscoff, France (Pennati et al., 2015), we noticed that their expression was very weak (data not shown). This led us to re-cloning the orthologous sequences from the *C. intestinalis* Type B genome (Satou et al., 2021)(**Supplemental Sequence File**). Electroporation of Type B embryos with Type B-specific reporter plasmids resulted in much stronger, reliable expression (**Figure 2K,L**). This suggests relatively significant changes to the *cis-*regulatory sequences of these cell type-specific genes in these otherwise nearly indistinguishable cryptic species.

Although we also obtained additional reporters that labeled one or more different papilla cell types (**Supplemental Figure 2F,G**), we now had a full set of papilla cell type-specific marker genes and reporter plasmids for a deeper investigation of papilla patterning and development (**Figure 2M**). Finally, it is also important to note that some of these reporters also label cell types outside the papillae.

### Specification of ACCs, ICs, and OCs by Islet and Sp6/7/8 combinatorial logic

How are the cell types of the papillae (ACCs, ICs, OCs, and PNs) specified? *In situ* mRNA hybridization previously revealed partially overlapping expression territories of three genes encoding sequence-specific transcription factors (**Figure 3A**): a central domain of *Islet+* cells, surrounded by a ring of cells that express both *Islet* and *Sp6/7/8* (and *Emx,* though distinct from the earlier expression of *Emx* at neurula stages), and additional cells surrounding them expressing only *Sp6/7/8* (Wagner et al., 2014). Additionally, overexpression of *Islet* had been previously shown to generate a single large papilla expressing the ACC reporter *CryBG>GFP* (Wagner et al., 2014). We therefore asked whether these transcription factors might be patterning the papillae into an ordered array of cell types (**Figure 3A**).

**Figure 3.**
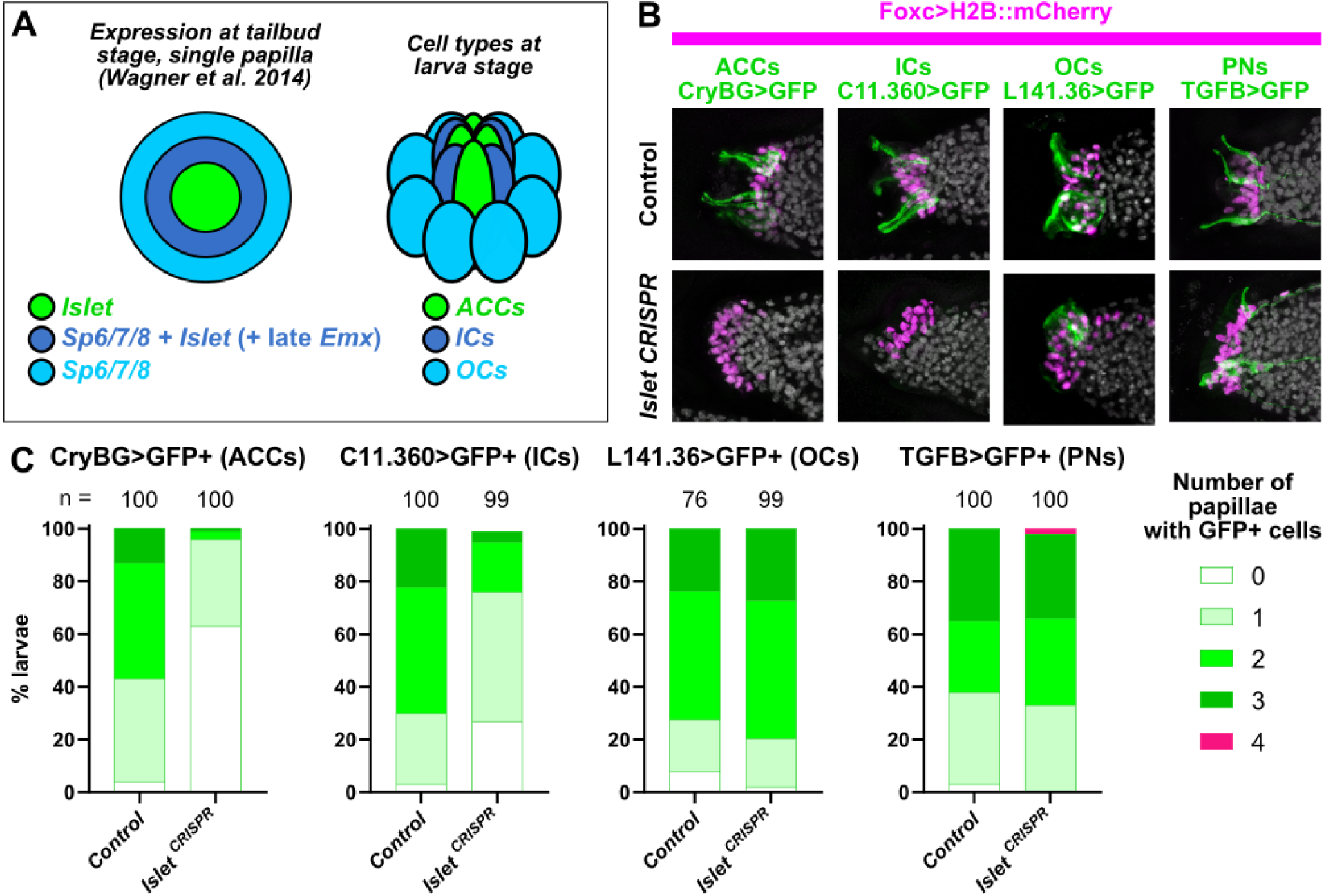
The transcription factor Islet is required for specification of ACCs and ICs. A) Diagram depicting a partially overlapping expression patterns of *Islet* and *Sp6/7/8,* as originally shown by *in situ* mRNA hybridizations (Wagner et al. 2014), and the correlation of these patterns with the later arrangement of ACCs, ICs, and OCs in the papillae. “Late” *Emx* expression in a ring of cells expressing both *Islet* and *Sp6/7/8* appears to be distinct from earlier *Emx* expression in *Foxg-negative* cells (see text and Supplemental Figure 2 for details). B) Papilla lineage-specific CRISPR/Cas9-mediated mutagenesis of *Islet* using *Foxc>Cas9* and a the *U6>Islet.2* single-chain guide RNA (sgRNA) plasmid shows reduction of larvae showing expression of reporters labeling ACCs and ICs, but not OCs or PNs. Results compared to a negative “control” condition using a negative control sgRNA (*U6>Control,* see text for details). Nuclei counterstained with DAPI (white). C) Scoring data for larvae represented in panel B. *Foxc>H2B::mCherry+* larvae were scored for quantity of papillae showing visible expression of the corresponding GFP reporter plasmid. Due to mosaic uptake or retention of the plasmids after electroporation, number of papillae with GFP fluorescence is variable and rarely seen in all three papillae even in control larvae. Normally larvae have three papilla (GFP+ or not), but some mutants have more/fewer than three. ACC/IC/OC sub-panels in panel B at 20 hpf/20°C, PN sub-panels at 21 hpf/20°C. Same applies to scoring data in Panel C.

First we asked, does Islet specify the centrally-located ACCs and ICs? To test this, we turned to tissue-specific CRISPR/Cas9-mediated mutagenesis (Stolfi et al., 2014). To knock out *Islet* in the papillae, we electroporated a previously validated sgRNA expression construct targeting its intron/exon 2 boundary (*U6>Islet.2,* 44% mutagenesis efficacy, **Supplemental Figure 3**)(Gandhi et al., 2017) together with *Foxc>Cas9.* Papilla-specific CRISPR-based knockout of *Islet* and resulting loss of ACC cell fate was confirmed by loss of *CryBG>GFP* expression, compared to negative control individuals electroporated instead with previously published *U6>Control* sgRNA vector (Stolfi et al., 2014) targeting no sequence (**Figure 3B,C**). Therefore, we conclude that *Islet* is required for the specification and differentiation of ACCs. A smaller portion of larvae completely lost expression of the IC reporter, *C11.360>GFP,* but expression was still substantially reduced relative to the control (**Figure 3B,C**). This difference might be due to lower sensitivity of the IC reporter to *Islet* knockout, or might simply reflect the lower level of mosaicism of *C11.360>GFP* expression observed in the control.

To test whether *Islet* is required for other cell types of the papillae, we repeated papilla-specific *Islet* CRISPR knockout using our different reporters to monitor the specification or differentiation of OCs *(L141.36>GFP*) and PNs (*TGFB>GFP).* While *Islet* knockout altered the general morphology of the papillae (see further below), it did not cause any noticeable loss of OC or PN reporter expression (**Figure 3B,C**). We therefore conclude that *Islet* is required for the specification and/or differentiation of ACCs and ICs, but not OCs or PNs.

Because it was reported that an outer *Emx*+ “ring” of *Islet+* cells in each papilla co-express *Sp6/7/8* (Wagner et al., 2014), we hypothesized that *Sp6/7/8* might be required for a fate choice between ACCs and ICs. Corroborating the idea that these outer *Islet+* cells are specified as ICs, we cloned an intronic *cis*-regulatory element from the *Emx* gene that is sufficient to drive late expression specifically in ICs (**Supplemental Figure 2F**). This late ring of *Emx* expression is not to be confused with the earlier expression of *Emx* in *Foxc+/Foxg-*negative cells at the neurula stage (Liu and Satou, 2019), which represent a distinct lineage (**Figure 1**). To test the role of Sp6/7/8 in IC vs. ACC fate choice, we used the papilla-specific *Islet cis-*regulatory element to overexpress Islet or Sp6/7/8. While *Islet>Islet* did not reduce expression of either reporter, *Islet>Sp6/7/8* specifically abolished ACC reporter expression, but not that of the IC reporter (**Figure 4A,B**). In fact, IC reporter expression appeared to be slightly expanded in ∼29% of larvae electroporated with *Islet>Sp6/7/8.* Taken together, these results suggest that overexpression of Sp6/7/8 in the Islet+ cells of the papillae is sufficient to convert ACCs to an IC-like cell fate instead.

**Figure 4.**
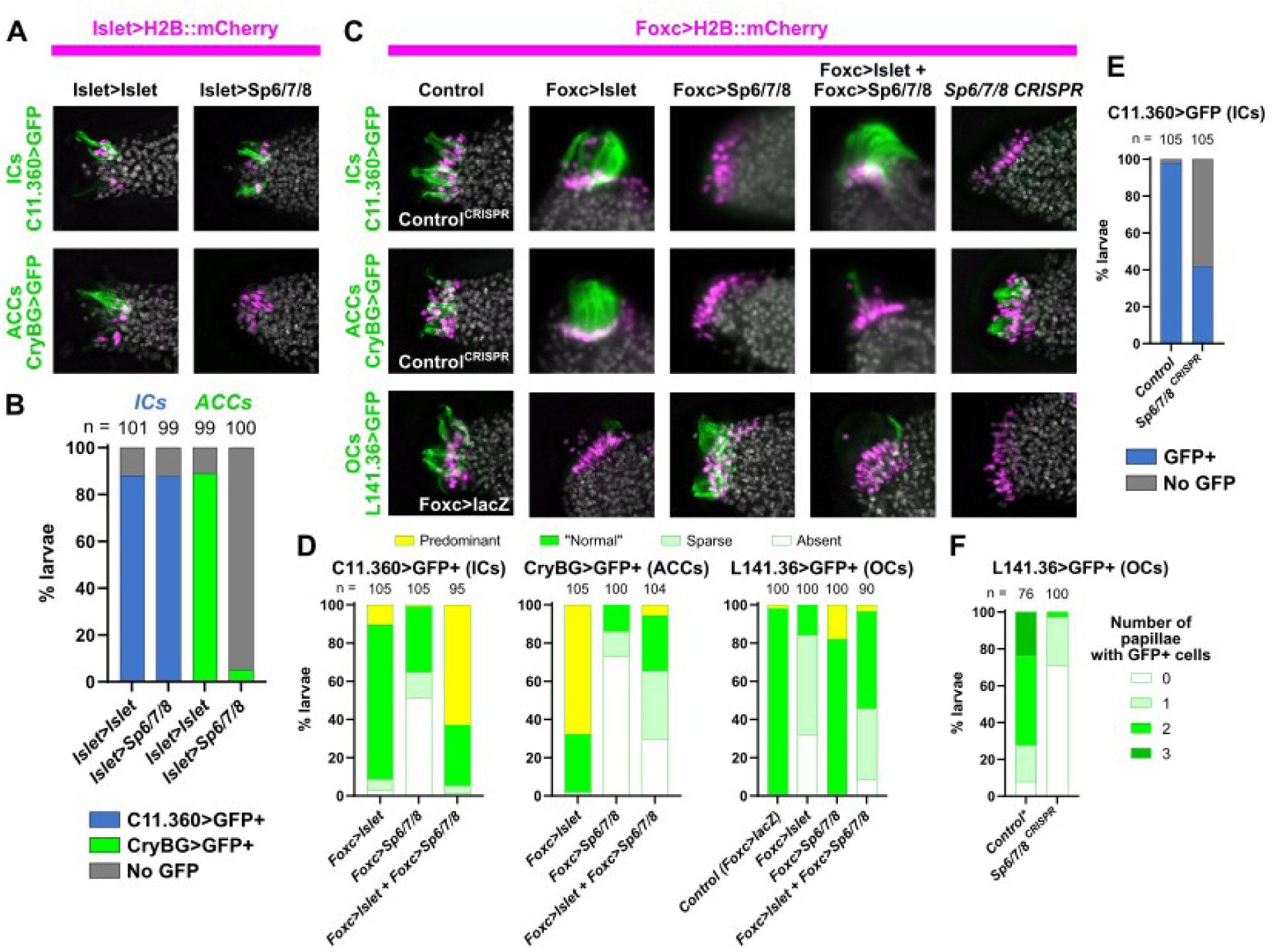
Specification of ACCs, ICs, and OCs by a combinatorial logic of Islet and Sp6/7/8. A) Overexpression of Sp6/7/8 (using the *Islet>Sp6/7/8* plasmid) in all Islet+ papilla cells results in loss of ACCs (assayed by expression of *CryBG>Unc-76::GFP*, green), but not of ICs (assayed by expression of *C11.360>Unc-76::GFP,* green). Islet overexpression (with *Islet>Islet*) does not significantly impact the specification of ACCs or ICs. Larvae at 20 hpf/20°C. B) Scoring data showing presence or absence of ICs or ACCs in the larvae represented in panel A. C) Cell type specification assayed by reporter plasmid expression (green) in larvae subjected to various *Islet* and/or *Sp6/7/8* perturbation conditions (see main text for details). For ICs and ACCs, the “control” condition is negative control CRISPR (*U6>Control*), while for OCs it is *Foxc>lacZ.* Overexpression ACC/IC sub-panels are at 18.5 hpf/20°C, all CRISPR and OC panels at 20 hpf/20°C. D) Scoring data for most larvae represented in panel C. Foxc>H2B::mCherry+ larvae were scored for cell type-specific GFP reporter expression that was “normal” (as seen in Wild Type larvae normally), “predominant” (ectopic/supernumerary GFP+ cells), “sparse” (reduced frequency/intensity of GFP expression), or “absent” (no GFP visible). E) IC reporter (*C11.360>Unc-76::GFP)* expression scored in *Foxc>H2B::mCherry+* larvae represented by the top/right-most subpanel in panel B. F) OC-specific reporter (*L141.36>Unc-76::GFP)* expression scored in *Foxc>H2B::mCherry+* larvae represented by the bottom/right-most subpanel in panel B. Scoring strategy same as in Figure 3. Asterisk denotes that the negative control was the same as in Figure 3, as the experiments were performed in parallel with the same control sample. *Foxc>Cas9* used for all CRISPR/Cas9 experiments. The *Islet* promoter used (panels A-C) was always the *Islet intron 1 + -473/-9* sequence. For overexpression conditions, *Foxc>lacZ* or *Islet>LacZ* were used to normalize the total amount of DNA introduced (see supplemental sequence file for detailed electroporation recipes).

To further show that the combination of Islet and Sp6/7/8 is sufficient to specify IC cell fate, we used the *Foxc* promoter to drive expression of Islet, Sp6/7/8, or a combination of both in the entire papilla territory. *Foxc>Islet* alone strongly promoted ACC reporter expression, as previously reported (Wagner et al., 2014), but resulted in more scattered IC reporter expression (**Figure 4C,D**). In contrast, co-electroporation of *Foxc>Islet* and *Foxc>Sp6/7/8* resulted in a large, single papilla expressing predominantly the IC reporter, not the ACC reporter (**Figure 4C,D**).

Finally, we performed papilla-specific CRISPR knockout of *Sp6/7/8,* following the same strategy for *Islet* detailed above, using new sgRNAs that we designed and validated (**Supplemental Figure 3**). Indeed, CRISPR/Cas9-mediated mutagenesis of *Sp6/7/8* in the papilla territory resulted in loss of IC cell fate, as assayed by expression of *C11.360>GFP* (**Figure 4C,E**). In contrast, the same perturbation did not diminish the expression of the ACC reporter (**Figure 4C**).

We noticed that *Foxc>Sp6/7/8* alone resulted in a large proportion of larvae lacking either ACC or IC reporter expression (**Figure 4D**). This suggested the possibility that Sp6/7/8 alone might be promoting another papilla cell fate. Indeed, we found that *Sp6/7/8* knockout by CRISPR abolishes the expression of the OC reporter (*L141.36>GFP)*, while *Foxc>Sp6/7/8* expands it slightly (**Figure 4C,D,F**). In contrast, *Foxc>Islet* alone or in combination with *Foxc>Sp6/7/8* suppressed OC reporter expression (**Figure 4C,D**), while *Islet* knockout did not affect it, as shown further above (**Figure 3B,C**). Taken together, these results suggest that a combinatorial transcriptional logic underlies papilla cell fate choices between ACCs (Islet alone), ICs (Islet + Sp6/7/8), and OCs (Sp6/7/8 alone).

### Identifying the adhesive-secreting cells of the papillae

Previous data revealed peanut agglutinin (PNA) staining as a marker for glue-secreting cell granules, the adhesive papillary cap, and adhesive prints left by larvae on the substrate (Zeng et al., 2019a; Zeng et al., 2019b). The delineation of two collocyte populations opened the question of whether both (ICs and OCs) are equally PNA-positive. To answer this question, we performed PNA stainings on larvae expressing IC or OC reporter plasmids (**Figure 5A,B**).

**Figure 5.**
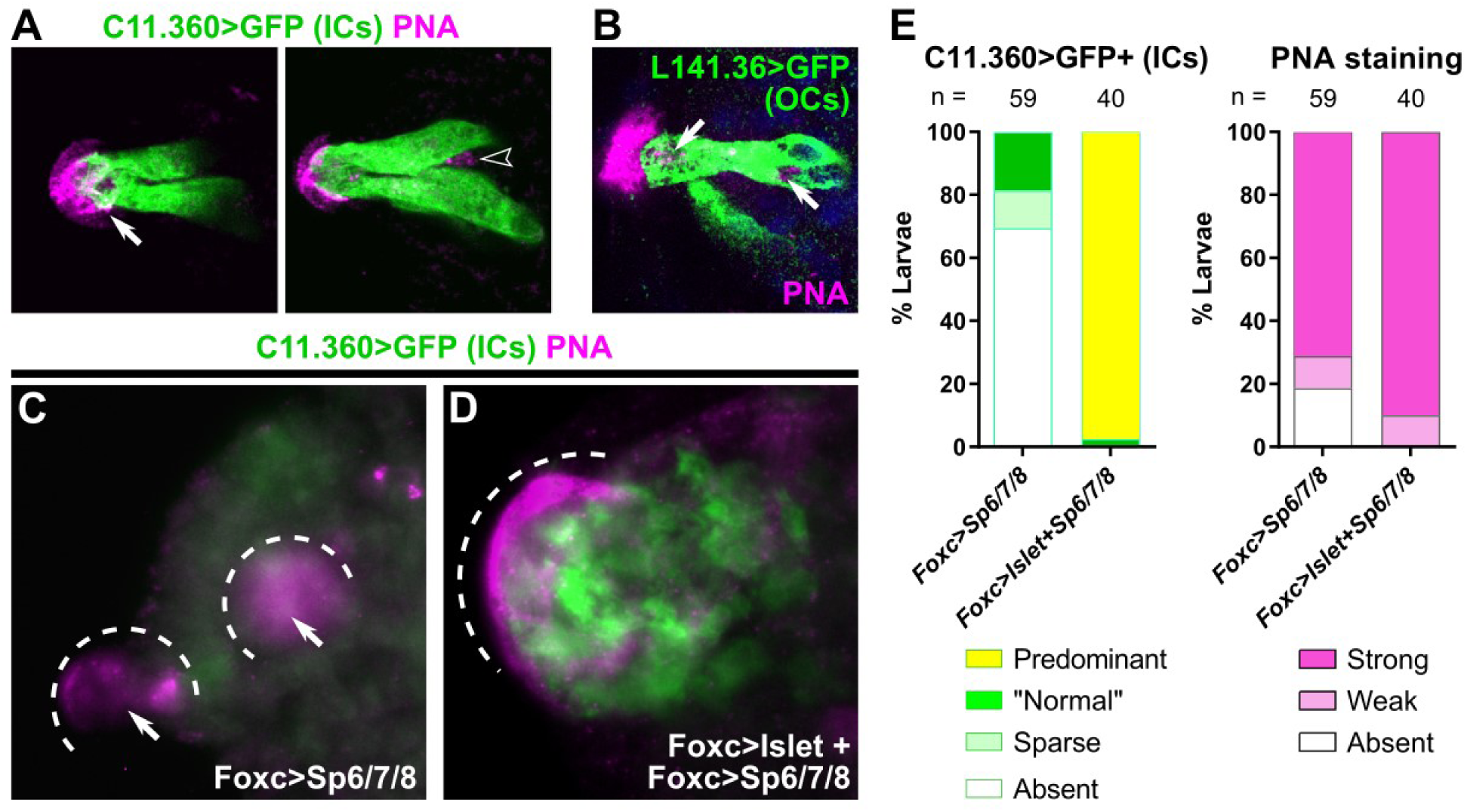
Both types of collocytes contribute to production of adhesive material. A) PNA-stained granules (pink) are seen in the hyaline cap and the apical tip of ICs (left panel, white arrow) in a *C. intestinalis* larva labeled by the *C. intestinalis C11.360>Unc-76::GFP* reporter (green). PNA-stained granules are also seen in cells not labeled by the IC reporter (right subpanel, hollow arrowhead), suggesting they are localized in a different cell type. Left and right subpanels are from different focal planes of the same papilla. B) OCs labeled with *C. robusta L141.36>Unc-76::GFP* (green) in a *C. robusta* larva, with PNA-stained granules (pink) in both apical and basal positions within the cell (white arrows). See **Supplemental Movies** for full confocal stacks. DAPI in blue. C) PNA staining (pink) in *C. robusta* upon overexpression of *Sp6/7/8* alone, showing reduction of IC specification as assayed by *C11.360>Unc-76::GFP* expression (green). PNA staining is still visible in papillae (white arrows, dashed outlines), suggesting that ICs are not the only cell type involved in adhesive glue formation. D) PNA staining (pink) and *C11.360>Unc-76::GFP* expression (green) in *C. robusta* upon overexpression of both *Islet* and *Sp6/7/8,* showing expansion of IC fate in a single large papilla (dashed outline). PNA staining is similarly expanded over the entire IC cluster, confirming that ICs produce the adhesive glue. E) Scoring larvae represented in panels C and D. PNA staining is observed despite loss of IC fate or expansion of supernumerary ICs, confirming that this cell type is one of the contributors of PNA-positive adhesive glue. *C. intestinalis* raised to 20-22 hpf at 18°C, *C. robusta* raised to 20 hpf at 20°C.

Interestingly, ICs contained PNA-stained granules only at the very apical tip (**Figure 5A**, **Supplemental Movie 1**), while the majority of PNA-stained granules were not within the ICs (**Figure 5A**). Consistently, the OCs were the main cells showing PNA-stained granules located within the papillae (**Figure 5B**, **Supplemental Movie 2**). This distribution of PNA staining corresponds to the distribution of granules previously identified by high-pressure freezing electron microscopy (Zeng et al., 2019b), in which collocytes located in the central core of the papilla contain granules mostly at their apical end. Indeed, in cross-sections, granules were most abundant inside the papillary body, likely in cells identified here as OCs.

To further demonstrate that both ICs and OCs are likely glue-secreting cells, we performed PNA staining on larvae in distinct perturbation conditions. Namely, we electroporated larvae with *Foxc>Sp6/7/8,* which was shown above to suppress IC specification, or with *Foxc>Islet* and *Foxc>Sp6/7/8* combined, which was shown to convert most of the papilla territory into ICs. Although *Foxc>Sp6/7/8* eliminated most IC reporter expression (**Figure 5C,E**), PNA staining was still present, likely due to continued presence of OCs. Similarly, although *Foxc>Islet* + *Foxc>Sp6/7/8* resulted in a single enlarged papilla with supernumerary ICs (**Figure 5D,E**), the entire papilla was often covered by PNA staining (**Figure 5D**). Taken together, these results suggest that both ICs and OCs contribute to the production of adhesive material.

### Specification of PNs and OCs from cells that downregulate Foxg

With the specification of ACCs/ICs/OCs explained in large part due to overlapping expression domains of Islet and Sp6/7/8, the precise developmental origins of the PNs and OCs still remained elusive. While it has become clear that the Islet+ cells at the core of each papilla give rise to ACCs and ICs, Papilla-specific *CRISPR* knockout of *Islet* did not abolish PNs or OCs, as shown above (**Figure 3**). This suggested they do not arise from these core Islet+ cells, consistent with their more lateral positions as shown previously by TEM (Zeng et al., 2019b). Furthermore, recently published *in situ* hybridization data showing presumptive *Pou4-*expressing PN precursors surrounding *Islet-*expressing cells at late tailbud stage (Roure et al., 2022). Indeed, co-electroporation of *Islet* reporter and PN- or OC-specific reporter plasmids clearly showed PNs and OCs immediately adjacent to, but distinct from, Islet+ cells (**Figure 2D,I**).

Might PNs and OCs be arising from the cells in between (and flanking) the three spots that downregulate *Foxg* (via repression by Sp6/7/8) and do not go on to express *Islet* (**Figure 6A**)(Liu and Satou, 2019; Roure et al., 2022; Wagner et al., 2014)? To test this, we used the MEK (MAPK kinase) inhibitor U0126 to expand *Islet* expression as previously done (**Figure 6A**)(Wagner et al., 2014). While treatment with 10 µM U0126 at 7.5 hpf (between neurula and early tailbud stages) predictably expanded *Islet* reporter expression, it also eliminated expression of the PN reporter *C4.78>GFP*, as well as that of the OC reporter *L141.36>GFP* (**Figure 6B,C**). These results suggest that *Foxg+* papilla cells that maintain *Foxg* expression go on to express *Islet* and give rise to ACCs and ICs, while the cells that activate *Sp6/7/8* and downregulate *Foxg* in response to MAPK signaling go on to give rise to OCs and PNs instead.

**Figure 6.**
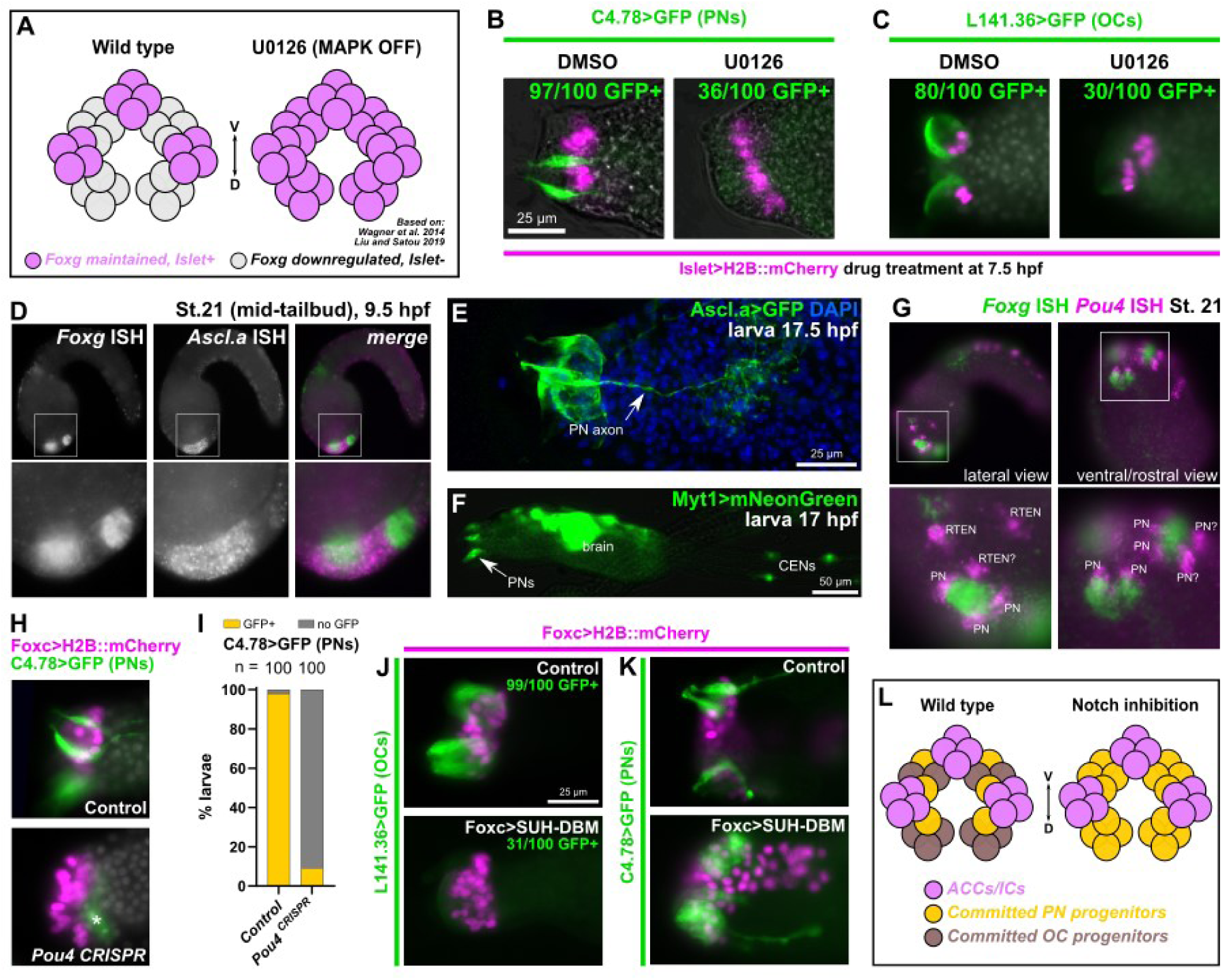
Specification of PNs and OCs from *Islet-*negative cells by MAPK and Notch pathways. A) Diagram showing effect of MAPK inhibition with the pharmacological MEK inhibitor U0126, based on findings from Wagner et al. 2014 and Liu and Satou 2019. Inhibition of FGF/MAPK results in expansion of *Foxg* and *Islet* from three discrete foci to a large “U-shaped” swath, transforming three papillae into a single, enlarged papilla (similar results reported with BMP inhibition by Roure et al. 2022). B) U0126 treatment at 7.5 hpf/20°C results in loss of PNs (assayed by *C4.78>Unc-76::GFP,* green) upon expansion of Islet+ cells (pink nuclei), relative to DMSO alone. C) The same treatment results in loss of OCs (*L141.36>Unc-76::GFP,* green) upon expansion of Islet+ cells (pink nuclei). All larvae in B,C at 17 hpf/20°C. D) Two-color, whole-mount mRNA *in situ* hybridization for *Foxg* (green in merged image) and *Ascl.a* (*KH. L9.13,* pink). E) Larva electroporated with *Ascl.a>Unc-76::GFP* labeling several papilla cells including PNs. F) *Myt1>*mNeonGreen labeling PNs and other neurons including Caudal Epidermal Neurons (CENs). G) Two-color *in situ* hybridization of *Foxg* (green) and *Pou4* (pink), the latter labeling adjacent PNs and possibly Rostral Trunk Epidermal Neurons (RTENs). H) Lineage-specific CRISPR/Cas9-mediated mutagenesis of *Pou4* results in loss of PN reporter expression *(C4.78>Unc-76::GFP,* green). Larvae at 17 hpf/20°C. I) Scoring of *Foxc>H2B::mCherry+* larvae represented in panel H. J) Inhibition of Delta/Notch signaling using *Foxc>SUH-DBM* results in reduced expression of OC reporter (*L141.36>Unc-76::GFP,* green) at 21 hpf/20°C and K) concomitant expansion of supernumerary PNs at 17 hpf/20°C (labeled by *C4.78>Unc-76::GFP,* green) relative to *Foxc>lacZ* control. L) Summary diagram and model of effects of Delta/Notch inhibition on PN/OC fate choice in *Islet-*negative (but formerly *Foxg+*) papilla progenitor cells. All *Islet* reporters are the *Islet intron 1 + bpFOG>H2B::mCherry*.

### PNs are specified by common peripheral neuron regulators

Previous papilla-specific TALEN knockout of the neuronal transcription factor-encoding gene *Pou4* successfully eliminated PNs and the larva’s tail resorption response to mechanical stimuli (Sakamoto et al., 2022). Pou4 has been previously implicated in a Myt1-dependent regulatory cascade that specifies the caudal epidermal neurons (CENs) of the tail, from neurogenic midline cells expressing the proneural bHLH transcription factor *Ascl.a (KH.L9.13,* sometimes called *Ascl2* or *Ascl.b* previously)(Pasini et al., 2006; Roure et al., 2022; Roure and Darras, 2016; Tang et al., 2013; Waki et al., 2015). To precisely visualize the neurogenic cells of the papillae, we performed double (two-color) mRNA *in situ* hybridization for *Ascl.a* and *Foxg* at the mid-tailbud stage. Indeed, *Ascl.a* expression was seen broadly in the papilla territory surrounding the three *Foxg+* cell clusters (**Figure 6D**). This was confirmed by an *Ascl.a* fluorescent protein reporter plasmid that labeled a broad set of papilla territory cells, including PNs and their axons (**Figure 6E**). Furthermore, a previously published *Myt1* reporter (Tolkin and Christiaen, 2016) was also found to be expressed in the PNs (**Figure 6F**). Double *in situ* of *Pou4* and *Foxg* revealed *Pou4+* cells surrounding each *Foxg+* cluster, corroborating a recent report (Roure et al., 2022)(**Figure 6G**). It was not immediately clear which Pou4+ cells were PN precursors and which were nearby Rostral Trunk Epidermal Neuron (RTEN) precursors. Based on our images and those of the most recent study (Roure et al., 2022), we propose that there are initially two Pou4+ cells per papilla, later dividing to give rise to the four PNs per papilla as previously described (Zeng et al., 2019b). This would mirror the development of the epidermal neurons of the tail, in which neurons are born side-by-side as pairs after a final cell division by a committed mother cell (Pasini et al., 2006). Papilla-specific CRISPR knockout of *Pou4* with new sgRNAs (**Supplemental Figure 3**) recapitulated the loss of PN differentiation by the previously published TALEN knockout (Sakamoto et al., 2022), as assayed by *C4.78* reporter expression (**Figure 6H,I**). In contrast, *Pou4* knockout had no effect on the specification of ACCs or OCs, suggesting Pou4 function is specific for PN fate in the papillae (**Supplemental Figure 4**). Taken together, these results suggest that PNs are specified from interspersed neurogenic progenitors that are carved out by MAPK signaling through Sp6/7/8-dependent repression of *Foxg/Islet*.

### Notch signaling regulates the fate choice between PNs and OCs

Because both OCs and PNs appeared to arise from *Foxg-*downregulating, *Islet*-negative cells, we sought to test whether an additional regulatory step is required for the fate choice between these two cell types. In the neurogenic midline territory of the tail epidermis, lateral inhibition by Delta/Notch signaling regulates the final number and spacing of CENs (Chen et al., 2011; Pasini et al., 2006; Tang et al., 2013). Delta/Notch limits the expression of Myt1, which in turn activates *Pou4* expression. We therefore decided to test whether a similar mechanism in controlling the number of PNs and OCs surrounding each papilla. To test the requirement of Delta/Notch, we overexpressed a DNA-binding mutant of the Notch co-factor RBPJ/SUH (SUH-DBM)(Hudson and Yasuo, 2006). Indeed, electroporation with *Foxc>SUH-DBM* resulted in loss of OC reporter expression (**Figure 6J**), and concomitant expansion of PN reporter expression (**Figure 6K**). We conclude that Delta/Notch signaling regulates PN vs. OC fate choice in neurogenic progenitor cells surrounding each presumptive papilla, with Notch delimiting the specification of supernumerary neurons, thus allowing OCs to form (**Figure 6L**). This common origin of PNs and OCs is also supported by the recent finding that the latter appear to have basal bodies like the PNs, but without the accompanying sensory cilia (Zeng et al., 2019b). Interestingly, papilla-specific knockout of *Foxg* resulted in moderate loss of PN reporter expression (*TGFB>GFP)*, and very little effect on the OC reporter (**Supplemental Figure 4**). This suggests differing requirement for Foxg in different cell type-specific branches of the papilla regulatory network, despite all these cell types arising from cells that initially express Foxg.

### Regulation of papilla morphogenesis by Islet

It was previously shown that *Foxg* or *Islet* overexpression induces the formation of a single enlarged “megapapilla”, in which all cells are substantially elongated relative to the rest of the epidermis (Liu and Satou, 2019; Wagner et al., 2014). We have shown above that this appears to be driven by expansion of ACCs and/or ICs, which are atypically elongated in the apical-basal direction and form apical protrusions and microvilli. Islet is sufficient for apical-basal elongation of epidermal cells (Wagner et al., 2014), and morpholino-knockdown of *Foxg* (which is upstream of *Islet)* also impairs proper papilla morphogenesis (Liu and Satou, 2019). We asked if *Islet* is required for papilla morphogenesis, using papilla-specific CRISPR knockout of *Islet.* Knocking out *Islet* in the papilla territory impaired the formation of the typically “pointy-shaped” papillae, resulting instead in blunt cells with flat, broader apical surfaces (**Figure 7A,B**). This result suggested that transcriptional targets downstream of *Islet* might be regulating the distinct cell shape of ACCs/ICs.

**Figure 7.**
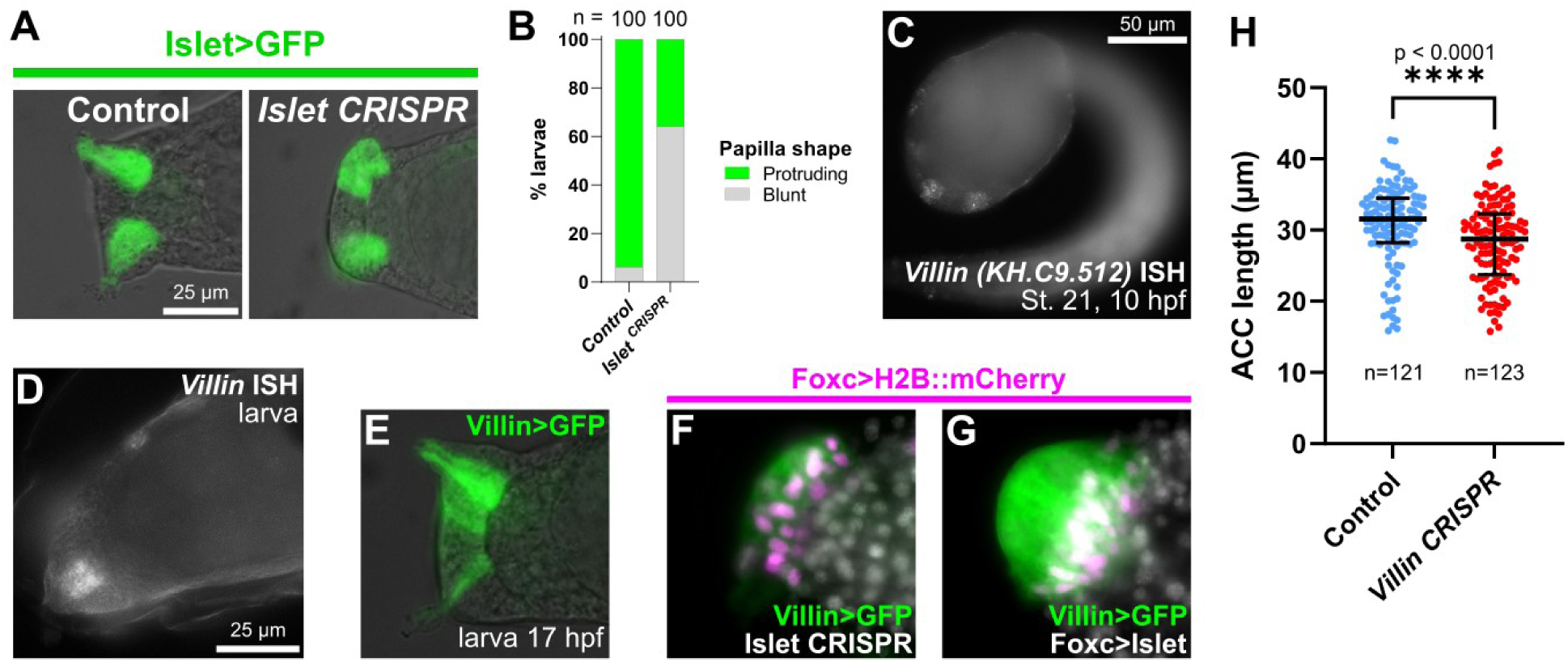
Islet is also required for papilla morphogenesis and regulates expression of the cytoskeletal effector Villin. A) Papilla shape is blunt at the apical end upon tissue-specific CRISPR/Cas9-mediated mutagenesis of *Islet.* Embryos were electroporated with *Islet intron 1 + -473/-9>Unc-76::GFP* and *Foxc>Cas9. Islet* CRISPR was performed using *U6>Islet.2* sgRNA plasmid, and the negative control used *U6>Control.* Larvae were imaged at 20 hpf/20°C. B) Scoring of percentage of GFP+ larvae classified as having normal “protruding” or blunt papillae, as represented in panel A. C) *In situ* mRNA hybridization of *Villin,* showing expression in *Foxg+/Islet+* central papilla cells at 10 hpf/20°C (stage 21) and D) at the larva stage. E) *Villin - 721/-1>Unc-76::GFP* showing expression in the papilla territory, strongest in the central cells. F) *Villin -721/-1>Unc-76::GFP* is downregulated by tissue-specific CRISPR/Cas9 mutagenesis of *Islet* (*Foxc>Cas9 + U6>Islet.1 + U6>Islet.2,* see text for details) and G) upregulated by overexpressing Islet (*Foxc>Islet,* see text for details). F and G panels both at 17 hpf/20°C. H) Quantification of ACC lengths measured in negative control and papilla-specific *Villin* CRISPR larvae at 17 hpf/20°C. Large bars indicate medians, smaller bars indicate interquartile ranges. P-value denotes one-tailed Mann-Whitney test. Raw measurements available in **Supplemental Table 3.**

To identify potential candidate effectors of morphogenesis downstream of *Islet*, we used bulk RNAseq to measure differential gene expression between different Islet perturbation conditions (**Figure 7C**). We compared “negative control” embryos to (1) embryos in which *Islet* was overexpressed in the whole territory using the *Foxc* promoter (*Foxc>Islet)*, and (2) embryos in which *Islet* was knocked out specifically in the papilla lineage by CRISPR/Cas9. For this, we designed an additional sgRNA targeting the first exon of *Islet,* to be used in combination with the already published sgRNA to generate larger deletions. This new sgRNA vector, which we named *U6>Islet.1,* resulted in a mutagenesis efficacy of 20%. (**Supplemental Figure 3**).

Whole embryos from each condition were collected at 12 hpf (*Islet* conditions) at 20°C in biological triplicate. RNA was extracted from pooled embryos in each sample, and RNAseq libraries were prepared from poly(A)-selected RNAs and sequenced by Illumina NovaSeq. This bulk RNAseq approach revealed that Islet overexpression results in the upregulation of several ACC markers from previous scRNAseq analysis (**Supplemental Table 2**)(Sharma et al., 2019). With Islet overexpression, this included ACC markers previously validated by mRNA *in situ* hybridization or reporter gene expression, such as *CryBG* and *Atp2a (KH.L116.40).* Many ACC markers were conspicuously absent, but this may be due to the relatively early timepoint (12 hpf, late tailbud stage), well before hatching and ACC differentiation. This was a deliberate choice, as we were focused on papilla morphogenesis, which begins around this stage (Wagner et al., 2014). One resulting candidate *Islet* target revealed by RNAseq was *Astl-related (KH.C9.850),* and its expression in the Islet+ cells of the papillae was confirmed by *in situ* hybridization (**Supplemental Figure 5A**). Indeed, *Islet* knockout by CRISPR eliminated *Astl-related* reporter expression, supporting our approach to identifying new targets of Islet (**Supplemental Figure 5B,C**)

One particularly interesting ACC-specific candidate that was amongst the genes most highly upregulated by Islet overexpression was *Villin (KH.C9.512),* an ortholog of the *Villin* family of genes encoding effectors of actin regulators (Khurana and George, 2008). We confirmed the expression of *Villin* in the papillae by *in situ* hybridization and reporter plasmids (**Figure 7C-E**). In the *Islet* CRISPR condition, *Villin* was the top downregulated gene by *Islet* CRISPR knockout as well. *Villin* reporter expression was reduced in intensity but not completely lost upon knockout of *Islet* by CRISPR (68/100 electroporated larvae were still GFP+, vs. 97/100 in the negative control, **Figure 7H**), yet was also dramatically upregulated by Islet overexpression (94/100 larvae, **Figure 7I**). This suggests partially redundant activation of *Villin* by another factor, likely at earlier developmental stages (e.g. by *Foxc* or *Foxg*), and that Islet might be required for its sustained expression specifically in the central cells of the papilla throughout morphogenesis. This is consistent with the weak but broad expression of *Villin>GFP* in the entire papilla territory (**Figure 7E**).

To show that *Villin* is required for proper morphogenesis of *Islet+* cells in the papilla, we performed tissue-specific CRISPR knockout using a combination of three validated sgRNAs spanning most of the coding sequence (**Supplemental Figure 3E**). Because the functionally important “headpiece” domain is encoded by the last exon, we combined an sgRNA targeting this exon with two sgRNAs targeting more upstream exons. In *Villin* CRISPR larvae, ACCs were slightly but significantly shorter in length along the apical-basal axis (**Figure 7F, Supplemental Table 3**). Taken together, these results suggest that Islet is required for proper papilla morphogenesis, possibly through its ability to activate the expression of cytoskeletal effector genes such as *Villin* during this process.

### An investigation into the cell and molecular basis of larval settlement metamorphosis

With our different CRISPR knockouts affecting different cell types of the papillae, we asked how these different perturbations might affect larval metamorphosis. Only the involvement of the PNs in triggering metamorphosis has been demonstrated (Sakamoto et al., 2022; Wakai et al., 2021), but it is not yet known how the regulatory networks and cell types of the papillae affect different processes during metamorphosis. We performed papilla-specific CRISPR as above using the *Foxc>Cas9* vector, targeting the four different transcription factors we have shown to be involved in patterning the cell types of the papillae: *Pou4, Islet, Foxg,* and *Sp6/7/8.* We assayed tail retraction and body rotation at the last stage of metamorphosis (Hotta et al., 2020) (**Figure 8A,B**), as these are two processes that can be uncoupled in certain genetic perturbations or naturally occurring mutants (Nakayama-Ishimura et al., 2009).

**Figure 8.**
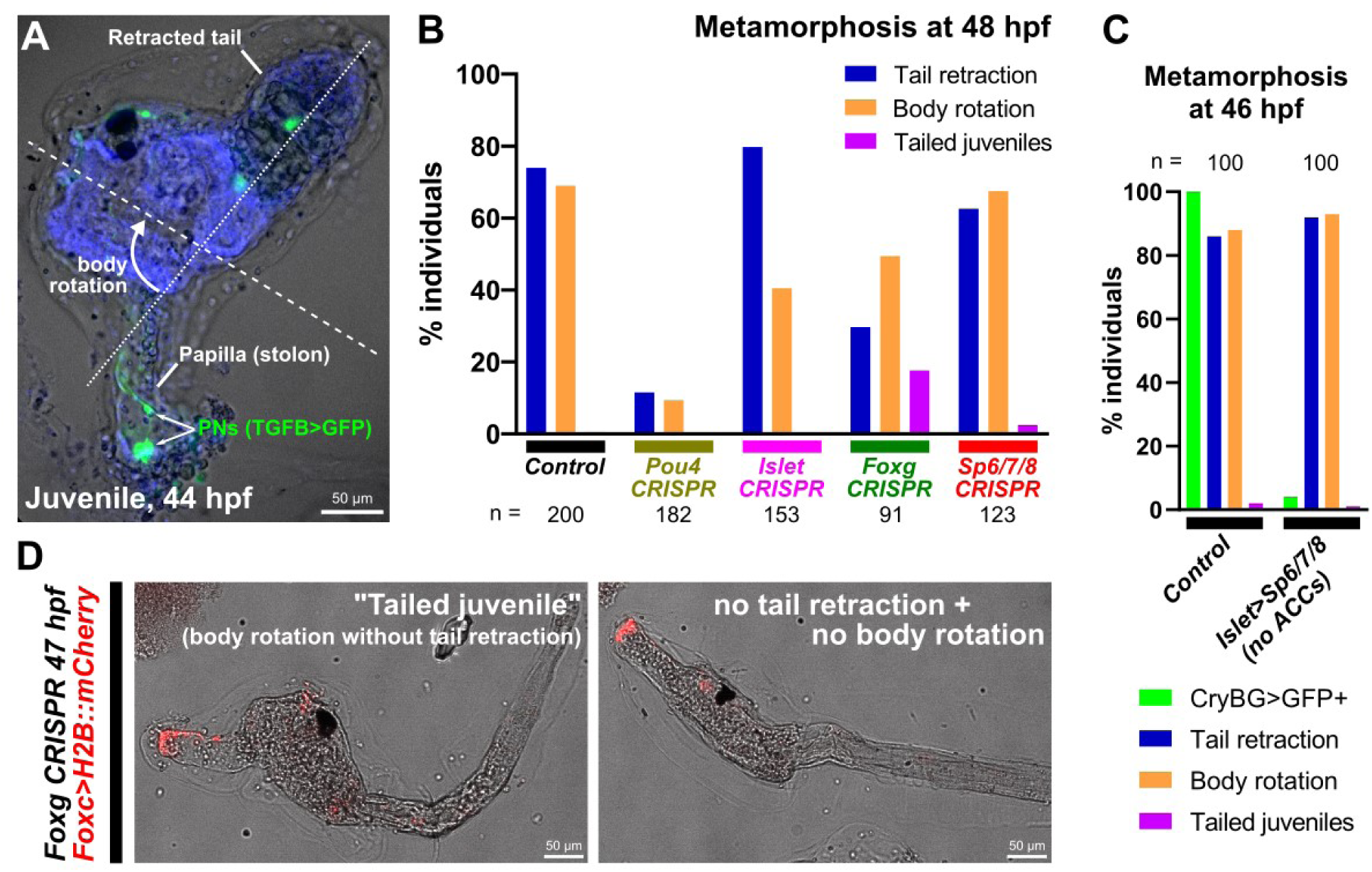
Genetic perturbations of metamorphosis. A) *Ciona robusta* juvenile undergoing metamorphosis, showing the retracted tail and rotated anterior-posterior body axis (dashed lines). Papilla Neurons (PNs) in the former papilla (now substrate attachment stolon, or holdfast) labeled by *TGFB>Unc-76::GFP* (green). Animal counterstained with DAPI (blue). B) Scoring of total individuals showing tail retraction and/or body rotation at 48 hpf/20°C in various papilla territory-specific (using *Foxc>Cas9*) CRISPR-based gene knockouts. “Tailed juveniles” have undergone body rotation but not tail retraction, whereas normally body rotation follows tail retraction. The sgRNA plasmids used for each condition were as follows-Control: *U6>Control; Pou4: U6>Pou4.3.21 + U6>Pou4.4.106; Islet: U6>Islet.2; Foxg: U6>Foxg.1.116 + U6>Foxg.5.419; Sp6/7/8: U6>Sp6/7/8.4.29 + U6>Sp6/7/8.8.117.* C) Plot showing lack of any discernable metamorphosis defect after eliminating ACCs using *Islet intron 1 + bpFOG>Sp6/7/8* (images not shown). Only *Islet intron 1 + bpFOG>H2B::mCherry+* individuals were scored. ACC specification was scored using the *CryBG>Unc-76::GFP* reporter. D) Example of “tailed juveniles” at 47 hpf/20°C compared to a larva in which no tail retraction or body rotation has occurred, elicited by tissue-specific *Foxg* CRISPR (*Foxc>Cas9 + U6>Foxg.1.116 + U6>Foxg5.419).* See Supplemental Figure 6 for scoring.

Knockout of *Pou4* recapitulated recent published results on this transcription factor (Sakamoto et al., 2022). Namely, both tail retraction and body rotation were blocked in the vast majority of individuals This suggests that proper specification and/or differentiation of PNs by *Pou4* is crucial for the ability of the larva to trigger the onset of metamorphosis. In contrast, *Islet* knockout did not affect tail retraction, but body rotation appeared somewhat impaired. This suggested that ACCs/ICs are not required for tail retraction, but might play a role in regulating body rotation downstream of it. Eliminating ACCs using *Islet>Sp6/7/8* had no effect on either tail retraction or body rotation (**Figure 8C**), confirming that ACCs are not required for metamorphosis, but that perhaps certain Islet targets might specifically regulate body rotation. Unsurprisingly, *Foxg* knockout impaired both tail retraction and body rotation, but also resulted in a noticeable fraction (∼17%) of “tailed juveniles” in which body rotation begins even in the absence of tail retraction (**Figure 8B,D**). This unusual effect was seen even when repeating the experiment independently, revealing consistent uncoupling of these two processes upon *Foxg* knockout (**Supplemental Figure 6**). Finally, *Sp6/7/8* CRISPR did not substantially alter either tail retraction nor body rotation. Taken together, these results paint a more complex picture of regulation of metamorphosis by the papillae. Our findings suggest that different cell types of the papillae might play distinct roles in the regulation of metamorphosis, perhaps interacting with one another to regulate different steps, or that certain transcription factors might be required for the expression of key rate-limiting components of these different processes. Further work will be required to disentangle these different cellular and genetic factors, which we hope will be aided by our cell type-specific reporters and CRISPR reagents.

## Discussion

Sensory systems are crucial for interactions between organisms and their environment. The concentration of sensory functions in the head is thought to have played a central role in vertebrate evolution, leading to a more active behavior emerging from early filter-feeding chordate ancestors (Diogo et al., 2015; Gans and Northcutt, 1983; Patthey et al., 2014). The peripheral components of the sensory systems in vertebrates arise from two physically close but distinct ectodermal cell populations, the cranial sensory placodes and the neural crest (Martik and Bronner, 2021). Cranial sensory placodes are characterized by their common ontogenetic origin from a crescent-shaped region surrounding the anterior neural plate. Our understanding of the evolutionary origins of structures long presented as vertebrate novelties has benefited from an increasing number of comparative studies with tunicates. Several discrete populations of peripheral sensory cells originating from distinct ectodermal regions in tunicates have respectively been linked to neural crest and cranial placodes, among them the sensory adhesive papillae (Abitua et al., 2015; Abitua et al., 2012; Horie et al., 2018; Papadogiannis et al., 2022).

Our results have confirmed the existence of molecularly distinct cell types in the *Ciona* papillae, and the developmental pathways that specify them (summarized in **Figure 9**). Using CRISPR/Cas9-mediated mutagenesis, we have shown that different transcription factors are required for their specification, differentiation, and morphogenesis. Namely, ACCs and ICs are specified from Foxg+/Islet+ cells at the center of each of the three papillae, while OCs and PNs are specified from interleaved Islet-negative cells that nonetheless derive from initially Foxg+ cells. While Sp6/7/8 specifies IC vs. ACC fate among Islet+ cells, Delta/Notch signaling suppresses PN fate and promotes OC fate among Islet-negative cells. While there appear to be two molecularly distinct collocyte subtypes (OCs and ICs), both contain granules that are stained by PNA, and therefore both are likely to be involved in glue production. Where they differ might be in the timing of glue production and/or secretion, as they showed distinct subcellular localization of PNA+ granules, and PNA production was previously shown to start very early (Zeng et al., 2019b).

**Figure 9.**
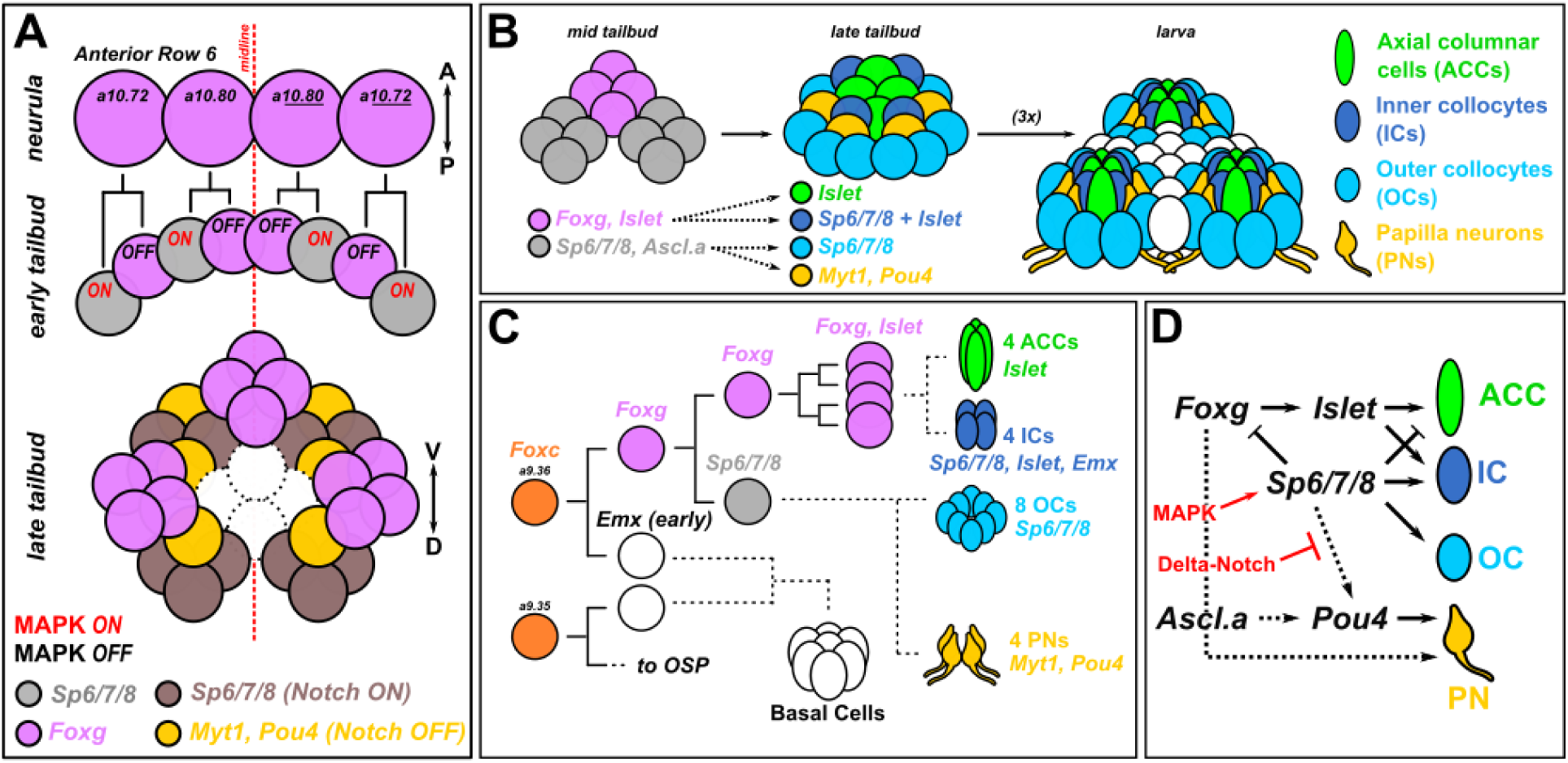
Summary diagram. A) Updated diagram of the development of the anterior descendants of Row 6 in the neural plate to show the proposed patterns of MAPK and Delta/Notch signaling that set up the three *Foxg+* clusters and interleaved *Foxg-*negative neurogenic cells. B) Diagram proposing the contributions of *Foxg+* and *Foxg-*negative cells to later patterns of transcription factors that specify the different cell types found in each papilla, which is in turn is repeated three times, thanks to the process shown in panel A. C) Papilla development shown as cell lineages, with dashed lines indicating uncertain cell divisions and lineage history. Cell type numbers based on Zeng et al. 2019b. D) Provisional gene regulatory network diagram of the signaling pathways and transcription factors involved in specification and differentiation of the different papilla cell types. Arrowheads indicate activating gene expression or promoting cell fate, while blunt ends indicate repression of gene expression of cell fate. Solid lines indicate regulatory links (direct or indirect) that are supported by the current data and literature. Dashed lines indicate regulatory links that have not been tested, or need to be investigated in more detaill. OSP: oral siphon primordium. A-P: anterior-posterior. D-V: dorsal-ventral.

Our results also demonstrate a clear distinction between CryBG+ ACCs and Pou4+ PNs. Previously these cells types have been confused and only recently distinguished by TEM and different molecular markers (Zeng et al., 2019b). Here we show that, while both arise from Foxc+/Foxg+ cells, ACCs are not specified by Pou4, and PNs are not specified by *Islet*. However, because *Pou4* can activate *Foxg* expression in a proposed feedback loop (Chacha et al., 2022), overexpression of *Pou4* might result in ectopic activation of ACC markers via ectopic *Foxg* and *Islet* activation.

There are still unanswered questions that we hope future work will address:

1. *How do the three “spots” of Foxg+/Islet+ cells form in an invariant manner?* Ephrin-Eph signaling is often responsible for suppression of FGF/MAPK signaling in alternating cells in *Ciona* embryos, via asymmetric inheritence/activation of p120 RasGAP (Haupaix et al., 2014; Haupaix et al., 2013). This is also true in the earlier patterning of the papilla territory, where EphrinA.d suppresses FGF/MAPK to promote *Foxg* activation (Liu and Satou, 2019). Curiously, later expression of *EphrinA.d* in the lineage appears to be stronger in medial *Foxg+* cells than in lateral cells (Liu and Satou, 2019). This distribution would suffice to result in the alternating ON/OFF pattern of MAPK activation at the tailbud stage that results in the three foci of *Foxg/Islet* expression. Thus it may be informative to test the ongoing functions of Ephrin-Eph signaling in this lineage throughout development.
2. *How are PNs specified adjacent to the Islet+ cells?* Since Delta/Notch signaling is involved in PN vs. OC fate, we propose that there is something that biases Notch signaling to be activated preferentially in those cells not touching the *Islet+* cells. This could be due to cell-autonomous activation of Notch signaling in the *Islet+* cells, which in turn would allow for suppression of Notch in adjacent cells fated to become PNs. A recent study showed *Pou4* expansion with concomitant “U”-shaped expansion of *Islet* when inhibiting BMP signaling (Roure et al., 2022), while our results with the MEK inhibitor U0126 (based on experiments from Wagner et al. 2011) suggests the opposite, the elimination of *Pou4+* PNs. Why the discrepancy? One possibility is that expansion of *Islet* (with or without BMP inhibition) results in specification of supernumerary RTEN-like neurons from adjacent epidermis, not PNs. While *Pou4* (and other markers, like *VGluT)* are expressed in all epidermal neurons, our preferred PN marker *KH.C4.78* is not expressed in RTENs.
3. *What activates the expression of Sp6/7/8 in Islet+ cells, ultimately promoting IC specification?* We do not yet know the exact mitotic history of the ACCs/ICs. How do the initially four Islet+ cells divide, and which daughter cells give rise to ACCs vs. ICs? Are ACCs/ICs specified in an invariant manner, or is there some variability? Finally, what allows the “creeping” activation of *Sp6/7/8* in the outer ring of cells that likely become the ICs? Is this due to additional asymmetric FGF/MAPK activation downstream of Ephrin-Eph? Or could this be due to some other signaling pathway? Is there an inductive signal from adjacent cells, for instance PNs or common PN/OC progenitors?
4. *How do the different papilla cell types regulate metamorphosis?* We noticed some uncoupling of tail resorption and body rotation upon targeting different transcription factors for deletion in the papillae (**Figure 8**). This was most apparent in the *Foxg* knockout, in which a substantial portion of individuals displayed the “tailed juvenile” phenotype in which body rotation proceeds even in the absence of tail resorption. From the *Pou4* knockout, it is clear that PNs are upstream of both tail resorption and body rotation, but the partial uncoupling seen with the other manipulations were particularly intriguing. This uncoupling has been reported before in *Cellulose synthase* mutants, which results in similar tailed juveniles (Sasakura et al., 2005). Additionally, perturbation of Gonadotropin-releasing hormone (GnRH) or the prohormone convertase enzyme (PC2) necessary for its processing similarly blocks tail resorption but not body rotation and further adult organ growth (Hozumi et al., 2020). Thus, it is possible that while Pou4 disrupts PN specification altogether, Foxg might be more specifically required for GnRH or other neuropeptide expression/processing in the PNs. Supporting this idea, the *Foxg* CRISPR did not disrupt PN specification (as assayed by *TGFB>GFP* reporter expression) as robustly as did *Pou4* CRISPR. Finally, the appearance of juveniles with resorbed tails but no further body rotation in the *Islet* CRISPR condition suggests a crucial role for ACCs or ICs in metamorphosis downstream of PNs. Clearly, more work will be required to understand the contributions of different cell types, and potentially different molecular pathways in the same cell type, towards either activation or suppression of specific body plan rearrangement processes in tunicate larval metamorphosis.
5. *Are the tunicate larval papillae homologous to vertebrate cement glands?* The papillae have often been compared to the cement glands of fish and amphibian larvae, which are transient adhesive organs secreting sticky mucus (Rétaux and Pottin, 2011). Even though they are innervated by trigeminal fibers, the secreting cells from the cement gland differentiate from a surface ectoderm region anterior to the oral ectoderm and the panplacodal domain (Pottin et al., 2010). Therefore, they are usually not considered placodal derivatives. Despite their variability in size, number, and location, head adhesive organs are proposed to be homologous across vertebrate species based on their shared expression of Pitx1/2 and BMP4 genes, innervation by trigeminal fibers, and inhibiting mechanism of swimming behavior (Pottin et al., 2010; Rétaux and Pottin, 2011). While recent papers have revealed an important role for BMP signaling in pattering the tunicate papilla territory (Liu et al., 2023; Roure et al., 2022), additional work on the molecular basis of the papillary glue in tunicates will be required to answer questions of homology between these adhesive organs. Our identification of molecular signatures for both collocyte subtypes in the papillae of *Ciona* provides a starting point for future investigations.

## Supporting information

Supplemental Sequences File

## Acknowledgments

We thank members of the labs at Georgia Tech and Innsbruck, for critical feedback and support. We thank Susanne Gibboney, Tanner Shearer, Alex Gurgis, Lindsey Cohen, and Akhil Kulkarni for technical assistance. This work was funded by NSF grant 1940743 and NIH grants GM143326 and HD104825 to AS; an NSF graduate fellowship to CJJ; and by FWF grant P 35402-B to UR.

---

**Supplemental Figure 1.**
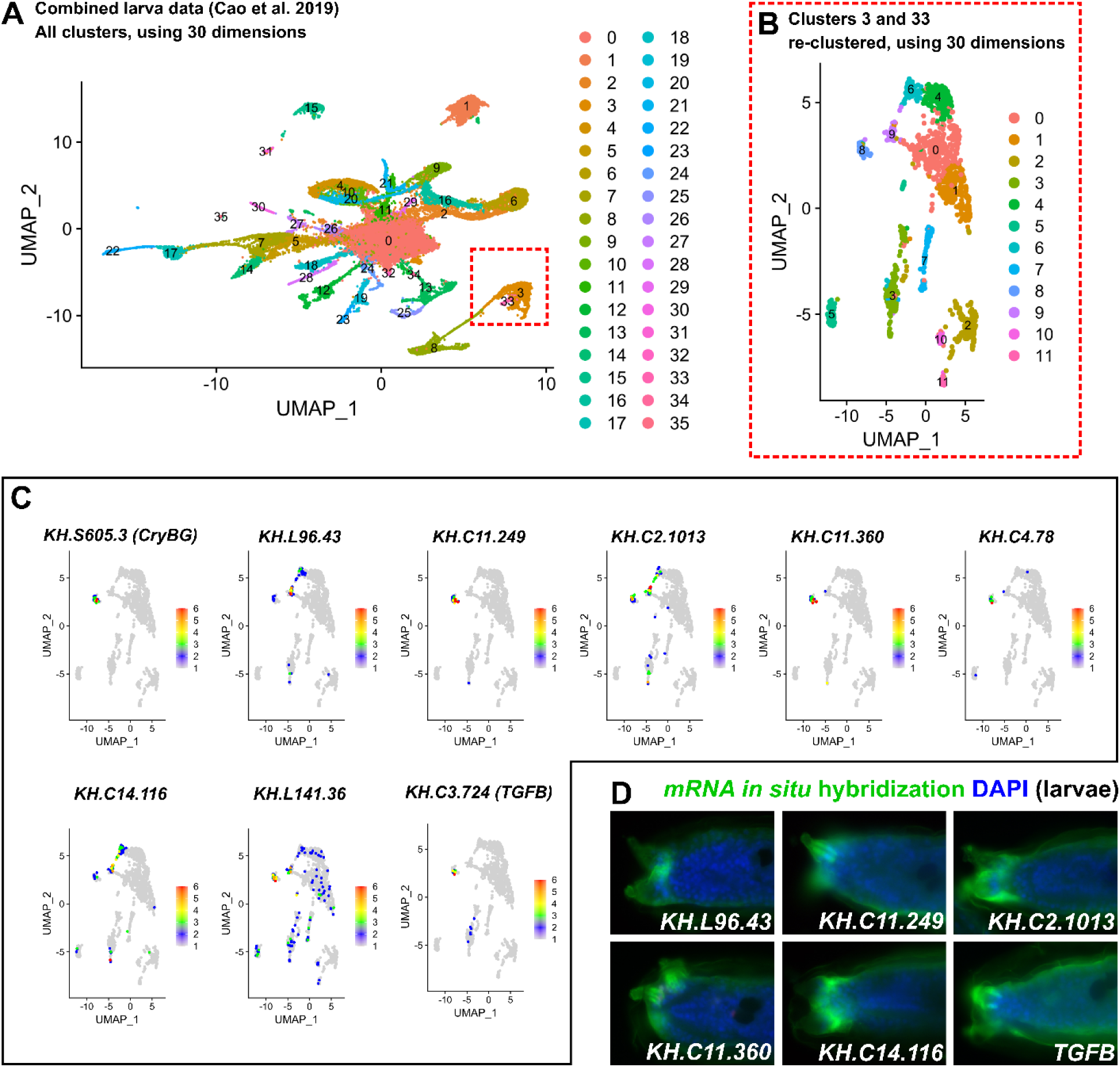
Finding papilla cell type-specific markers in single-cell RNAseq data. A) Cell clusters based from reanalysis and re-clustering of whole-larva single-cell RNA sequencing (scRNAseq) data from Cao et al. 2019. Dashed red box indicated clusters 3 and 33, which appeared to correspond to several papilla cell types. B) Cells from clusters 3 and 33 from plot A set aside and re-clustered. C) Differential expression plots showing examples of candidate papilla cell type marker genes mapped onto clusters in B. D) Fluorescent, whole-mount *in situ* mRNA hybridization (green) for certain genes plotted in C, labeling different cells in the papillae of *Ciona robusta* (*intestinalis* Type A) hatched larvae. Unless specifically named, genes are indicated by KyotoHoya (KH) ID numbers (e.g. *KH.L96.43*). All larvae were fixed at 18 hours post-fertilization (hpf), 20°C, except for *C11.360* and *C2.1013* (18.5 hpf). Blue counterstain is DAPI.

**Supplemental Figure 2.**
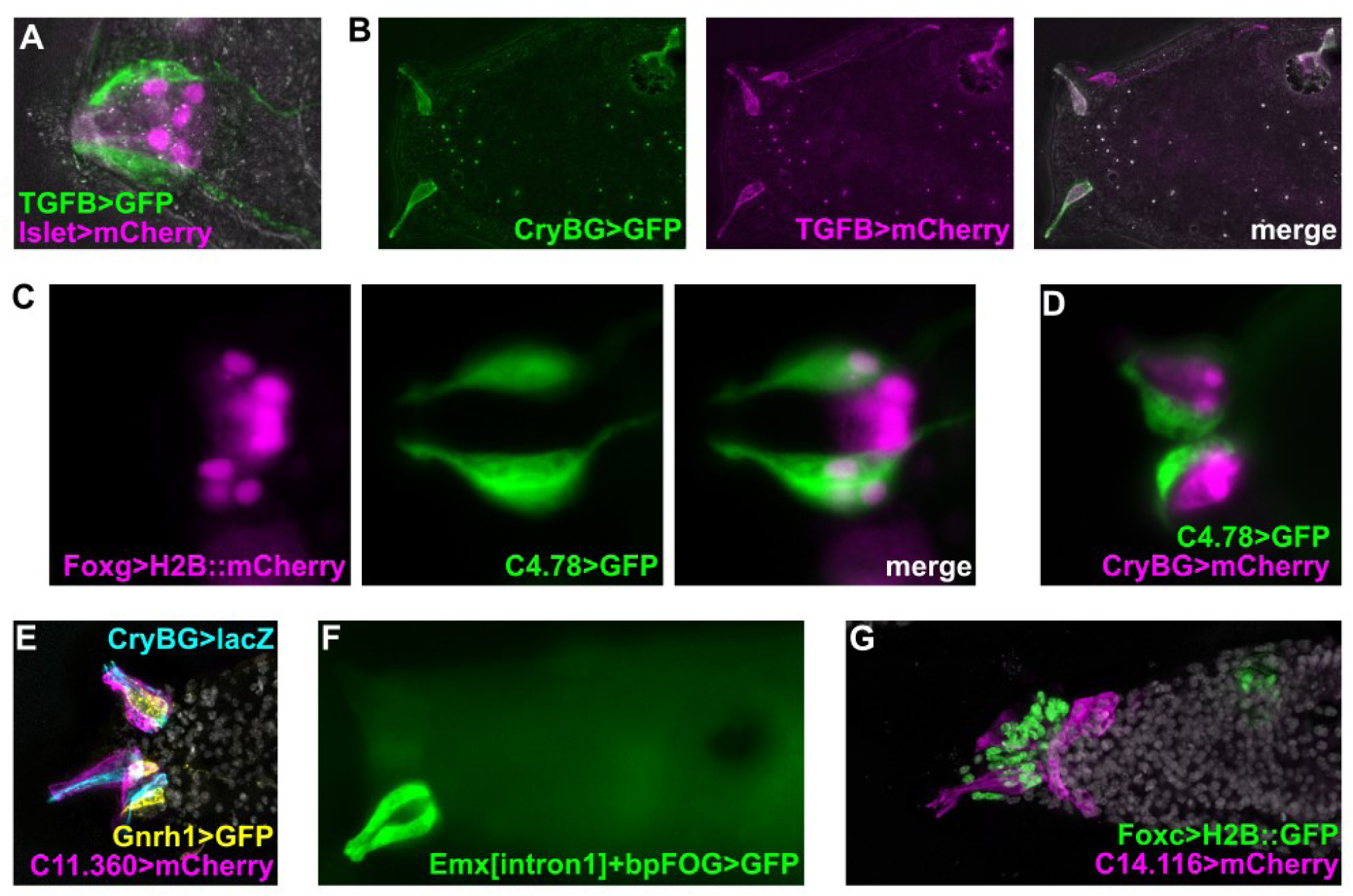
Additional marker genes and reporter plasmids expressed in papillae. A) *TGFB>Unc-76::GFP* reporter (green) is not co-expressed in the same cells as the *Islet intron 1 + -473/- 9>mCherry* reporter (pink). B) Cross-talk between *CryBG>Unc-76::GFP* and *TGFB>Unc-76::mCherry* reporter plasmids, showing aberrant co-expression in ACCs and/or PNs only when co-electroporated. C) Double electroporation showing that *Foxg>H2B::mCherry* (pink nuclei) is expressed in PNs labeled by *C4.78>Unc-76::GFP* reporter (green). D) Mutually exclusive expression of *C4.78>Unc-76::GFP* in PNs (green) and *CryBG>mCherry* in ACCs (pink). E) Mutually exclusive expression of *CryBG>lacZ* in ACCs (cyan), *Gnrh1>Unc-76::GFP* in PNs (yellow), and *C11.360>Unc-76::mCherry* in ICs (magenta), with DAPI counterstained in grey. This larva is the same as in main Figure 2G, with an additional channel and different false coloring. F) Reporter plasmid containing the 1^st^ intronic region of *Emx* drives expression in ICs, likely corresponding to the “ring” of late *Emx* expression in *Islet+* cells reported in Wagner et al. 2014 and distinct from earlier *Emx* expression in the papilla lineage as decribed in Liu and Satou 2019. G) *C14.116>Unc-76::mCherry* reporter expressed in central cells (ACCs+ICs, pink) and Basal Cells around the three papillae. DAPI in grey.

**Supplemental Figure 3.**
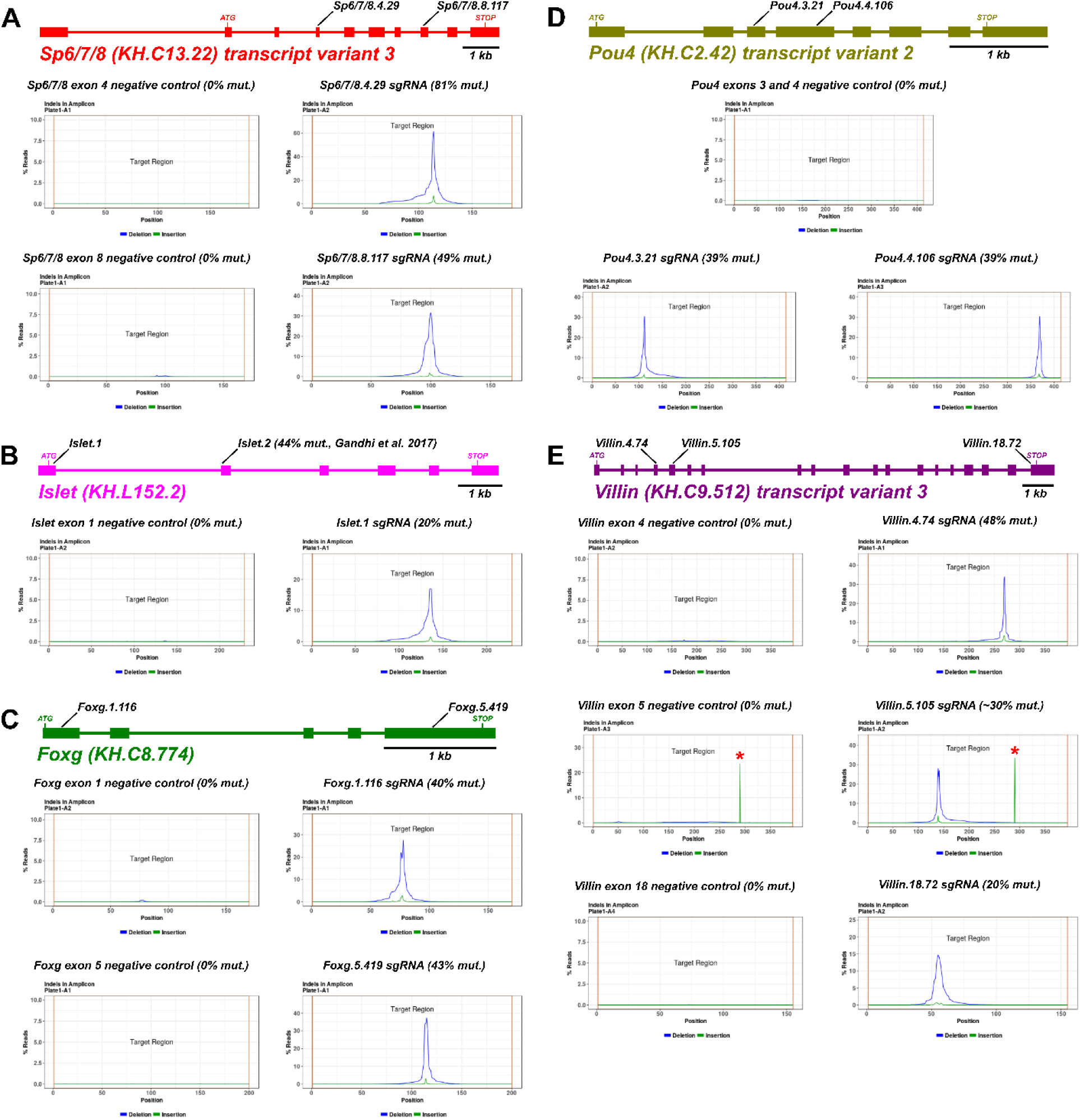
Validation of sgRNAs for CRISPR/Cas9-mediated mutagenesis. Gene loci diagrams for the four transcription factor-encoding genes investigated in this study: *Sp6/7/8, Foxg, Islet,* and *Pou4.* Plots underneath each gene show validation by Illumina sequencing (“Next-generation sequencing”, or NGS) of amplicons, performed as “Amplicon-EZ” service by Azenta. Mutagenesis efficacies are calculated by this service, and histograms of mapped reads show specificity of indels elicited by each sgRNA. Negative control amplicons are amplified from samples that were electroporated with no sgRNA, *U6>Control* sgRNA, or sgRNAs targeting unrelated amplicon regions. Note different y axis scales for each plot. Asterisks in *Villin* exon 5 amplicon plot indicate naturally occurring indel. Precise calculation of mutagenesis efficacy for *Villin.5.105* sgRNA was not given due to this natural indel.

**Supplemental Figure 4.**
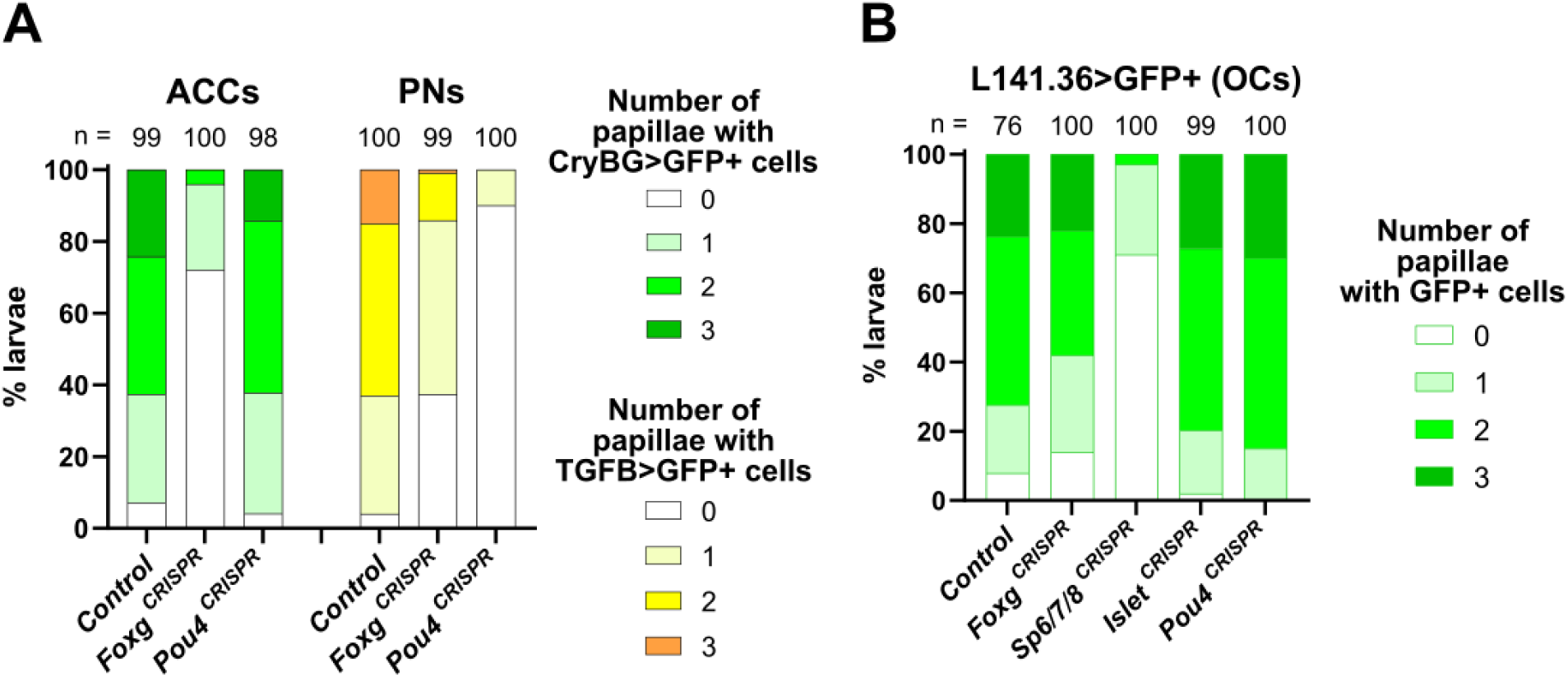
Effect of various CRISPR knockouts on specification of ACCs, PNs, and OCs. A) Scoring of effect of papilla-specific CRISPR knockout of *Foxg* or *Pou4* on specification of ACCs and PNs. Embryos were electroporated with *Foxc>H2B::mCherry, Foxc>Cas9, CryBG>Unc-76::GFP* (ACC reporter) or *TGFB>Unc-76::GFP* (PN reporter), and gene-specific sgRNA combinations (see below for specific combinations). B) Scoring of effects of papilla-specific CRISPR knockouts on OC specification as assayed by *L141.36>Unc-76::GFP* reporter. All were performed in parallel, but some are represented in Figure 4 also. For all plots, only larvae showing *Foxc>H2B::mCherry* expression in the papillae were scored. Specific sgRNAs used: *Foxg: U6>Foxg.1.116 + U6>Foxg.5.419; Pou4: U6>Pou4.3.21 + U6>Pou4.4.106; Sp6/7/8: U6>Sp6/7/8.4.29 + U6>Sp6/7/8.8.117; Islet: U6>Islet.2; Control: U6>Control*.

**Supplemental Figure 5.**
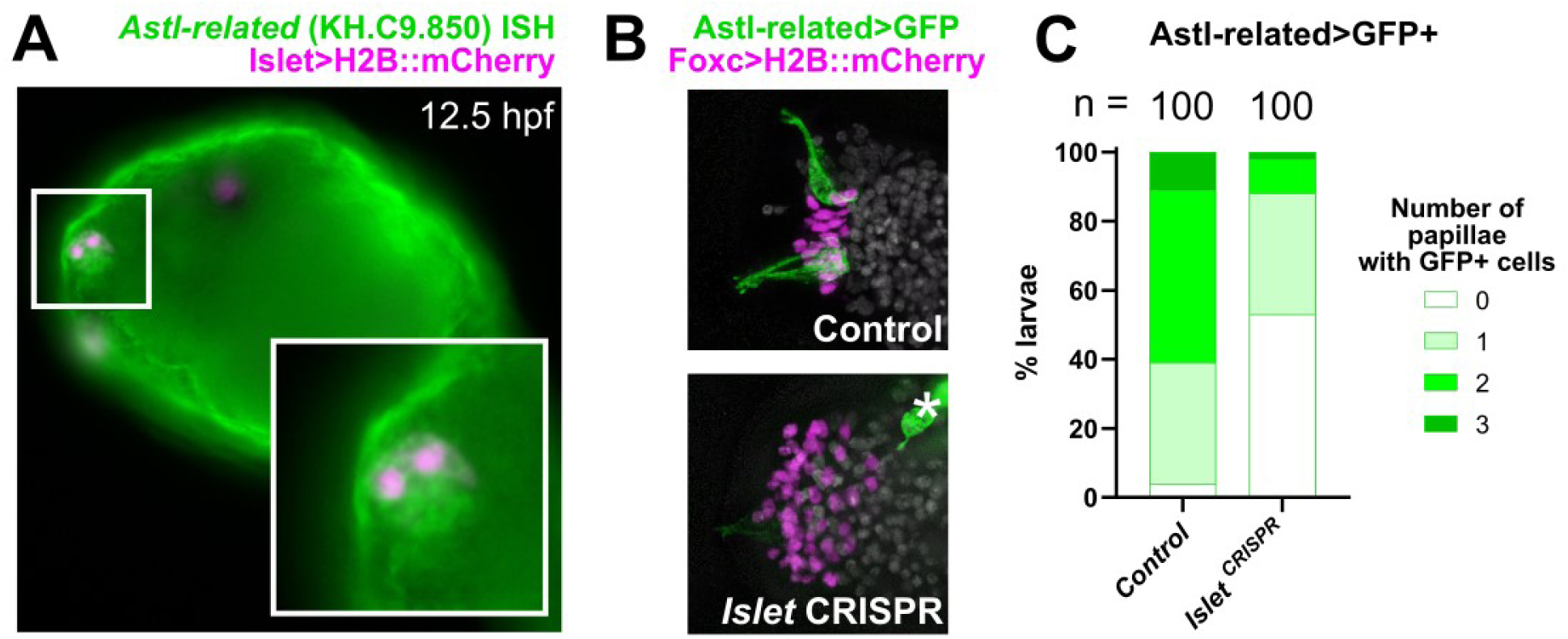
Validation of *Astl-related,* a transcriptional target of Islet confirmed by RNAseq. A) *In situ* mRNA hybridization (ISH) showing expression of *Astl-related* (green) specifically in the *Islet+* cells of the papillae (labeled by *Islet intron 1 + -473/-9>mCherry*, pink nuclei) B) Tissue-specific CRISPR/Cas-mediated mutagenesis of *Islet* results in loss of *Astl-related>Unc-76::GFP* reporter expression in ACCs/ICs (green). *Foxc>Cas9* was used to restrict CRISPR/Cas9 to the papilla territory (labeled by *Foxc>H2B::mCherry,* pink nuclei). Asterisk denotes residual reporter expression in cells outside the papilla territory. C) Scoring of larvae represented in panel B, following criteria used for Figure 3.

**Supplemental Figure 6.**
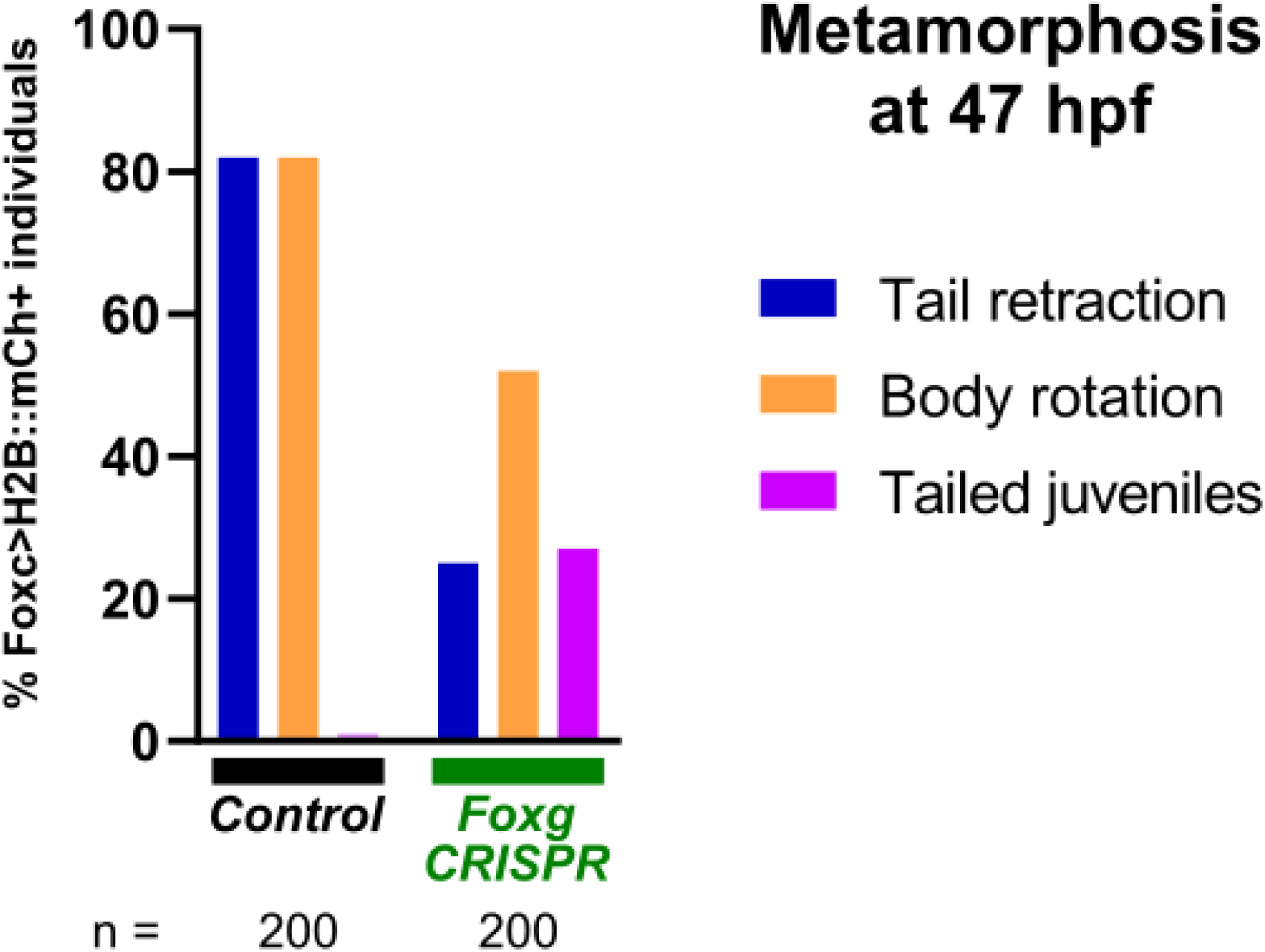
Replicate of *Foxg* CRISPR effects on metamorphosis. Scoring of individuals represented in Figure 7D. See supplemental sequence file for detailed plasmid electroporation recipes.

## REFERENCES

Abitua, P.B., Gainous, T.B., Kaczmarczyk, A.N., Winchell, C.J., Hudson, C., Kamata, K., Nakagawa, M., Tsuda, M., Kusakabe, T.G., Levine, M., 2015. The pre-vertebrate origins of neurogenic placodes. Nature.

Abitua, P.B., Wagner, E., Navarrete, I.A., Levine, M., 2012. Identification of a rudimentary neural crest in a non-vertebrate chordate. Nature 492, 104.

Afgan, E., Nekrutenko, A., Grüning, B.A., Blankenberg, D., Goecks, J., Schatz, M.C., Ostrovsky, A.E., Mahmoud, A., Lonie, A.J., Syme, A., 2022. The Galaxy platform for accessible, reproducible and collaborative biomedical analyses: 2022 update. Nucleic acids research.

Beh, J., Shi, W., Levine, M., Davidson, B., Christiaen, L., 2007. FoxF is essential for FGF-induced migration of heart progenitor cells in the ascidian *Ciona intestinalis*. Development 134, 3297–3305.

Caicci, F., Zaniolo, G., Burighel, P., Degasperi, V., Gasparini, F., Manni, L., 2010. Differentiation of papillae and rostral sensory neurons in the larva of the ascidian Botryllus schlosseri (Tunicata). Journal of Comparative Neurology 518, 547–566.

Cao, C., Lemaire, L.A., Wang, W., Yoon, P.H., Choi, Y.A., Parsons, L.R., Matese, J.C., Levine, M., Chen, K., 2019. Comprehensive single-cell transcriptome lineages of a proto-vertebrate. Nature 571, 349–354.

Chacha, P.P., Horie, R., Kusakabe, T.G., Sasakura, Y., Singh, M., Horie, T., Levine, M., 2022. Neuronal identities derived by misexpression of the POU IV sensory determinant in a protovertebrate. Proceedings of the National Academy of Sciences 119, e2118817119.

Chen, J.S., Pedro, M.S., Zeller, R.W., 2011. miR-124 function during Ciona intestinalis neuronal development includes extensive interaction with the Notch signaling pathway. Development 138, 4943–4953.

Christiaen, L., Wagner, E., Shi, W., Levine, M., 2009a. Electroporation of transgenic DNAs in the sea squirt Ciona. Cold Spring Harbor Protocols 2009, pdb. prot5345.

Christiaen, L., Wagner, E., Shi, W., Levine, M., 2009b. Isolation of sea squirt (Ciona) gametes, fertilization, dechorionation, and development. Cold Spring Harbor Protocols 2009, pdb. prot5344.

DeBiasse, M.B., Colgan, W.N., Harris, L., Davidson, B., Ryan, J.F., 2020. Inferring tunicate relationships and the evolution of the tunicate Hox cluster with the genome of Corella inflata. Genome Biology and Evolution 12, 948–964.

Delsuc, F., Philippe, H., Tsagkogeorga, G., Simion, P., Tilak, M.-K., Turon, X., López-Legentil, S., Piette, J., Lemaire, P., Douzery, E.J.P., 2018. A phylogenomic framework and timescale for comparative studies of tunicates. BMC Biology 16, 39.

Diogo, R., Kelly, R.G., Christiaen, L., Levine, M., Ziermann, J.M., Molnar, J.L., Noden, D.M., Tzahor, E., 2015. A new heart for a new head in vertebrate cardiopharyngeal evolution. Nature 520, 466.

Dolcemascolo, G., Pennati, R., De Bernardi, F., Damiani, F., Gianguzza, M., 2009. Ultrastructural comparative analysis on the adhesive papillae of the swimming larvae of three ascidian species. Invertebrate Survival Journal 6, S77–S86.

Fodor, A.C.A., Liu, J., Turner, L., Swalla, B.J., 2021. Transitional chordates and vertebrate origins: Tunicates. Current Topics in Developmental Biology 139, 325–374.

Gandhi, S., Haeussler, M., Razy-Krajka, F., Christiaen, L., Stolfi, A., 2017. Evaluation and rational design of guide RNAs for efficient CRISPR/Cas9-mediated mutagenesis in Ciona. Developmental biology 425, 8–20.

Gans, C., Northcutt, R.G., 1983. Neural crest and the origin of vertebrates: a new head. Science 220, 268–273.

Haeussler, M., Schönig, K., Eckert, H., Eschstruth, A., Mianné, J., Renaud, J.-B., Schneider-Maunoury, S., Shkumatava, A., Teboul, L., Kent, J., 2016. Evaluation of off-target and on-target scoring algorithms and integration into the guide RNA selection tool CRISPOR. Genome biology 17, 1–12.

Haupaix, N., Abitua, P.B., Sirour, C., Yasuo, H., Levine, M., Hudson, C., 2014. Ephrin-mediated restriction of ERK1/2 activity delimits the number of pigment cells in the Ciona CNS. Developmental Biology 394, 170–180.

Haupaix, N., Stolfi, A., Sirour, C., Picco, V., Levine, M., Christiaen, L., Yasuo, H., 2013. p120RasGAP mediates ephrin/Eph-dependent attenuation of FGF/ERK signals during cell fate specification in ascidian embryos. Development 140, 4347–4352.

Horie, R., Hazbun, A., Chen, K., Cao, C., Levine, M., Horie, T., 2018. Shared evolutionary origin of vertebrate neural crest and cranial placodes. Nature 560, 228.

Hotta, K., Dauga, D., Manni, L., 2020. The ontology of the anatomy and development of the solitary ascidian Ciona: the swimming larva and its metamorphosis. Scientific Reports 10, 17916.

Hozumi, A., Matsunobu, S., Mita, K., Treen, N., Sugihara, T., Horie, T., Sakuma, T., Yamamoto, T., Shiraishi, A., Hamada, M., 2020. GABA-Induced GnRH Release Triggers Chordate Metamorphosis. Current Biology.

Hudson, C., Yasuo, H., 2006. A signalling relay involving Nodal and Delta ligands acts during secondary notochord induction in *Ciona* embryos. Development 133, 2855–2864.

Ikuta, T., Saiga, H., 2007. Dynamic change in the expression of developmental genes in the ascidian central nervous system: revisit to the tripartite model and the origin of the midbrain-hindbrain boundary region. Dev Biol 312.

Imai, J.H., Meinertzhagen, I.A., 2007. Neurons of the ascidian larval nervous system in Ciona intestinalis: II. Peripheral nervous system. Journal of Comparative Neurology 501, 335–352.

Karaiskou, A., Swalla, B.J., Sasakura, Y., Chambon, J.P., 2015. Metamorphosis in solitary ascidians. Genesis 53, 34–47.

Kari, W., Zeng, F., Zitzelsberger, L., Will, J., Rothbaecher, U., 2016. Embryo microinjection and electroporation in the chordate Ciona intestinalis. JoVE (Journal of Visualized Experiments), e54313.

Khurana, S., George, S.P., 2008. Regulation of cell structure and function by actin-binding proteins: villin’s perspective. FEBS letters 582, 2128–2139.

Kocot, K.M., Tassia, M.G., Halanych, K.M., Swalla, B.J., 2018. Phylogenomics offers resolution of major tunicate relationships. Molecular Phylogenetics and Evolution 121, 166–173.

Kusakabe, T.G., Sakai, T., Aoyama, M., Kitajima, Y., Miyamoto, Y., Takigawa, T., Daido, Y., Fujiwara, K., Terashima, Y., Sugiuchi, Y., 2012. A conserved non-reproductive GnRH system in chordates.

Lemaire, P., 2011. Evolutionary crossroads in developmental biology: the tunicates. Development 138, 2143–2152.

Liu, B., Ren, X., Satou, Y., 2023. BMP signaling is required to form the anterior neural plate border in ascidian embryos. Development Genes and Evolution, 1–11.

Liu, B., Satou, Y., 2019. Foxg specifies sensory neurons in the anterior neural plate border of the ascidian embryo. Nature communications 10, 1–10.

Martik, M.L., Bronner, M.E., 2021. Riding the crest to get a head: neural crest evolution in vertebrates. Nature Reviews Neuroscience 22, 616–626.

Matsunobu, S., Sasakura, Y., 2015. Time course for tail regression during metamorphosis of the ascidian Ciona intestinalis. Developmental biology 405, 71–81.

Nakayama-Ishimura, A., Chambon, J.-p., Horie, T., Satoh, N., Sasakura, Y., 2009. Delineating metamorphic pathways in the ascidian Ciona intestinalis. Developmental biology 326, 357–367.

Papadogiannis, V., Pennati, A., Parker, H.J., Rothbächer, U., Patthey, C., Bronner, M.E., Shimeld, S.M., 2022. Hmx gene conservation identifies the origin of vertebrate cranial ganglia. Nature 605, 701–705.

Pasini, A., Amiel, A., Rothbächer, U., Roure, A., Lemaire, P., Darras, S., 2006. Formation of the ascidian epidermal sensory neurons: insights into the origin of the chordate peripheral nervous system. PLoS biology 4, e225.

Patthey, C., Schlosser, G., Shimeld, S.M., 2014. The evolutionary history of vertebrate cranial placodes–I: cell type evolution. Developmental biology 389, 82–97.

Pennati, R., Ficetola, G.F., Brunetti, R., Caicci, F., Gasparini, F., Griggio, F., Sato, A., Stach, T., Kaul-Strehlow, S., Gissi, C., Manni, L., 2015. Morphological Differences between Larvae of the Ciona intestinalis Species Complex: Hints for a Valid Taxonomic Definition of Distinct Species. PLOS ONE 10, e0122879.

Pennati, R., Groppelli, S., De Bernardi, F., Mastrototaro, F., Zega, G., 2009. Immunohistochemical analysis of adhesive papillae of Clavelina lepadiformis (Müller, 1776) and Clavelina phlegraea (Salfi, 1929)(Tunicata, Ascidiacea). European journal of histochemistry: EJH 53.

Pennati, R., Zega, G., Groppelli, S., De Bernardi, F., 2007. Immunohistochemical analysis of the adhesive papillae of Botrylloides leachi (Chordata, Tunicata, Ascidiacea): Implications for their sensory function. Italian Journal of Zoology 74, 325–329.

Poncelet, G., Shimeld, S.M., 2020. The evolutionary origins of the vertebrate olfactory system. Open Biology 10, 200330.

Poncelet, G.J.F., Parolini, L., Shimeld, S., 2022. A microfluidic device for controlled exposure of transgenic Ciona intestinalis larvae to chemical stimuli demonstrates they can respond to carbon dioxide. bioRxiv, 2022-2008.

Pottin, K., Hyacinthe, C., Rétaux, S., 2010. Conservation, development, and function of a cement gland-like structure in the fish Astyanax mexicanus. Proceedings of the National Academy of Sciences 107, 17256–17261.

Razy-Krajka, F., Lam, K., Wang, W., Stolfi, A., Joly, M., Bonneau, R., Christiaen, L., 2014. Collier/OLF/EBF-dependent transcriptional dynamics control pharyngeal muscle specification from primed cardiopharyngeal progenitors. Developmental cell 29, 263–276.

Rétaux, S., Pottin, K., 2011. A question of homology for chordate adhesive organs. Communicative & Integrative Biology 4, 75–77.

Roure, A., Chowdhury, R., Darras, S., 2022. Regulation of anterior neurectoderm specification and differentiation by BMP signaling in ascidians. bioRxiv, 2022-2010.

Roure, A., Darras, S., 2016. Msxb is a core component of the genetic circuitry specifying the dorsal and ventral neurogenic midlines in the ascidian embryo. Developmental biology 409, 277–287.

Sakamoto, A., Hozumi, A., Shiraishi, A., Satake, H., Horie, T., Sasakura, Y., 2022. The TRP channel PKD2 is involved in sensing the mechanical stimulus of adhesion for initiating metamorphosis in the chordate Ciona. Development, Growth & Differentiation 64, 395–408.

Sasakura, Y., Nakashima, K., Awazu, S., Matsuoka, T., Nakayama, A., Azuma, J., Satoh, N., 2005. Transposon-mediated insertional mutagenesis revealed the functions of animal cellulose synthase in the ascidian *Ciona intestinalis*. Proceedings of the National Academy of Sciences of the United States of America 102, 15134.

Satija, R., Farrell, J.A., Gennert, D., Schier, A.F., Regev, A., 2015. Spatial reconstruction of single-cell gene expression data. Nature biotechnology 33, 495.

Satoh, N., 2013. Developmental genomics of ascidians. John Wiley & Sons.

Satou, Y., Sato, A., Yasuo, H., Mihirogi, Y., Bishop, J., Fujie, M., Kawamitsu, M., Hisata, K., Satoh, N., 2021. Chromosomal inversion polymorphisms in two sympatric ascidian lineages. Genome biology and evolution 13, evab068.

Satou, Y., Tokuoka, M., Oda-Ishii, I., Tokuhiro, S., Ishida, T., Liu, B., Iwamura, Y., 2022. A manually curated gene model set for an ascidian, Ciona robusta (Ciona intestinalis type A). Zoological Science 39.

Sharma, S., Wang, W., Stolfi, A., 2019. Single-cell transcriptome profiling of the Ciona larval brain. Developmental Biology 448, 226–236.

Shimeld, S.M., Purkiss, A.G., Dirks, R.P.H., Bateman, O.A., Slingsby, C., Lubsen, N.H., 2005. Urochordate βγ-crystallin and the evolutionary origin of the vertebrate eye lens. Current biology 15, 1684–1689.

Song, M., Yuan, X., Racioppi, C., Leslie, M., Stutt, N., Aleksandrova, A., Christiaen, L., Wilson, M.D., Scott, I.C., 2022. GATA4/5/6 family transcription factors are conserved determinants of cardiac versus pharyngeal mesoderm fate. Science Advances 8, eabg0834.

Stolfi, A., Gandhi, S., Salek, F., Christiaen, L., 2014. Tissue-specific genome editing in Ciona embryos by CRISPR/Cas9. Development 141, 4115–4120.

Stolfi, A., Levine, M., 2011. Neuronal subtype specification in the spinal cord of a protovertebrate. Development 138, 995–1004.

Stolfi, A., Wagner, E., Taliaferro, J.M., Chou, S., Levine, M., 2011. Neural tube patterning by Ephrin, FGF and Notch signaling relays. Development 138, 5429–5439.

Tang, W.J., Chen, J.S., Zeller, R.W., 2013. Transcriptional regulation of the peripheral nervous system in Ciona intestinalis. Developmental biology 378, 183–193.

Tolkin, T., Christiaen, L., 2016. Rewiring of an ancestral Tbx1/10-Ebf-Mrf network for pharyngeal muscle specification in distinct embryonic lineages. Development 143, 3852–3862.

Torrence, S.A., Cloney, R.A., 1983. Ascidian larval nervous system: primary sensory neurons in adhesive papillae. Zoomorphology 102, 111–123.

Turon, X., 1991. Morphology of the adhesive papillae of sorne ascidian larvae. Cah. Biol. Mar 32, 295–309.

Wagner, E., Stolfi, A., Choi, Y.G., Levine, M., 2014. Islet is a key determinant of ascidian palp morphogenesis. Development 141, 3084–3092.

Wakai, M.K., Nakamura, M.J., Sawai, S., Hotta, K., Oka, K., 2021. Two-Round Ca2+ transient in papillae by mechanical stimulation induces metamorphosis in the ascidian Ciona intestinalis type A. Proceedings of the Royal Society B 288, 20203207.

Waki, K., Imai, K.S., Satou, Y., 2015. Genetic pathways for differentiation of the peripheral nervous system in ascidians. Nature communications 6, 8719.

Zeng, F., Wunderer, J., Salvenmoser, W., Ederth, T., Rothbächer, U., 2019a. Identifying adhesive components in a model tunicate. Philosophical Transactions of the Royal Society B 374, 20190197.

Zeng, F., Wunderer, J., Salvenmoser, W., Hess, M.W., Ladurner, P., Rothbächer, U., 2019b. Papillae revisited and the nature of the adhesive secreting collocytes. Developmental biology 448(2), 183–198.

