## Supplemental Sequences File for "Specification of distinct cell types in a sensory-adhesive organ for metamorphosis in the *Ciona* larva"

**Supplemental sequence file**

Sequences based on genome assemblies and might not be verified by sequencing after cloning. ANISEED unique gene IDs at end of document.

>Islet intron 1 (0 to 2014) + -473/-9 (mutated ATG) XhoI

(originally from Wagner et al. 2014)

gcctcgcttaattgcggtaagtttgtgggttgtttaataaagtaggggggtttgggttcgaaatcaggccagggttgaaaatccagggatcccgggttctaatccaggtcagggttggagaaccagggatcctgggttcgaattcaggccaagaactagcattgtacgcgtaagtgtcttgaacaaaattgcgggtaagtgcttaagcaggaaaggcaacaattaaccttaggattcagtatatagtgcgaagacggatacattattaatgggggggcgacctttgtaagcgcctaacattggactgggccggggggttggggcgcgatgttcgcgatgaaattggatttttatcttttggcaacaaaattaccgttcgcgtagatcgccatctttaaaccgcgcggcgaatatctcaagccattgtcacgtcacaatcggcatttattatttatttacgtcacaacggcttaatctgaagctgcaaaaggtatttgtgacatcacaaaacataacaatgaagttgtgttaaagcattgtgaatcacaaatacgagataatatacatcgcttatgtcgtcataaccgcgtcgatgcagccgcgtaaatcgcactcaatgtcccattcaagctgctacgtcacaatcgctattctttccccacaatagacaatgttagttaacaatttcccgtgtgtgacgcaacaattaaaacaataaagaacaatgcagtgtattctaatacattgtagaatgctcgcgtcatcttacagcgcacaccgcggttgcatcgaatgaataatgcaacaacatatcgacccgcattcgataaaatatcagacttttatattttccttaaagacctcaagtattttaactttttttcgaaaatcttttttattttaataatattttcgcagtgacgtcattaatttatgacatcaccagtgatatcatgacgtcactcgatcaatattagccaagagcaaaaatagtggatatttttcattaattctttatttgtttatttaaccggaaacaaaatggccgccggtgacctggttgattgtgacgtatataacacgcaacacacgactattatgacgaaacacagacttgaattgaaaatctagaaccccggtcacgttatggtaatgcttatgacgaaataattgaaagtttttgcgattgtgacgtcataatacctttattgttaggcgattggttaattaggacgcgtgttcgtttttaattgtgacgtaataaacatgaaattaaacgagattgccaaagttaaatgatgcattatgacatcataatatataacttttatgtgattctagaaaagacgaaaccgcgaaagtagaacaagacgattgttaacaaagctacgtcataatccataatcaacacgcgaatcgcgtcgctaagtatccgcacctgtgacgtaacaatactaagacacaggcaaacttcgatatcgctaaaatatatgatttctatgcgatcgcgattaagtggttttaattgttcgctagcaaagtggtttgtggagattaatttgaggtaatgtagtttagaaccaacttattgtaaattatctaaagtggtcacatgtatacgcacgtgtatatggattaattaaggggggtcattaagtgtaaagtcgctaatccccctcacccgtgagactttagtcattaacaaagtgtttgttaattgatcgacaaagtgggtcgcggtgtaatagttgctgcattatgacataacatgcattataaatgtgacgtcacattgaataaacaatcgtctatatttggattttaggacttgagataaaaaagttaaaatatgtttgggtcttgatggcttgtaacttttgttggggattattttgttaattgttgggagggtcaggtttttaagagctttgttaagcgttaaataaagaaagaaagaaatttaaataaattaatttaattttagattgaactttcttaaacacagaaataaaataaagtttttgtttggcctcgagggttaacttaacatgggcgtgtgaaatgtaattagttaatgtagttagtcgtatctggatcggaaggcattcatttgcactgtaggcattgaactaggaggttcaccacatcgaggaatatttggtcgcgctgcggggattgagggccttgtagttggggggaggtttgaaagtttcccgaagtaatattaactttgttgtataggacccgttgttgctgttagagagaatcggagaaaaagtttcgagagaagctttagataaattgattctgtcgcattcgagtattacaacttgtgtctgtgacgtgagcgagtgaggacaatttactgtgacgtcataatagccagcaatataggggtggaaggtgaataccccgctctacaagcggagaatttcgaaatttcaaacctggatcccggatactcgccgccgttggcggacgactcaaggtcggggtcgttcacaagttttatcaacgaa

>Islet intron 1 (0 to 2014) + bpFOG XhoI

gcctcgcttaattgcggtaagtttgtgggttgtttaataaagtaggggggtttgggttcgaaatcaggccagggttgaaaatccagggatcccgggttctaatccaggtcagggttggagaaccagggatcctgggttcgaattcaggccaagaactagcattgtacgcgtaagtgtcttgaacaaaattgcgggtaagtgcttaagcaggaaaggcaacaattaaccttaggattcagtatatagtgcgaagacggatacattattaatgggggggcgacctttgtaagcgcctaacattggactgggccggggggttggggcgcgatgttcgcgatgaaattggatttttatcttttggcaacaaaattaccgttcgcgtagatcgccatctttaaaccgcgcggcgaatatctcaagccattgtcacgtcacaatcggcatttattatttatttacgtcacaacggcttaatctgaagctgcaaaaggtatttgtgacatcacaaaacataacaatgaagttgtgttaaagcattgtgaatcacaaatacgagataatatacatcgcttatgtcgtcataaccgcgtcgatgcagccgcgtaaatcgcactcaatgtcccattcaagctgctacgtcacaatcgctattctttccccacaatagacaatgttagttaacaatttcccgtgtgtgacgcaacaattaaaacaataaagaacaatgcagtgtattctaatacattgtagaatgctcgcgtcatcttacagcgcacaccgcggttgcatcgaatgaataatgcaacaacatatcgacccgcattcgataaaatatcagacttttatattttccttaaagacctcaagtattttaactttttttcgaaaatcttttttattttaataatattttcgcagtgacgtcattaatttatgacatcaccagtgatatcatgacgtcactcgatcaatattagccaagagcaaaaatagtggatatttttcattaattctttatttgtttatttaaccggaaacaaaatggccgccggtgacctggttgattgtgacgtatataacacgcaacacacgactattatgacgaaacacagacttgaattgaaaatctagaaccccggtcacgttatggtaatgcttatgacgaaataattgaaagtttttgcgattgtgacgtcataatacctttattgttaggcgattggttaattaggacgcgtgttcgtttttaattgtgacgtaataaacatgaaattaaacgagattgccaaagttaaatgatgcattatgacatcataatatataacttttatgtgattctagaaaagacgaaaccgcgaaagtagaacaagacgattgttaacaaagctacgtcataatccataatcaacacgcgaatcgcgtcgctaagtatccgcacctgtgacgtaacaatactaagacacaggcaaacttcgatatcgctaaaatatatgatttctatgcgatcgcgattaagtggttttaattgttcgctagcaaagtggtttgtggagattaatttgaggtaatgtagtttagaaccaacttattgtaaattatctaaagtggtcacatgtatacgcacgtgtatatggattaattaaggggggtcattaagtgtaaagtcgctaatccccctcacccgtgagactttagtcattaacaaagtgtttgttaattgatcgacaaagtgggtcgcggtgtaatagttgctgcattatgacataacatgcattataaatgtgacgtcacattgaataaacaatcgtctatatttggattttaggacttgagataaaaaagttaaaatatgtttgggtcttgatggcttgtaacttttgttggggattattttgttaattgttgggagggtcaggtttttaagagctttgttaagcgttaaataaagaaagaaagaaatttaaataaattaatttaattttagattgaactttcttaaacacagaaataaaataaagtttttgtttggcctcgagcagctgaagcttgcatgcctgcaggtcgactctagaggatccggcaaagcttcgtgtattgtaccggcccattgtcaatcatgcaaacttgatattatattgacaagagaagaaggcagtttaaattaaaactctaaagtagagagacattaatctcagctgacaaggcaggtggtcacagtaagttcatttaaatagttggccaacaatagcctttccaagaaagtatttttgttccaggtctatacaaaaataacacacatagc

>CryBG -1068/-24 (Based on Shimeld et al. 2005)

taattcttactgttcggttgaaactcttgaatccactatgacgtcatcatcgtcgctacaaccgctataaacagcgagaaacaaaataaaaacaatcgacttcattacggaagagttgggtcgaaagtggttgttgcatgaaatatatgtgtgttatatctcggtgctaagtggtacaacaaacaaataaaacgtctggtttatgagatgctcgcgtggtacgggctatgtgatgtcataatacctttgtctattatccttgttacgacataattaccatgaaacagcattgtgacgtcacagtgctgtttgtgttggacaaacaacacgtggtttctacgagggggaatcccctctctgtcattattacgtatgatgacgtataatatcgattattattcaccatccaaggttatatacgatatctatttggttttatccgcttggtggcgacatcaatcctcaactacaattggtattatgacgtagttcgtagttacgtcacaagtcacacaacaaagaatttattatgagcaaataaacgaactaatatgacgcaacaataactaacaatttatcacgtgatcgttttatttattttgtttgttatgtcataaccacttggtgcttaaaagggagcttatgacgtattaatgaactttaattattaaaacaatcttgcttatcgtgctgtgttacgtcataatacatgcattagcaattactttgttttgtcataatatgaattatgtaattatcgtgacgttgtttgttttaaaaacccatttattattacgtcataatactttagatggttgtgtatgttacgtcataatgttaattgttgaataaaaatgaagaaacaaagccatgaagaaggcggtttgttataaaacattgttgaagggatggaattaattatgttgttaacgtttgaattattgatgtaacaaggacgacaagtgagcgcaaatcagatttcgtttactttccttctaaccctcctaaccactgcttatttcgcattttgtacaatcgaagtttc

>Islet[BTN]+bpFOG (Islet -5915/-5356 (approximately) + bpFOG)

(from Stolfi et al. 2010)

attttgtgctaaccgtattcatatttgcaacccactgtgagttgcttcttattaggtttaacagccacggctttaactaaattttatttattttaaccttagcgtgttataacgactgttatttttttgttaattcccatttcttcgccatgttagttaatatgacttttgttattgtaattatataaaaatatatttttaaaataagtctaaaagtaaattccaaccaagttgtatttacaaatttccgcaccagaataggcagcaggcaaccgcggatcgaaacaagtccctgatttatggttttgttgcaactaattatggctgtacagtacacaggtcccaggattgccgtaaaacttaatttgctggaatatgttgaacaacagttgttacactctgttgctatattaagtttagcggccacgctttttttaagccttttattccagtattttcgctcgtagcgtgttgataacgactgttttgctgctaattctcttcgttgttttaggtaaacttcgctatcgtattcaaaactcgagcagctgaagcttgcatgcctgcaggtcgactctagaggatccggcaaagcttcgtgtattgtaccggcccattgtcaatcatgcaaacttgatattatattgacaagagaagaaggcagtttaaattaaaactctaaagtagagagacattaatctcagctgacaaggcaggtggtcacagtaagttcatttaaatagttggccaacaatagcctttccaagaaagtatttttgttccaggtctatacaaaaataacacacaacatg

>Foxg -2863/-3 (Loosely based on Cao et al. 2019, this variant showed weak and inconsistent expression, likely due to missing a key transcription start site sequence, and was replaced by the -2836/+54 driver further below)

gtttcattttccaacaaaacattataatgcgataagagaattggccatctttaccacagaacttaaatttaacggaaatataaagttgcaacgtacgtaaataacagaccacccgtatttattgcgtgacattaatacattggatgtcaaagaacaaagaacttgttgtttacgtcatataggatccggtacttgataaaccacttcaagaataagatttatgacatcataaccaaacatgagcgacgactttaattgggaaaaaatactttgaaatgttttttttatcagatatttgacaacaaaaatacaatagtcggaaaatattgtatttttcagttttatttgttcatatttaatatataagcacacctaacacctggctgataatgggacgtcactgggttaaaatagaaatcatataaagcacaagttcaattggtttcgggctaaaaattaaatcctttttcggtaaaccgaccgtttgtcacatcaaagccacagcgagcagtgataagagccgagtgtttttatgtaaacaggtttaggtaactcgtgtctgcttgatttggttgtaagattaattactcacagtgttacacatgttcccacaaacccagagacattgtaactggggcgtagatgatacgttataaaatataaaccaatacatgttgtataatataagtgcactcattttatccgcagagtgttataacgattttcgctttcaaaatatatagtagggtgggggaagatgggacacctttagcacataatatccaaatgttctcatcgcgttttgaaccattccaacggtctattgtcgtaaggatgcggttttgtgattctttgaatgttttttgtttactaccaagtgggacgagaaaatagaatgaaaatatgtcccatctttccccgctttaatatataactttttaaatctgtgtaatgtttcatttaaacatgttctgggctattgagttcttaaaacttatcaaacgtatgttggtatatctgcccgggcctttattattaaacaccaaaaccacatagaaaaagtaactattaaacgaaattctgtctatgtcagccgttaaactaacttgcagtgtgtatagatatcgttgggctctatttggtggtaaacaaagaatatttacagaattatataaccgtattcttacgactgtaaaagagtgttgttaattgtttaaaacacgaataaaaaatatgggagatcgtgtgctgatggttttcaccttatcacagttcagataaaaaattaaaagttctcaaaacaaactataggaacaaatatgtttttgatttatattattgccaaaatactatgttaaattggtctacctatttccaattaaaaaaacagtcacattttaattaattttacggaaacactgtacgttgtcagtgattttagatctgatcttataattgagtctggagtaaaacagaccttgtgattgacagtttgaattataagtatcagaaaacattgatcgtagatgtaaccaaagctctcatcagttgtggaccaaagtaaccacaaaacctacctaatttttgcagtgttgcctttagactaaacttttattccagtgttgccatttaaaaaaaaggtagatgtgaggtcggtgaataaaaagggtaaattattagttatatgaacgtagtttaagctataactagttaatacagatatattaaatcatgcagtttgcgtacaaattatattaattaaagtccattaaaaaatgtattcaatattacaaaaaaatcaatttctattaagaaaaacagaaaaacattaaaaaattggaattttttgacattttaatcgtggcaacatgacaaacttatgttagcagattatcacgccatcttaacatacagcacgtgtgaagcaagatcaaataaccaaggtgtcaagtgaaagttacagggcacaattatttaaccaagctaacgtaaatgaaaaacccaggcaaccggcttattactgctggctacttggtcagtgtggttaacatatatcaacggaaagccccgctctatatcatgacgtattaactagtattaacaagcccaacattgtacggtgtcgtgtaaaaccttatatggcatagagcttgttttgcggaaagctattttgtttttaaatctgaaaatatgagattgaatgtggtatgggaaaatgtactcctatagacacacctggtgtattcacacaccttatcctcactatgttgagttatagcatgggtaattcttatccaccacacatggtgtatttagtgtattcatacttagttataacagtattgtatatttaaatattgccaaaatccgtcgtagagtgacaaaaacaatcttgaaatactgcaaagtcattaggcgtattatctcacactctaaatcacttaaacacaataaagtcgcgagttttaaaagcctagatctaataataattgcaaatcaaacgcattatgtccattagttgtattttctttgtcgatttagtatccgggttatccttgttaacaaggtgtttgtgtgtctaagtatgtccatgattacgtcttcctaccagaggagtcgtgttgttttagtcaatacggtggtaattatccgattattgtgatatagcgagttacaggcgccctcagctcacggtggaagtgctaacttgtaagtcttaatgcatttctcttttcttcaaaggtttttgagtagattttcgtttgtaattataccggcgtctccttacgctgtaatccacgcttttaattgtgacgtggattgacagaaagagtagcgcgagagagaaacgcggcatacag

>Foxg -2863/+54 ATG start codon

gtttcattttccaacaaaacattataatgcgataagagaattggccatctttaccacagaacttaaatttaacggaaatataaagttgcaacgtacgtaaataacagaccacccgtatttattgcgtgacattaatacattggatgtcaaagaacaaagaacttgttgtttacgtcatataggatccggtacttgataaaccacttcaagaataagatttatgacatcataaccaaacatgagcgacgactttaattgggaaaaaatactttgaaatgttttttttatcagatatttgacaacaaaaatacaatagtcggaaaatattgtattttttagttttatttgttcatatttaatatataagcacacctaacacctggctgataatgggacgtcactgggttaaaatagaaatcatataaagcacaagttcaattggtttcgggctaaaaattaaatcctttttcggtaaaccgaccgtttgtcacatcaaagccacagcgagcagtgataagagccgagtgtttttatgtaaataggtttaggtaactcgtgtctgcttgatttggttgtaagattaattactcgcagtgttacacatgttcccacaaacccagagacattgtaactggggcgtagatgatacgttataaaatataaaccaacacatgttgtataatataagtgcactcattttatccgcagagtgttataacgattttcgcttttaaaatatatagtagggtgggggaagatgggacacctttagcacataatatccaaatgttctcatcgcgttttgaaccatttcaacggtctattgtcgtaaggatacggttttgtaaatctttgaatgttttttgtttactaccaagtgggacgagaaattagaatgaaaaggtgtcccatcttttcccgctctaatatataactttttaaatctgtgtaatgtttcatttaaagatgttctgggctattgagttcttaaaacttatcaaacgtatgttggtatatctgcccggacctttattattaaacaccaaaaccacatagaaaaagtaactattaaacgaaattctgtctatgtcagccgttaaactaacttgcagtgtgtatagatatcgttgggctctatttggtggtaaacaaagaatatttacagaattatataaccgtattcttacgactgtaaaagagtgttgttaattgtttaaaacacgaataaaaaatatgggagatcgtgtgctgatggttttcaccttatcacagttcagataaaaaattaaaagttctcaaaacaaactataggaacaaatatgtttttgatttatattattgcaaaaatactatgttaaattggcctacctatttccaattaaaaaaagtcacattttaattaattttacggaaacactgtacgttgtcagtgattttagacctgatcttataattgagtctggagtaaaacagaccttgtgattgacagtttgaattataagtatcagaaaacattgatcgtagatgtaaccaaagctctcatcagttgtggaccaaagtaaccacaaaacctacctaatttttgcagtgttgcctttagactaaacttttatttcagtgttgccatttaaaaaaaaagctagatgtgaggtcggtgaataaaaagggtaaattattagttatatgagcgtagtttaagctataactagttaatacagatatattaaatcatgcagtttgcgtacaaattatattaattaaagtccattaaaaaatgtattcaatattacaaaaaaatcaatttctattaagaaaaacagaaaaacattaaaaaattggaattttttgacattttaatcgtggcaacatgacaaacttatgttagcagattatcacgccatcttaacatacagcacgtgtgaagcaagatcaaataaccaaggtgtcaagtgaaagttacagggcacaattatttaaccaagctaacgtaaatgaaaaacccaggcaaccggcttatcactgctggctacttggtcagtgtggttaacatatatcaacggaaagccccgctctatatcatgacgtattaactagtattaacaagcccaacattgtacggtgtcgtgtaaaaccttatatggcatgaagcttgttttgcggaaagctattttgttttttaatctgaaaatatgagatgggtgtggtatgggaaaatgaacgcctatagacacacctggtgtattcacacaccttatcctcactatgttgagttatagcatgggtaattcttatccaccacacatggtgtatttagtgtattcatacttagttataacagtattgtatatttaaatattgccaaaatccgtcgtagagtgacaaaaacaatcttgaaatactgcaaagtcattaggcgtattatctcacactctaaatcacttaaacacaataaagtcgcgagttttaaaagcctagatctaataataattgcaaatcaaacgcattatgtccattagttgtattttctttgtcgatttagtatccgggttatccttgttaacaaggtgtttgtgtgtctaagtatgtccatgattacgtcttcctaccagaggagtcgtgttgttttagtcaatacggtggtaattatccgattattgtgatatagcgagttacaggcgccctcagctcacggtggaagtgctaacttgtaagtcttaatgcatttctcttttcttcaaaggtttttgagtagattttcgtttgtaattataccggcgtctccttacgctgtaatccacgcttttaattgtgacgtggattgacagaaagagtagcgcgagagagaaacgcggcatacagaatATGACGAACGACGCGGCCGAGTCTGGTCATTCAAAGCGAGAAAACTTCATAGAG

>Foxc -2132/-1 (Based on Wagner and Levine 2012)

ccgcctacgtagggtaaataccgctcagggcgcgtacgttgcacacagcgggaaatgacaaagaaatggataaaaagatgcatggttttctaaattgtccggcatacagaaaacattcccctgcaaagttatatacgtggtaactcgtaagtgggaatgcggtgttataaaacaaaacacccatgttataacgaccgtcgttttctcggcacttgataataaataaatcgcattcattcaattaataatagatagccagcagcacgcgtatgttatttgagcaatcgcttgcaagaggcacgtccgggaagtccacaccttctacctgaaacgtctactttgacgttatgaacccgtgcaccgaaaagtgttagcgaaaccaagatggttaaactggaatagcttcctcaacacccacgtacactctttccttttgcgtcaagcggggtttcctttccctctagaagtgttgatataaatccttcatgacaccggacaccgcattaagcgcggttcaatcagaatatctatgcggcacccgcttcatgtagtaggccgggggaagatgggacacctttagcacataatacccaaacaacctaatcgtattttaaacagttaacaacggtctatggaagtcgtgaggatacggtttaataattctttaaatatttcttgtttactaccaaatgagacgagaaaatagaataaaaaagtgtcccatctccccccaccctactgtataaaccagaaaaagtggaaatgtccgaaaagagttttatcagcattttagtattttggcgatttaagctttagatacacaaggtgttagtaattgcggaaaggtgttttgttcagttaatctgacgaaaggagttggtttatatttttatttctaccaatatatatatattgctaccaggatttagtaaaaagcgttttatagttaatttaaaagtaaatattttaaacagtaagcaacaaatctgtacgattaattgctccgtaaacgtttactgcatgttcggttaaataaccacctcgcttttcatttttcaatcctactcgttaacgtcgttttgctaattccctttacttttttcaaggccaagttagttactataccaaacgcaatataacataacttaaactgttgtttagctatttaaatacagaaacaaaaataagattaattgaaatagcaaaccaagagaatcagcaacaaaactacacttgttaaaaatacgctatgaaggtaaaaaaaaactaagtaaaaatgtccaatatatttaataaaacgaagctatggtgggtggggaaaccttaagctaaatgctccaggaaaatatgaatcatcgacgcctaagttgccgccttagttgcataactcattgtatagcgagtcacggaaaatgctgcgcgtgtaaatttccgcatggtgtcgctgctgagccaaccggtctcgctcgtttcaaaaagtcgagttttaccgcaaaaaactcttcggttacatctcttatttataaacagcaacccgaggagtcacgctgtaaactgatcgggtcgtgacaaggttcggacaggagaggcagcttcagttataaccgctgaatatcaacggtgaacgttaaccgccatttttaatgaacgttggagttaaaaagttccaagattgagagattaatttaaaagttgtggtttatataaacagggctattgggtaaggctccatagtgagcggtgtcagcaggtgtttcgtaaggcggcgcgtgccaagttctctacttagagcttgtcaaaacacgatctaattactgcatcattagcgcgccattgttcctcgcgaaagttgattgggattatgacgctcctgctttccattgtttaaggggaagatgaactttttaccttcgctcaggctcgactcggtcgtgggcaggtaccggcagaaaacattcgattattgacacgaaggcagtgcgagtgttgtgagggaagtcgtttcggagcgacgtttgtttgcttgcagcgttggcgttcagattctaacttttatatatctcgggcagtgttagtgtaagttaagttacgttgaacacaggacaccgaatccttggtttgattctctata

>KH.L96.43 -1392/-1

atcctgtgccaacctctacgaaaatgccgtaatgtgcaacactgctgttttactgaggcctatgtaagcttgactgaccaggaagagatacacatcaaaatgttatttgctcaaaatagcatacaattaatacgtaatatgagttataaaaattgtaaccttaacatttttttcagtttcgtctttttacgtgtgtatttcgcgggtgattttcggtgctgtagttgtacatacatgttctctgtcgtgtgcctatagttaacagcataggcaattgaacacacggtgcattaaccaaacacgcactgctgctagagattgtctactgcttataagcccactaatgttgcaatgtttttaccttttccctttgcacgctgcccagcatgtcactgaccgcgtgcatgaggtcttaccttgtcggtactccctaagtaagccatgccaccggcatattgctacctttgtataacgactgtgttttaattgcgatttagtagagcggccaagtctttgtaccgaatacacatttcctattgaactttcgatgtgtgatcgtcgttttactgttttatttgctgtcgttttatgcgtgtgttttaattcatattacctgtcgtataatcccttcgtatcaccttgtaatcggcgatacattacttacctatcggcgtaaatgaagcgacgttggtaccagacactgtttttaccaaagcacgctaatcattacgtattgatcaccttattgctggttcatatgggtttgttttcatattgaacttaacgtacagtgctattaatacactgctgccccacctaattgactggcaagcatatcagtcttttagaattacaactatcttcgcctgcagcgatttgctgttctgtgcacattcgtctgattttttattgtgttttaagttttattaaaatacctgtacagaccgcaattaccacacgtgaaactacaacatctgatcgcatcccgtaccgtgttttgtcgccgctttttttcatcaaatcacttttgtgttttttatcttttcatttgtaccattgtgattatttgtattaaaagagatccaacctgtttgctctgtagcattataactttgcattagaacaagccaatgacagccaaactccatttgacaagtgccaaagacaaaaatacaagttctcatctacctatacacgacttagccatgtaatccaaggtaaaacaacttctaacctctcgggaattactgagttggagggacagtaccagcgagtggataataataatccccttgatttacagacgtacccgcgcgaaaacaacaacagcagaaaaaatacgttatcttgctactgcgtttgactggacggtgttgccatttgcttggagaa

>TGFB (KH.C3.724) -3218/+24

agagcggtgtggtttagtgggaaagtcaagttaaatctcagcaaggaggaaagagaatgagacagaaagtatgagaaagagaagacaatggggattatgacatcaaagttaaagtgagaggggagtgtattacgttggtttacatttgtttacgtgagtcatacgttctatacatattggaactacaaaagcatcgcctactgccctactaaatacatgtaggacaagatggatacccttatagtggaacctaatatccattatttttcgaacgtgtgtttaacaattaacaaagcttgataaagttgggagaatgcggttatttaaatctgcaaataatatttgtttttttaccaaattggacgagaaaagagaattaaacgtgtcccatattccccttttgtcttttgcgcaagagtatacccatttaaatgtacaatgggtctcaaaatttaaccttttttttaaatccttcaaaaattaacgtttttgcataacaacgggaaatttaatttttttggttgcttattatctttaccacgaaacgaagcccattgtggcgtaacaaagcctttgttgtgtatataaaccttcattattttctcttcggttgttgcaatgaatacggataattatagttaattgtcataaacaaataaaacgatgttttgtgacaattataatatagaacagtggggtaagatgaataccgttagcacttaatatccaatatttcctaatcgtgtttcaaacaactaacatcgctaagagtcgcggggctacggttacatcaatctgtaaatattctttgtttgctaccaaatggaacgataaaaatcccatcatgtaccagcacggaccgttaggtttaaaattctcgtgttttacaaaacgcagaccgcgtggttcgcgacctgctgcgataactgtggttccgaatgttcgaccaccacagcgcctgaacttgtggcgtcacaatggcgattgtgacgtaacagtggtcttatatcacacgcgaacgcgaattcgcgacctgtagaaaaagttgaaagtcattttcagctaatccctatttatgacgtcacttaactcgttatgacctaagttaaagaaaaattaaaatttcggttaacacaatatcatatatagctcgcctcggtaaaataattatataaccacatcctcgggattctaaaagagcgttctttattgttaaaaacacgattcgaaaatttggaatattgtgtgctagtggtatccatcttactccatagtactatgttgcaaattgttaaaacgcgtttaggaaatacagacattaggtggcaacggtatcccatctttcccctacagtattatgagcgtctagatgtctctgggttaaactacattgttaaacacaaaccccttaacaaggactccactaataacatgggattatcagattcaagtaaagctgtgtttatagtaaccgttgtctcatttccgctgcttttaaaccgggggtggttggtgcatgttaggacaatggttcggaagttggttgttctgattgtcgcttaatctttatctgttagaaaaccaacaccttagttcaggtgtatcacttggcatcgtgggtgagaatttttgcttttcccgtgttaattattgtgacgtcacgatgccgtattgtaaggtaacaatgtacaggaaggcggcagtgttgttcaagaactttttgagtgggtttgacgtaattcacagagtatatttttttattataattttctttgaaaagttaaaagaataaaaaaatcgattgcaaatttacatacgtggcaacgcacgaggtgtaagaaacggaacacccgtgttataacgactgtcattgctccgccactcgaggataaatgaagttacattcatttatattcattcgttcattcattatttaataatacgattgactgctttaaccttgtaaagccctacgtcatggttaattacgtaattacctgacgttgccatattaaggcatgttgtatgtgaattcgcatggggtcggttcccacgattgtgacgtcataaagtgcgcgaaaaggtcgcgcgagcactgatgggaataagaggtgaaacatggccgtctattttatcaactatcaatcgaaagaggtactaaccgattgatattacttgtaaaataagtgactggcatttaaaacgcctacaaattgaaacaattgatttttaagattatttctcgccgattttattcaaagcgcaagcccgcggtttgaggttcgctgtatatcatattgccgcataagcaagcatgagggagatgtgtttacgtgtgtgcttgataattgtaacctgccagtacttacggttcaagtggctgtcggtgttttaaagtaacgtttgtggattcttagatggattgggtttatgtgtaggtgtgtgtgacttttatttaaaggtgacgtaatcatacttgaggtgacgtcacttttatttgaacgtggtgtcataatgcttgaaatgattttatacatgacgtcattatgttagtgacgtcacgcggaaatgtgtggtaaatgcaatagatgtaaacattcgaacgttggataattaaatctaatgaataaaagtaacttaagtttttcattgagtgacgtgcaaggacagttcctaaaaacgggtattctgtttcatacacttcgtgcccacgatttattacgcatgtaactttgcaatcgaggtgtaatgtggaagctaaaagcgaaggaagaaagtaaataagaaataaaaagaaatcagttaaaaaaaacatagttgataaataagttcgaatcaattaaaaaaaagtttaaatttcaaaacgtacacgctgctagtgtcttggacaagtcaatacagaagcgaaaggcgtcgcgtattccagatacatgtcacttacgttcgagaatcgattcgagagagtgtgcggttagaaagaagtaacaactcggtttcatagttgggaggatcatttgtggcagagaaaaagggagaacttcctacgttagtttaaaactccagtgtcgcgattccaggctggttacttctagaaaccgcgtcataataaggattatgacgttcgaagaaataattgatgaaattttgctttgcagattttagatcagtttggtgtttttcattttttgttgataatttgaaaatttgaaaaatctgatatgtggaaaaggaacacaagggcg

>KH.C4.78 -2991/-12

ccatgatcgctgcaagtttaggttcgccgtttcgcactcatgttaccgaaccacctaacctcattacactggtgccatgaatctgtaaagttgtaatttatctgtcaccaaaaatgcccttacgttataggtaaacgggttatttagctataaaagaagtttttgcttgcggccgtcgtaaggaatccttgaccgacgtataataaagtttcttttcattttttaatatgtcatcgtaaaaacaaggatgaaactgtcgtaccacagctaaactcctatttttgccaatacaggtaaaatctcctgaaagtaggactgttttgccaattgatttaggagtcggacctacgcactaaaccgcgcttatatgtgccagcataatacactgcctttcaaagtcgtatcaacagaagcctatccctaaaggtggaattagcaaagcaaatattatctctgacgcggcctggtagcccttcttttacagacggtcaacattgtctttcaaataataataagtaagcgcaaatacgaatcagatggatgacttagtgctgccatggatataaaatattgttctgcataagttgataaatataaagttttggtttctaaaacagaagtaagacttttttaatttttatattacaaagtgttaataatgttcgccgttcagtgtaaatatgggggcggggggaaacgggacaccttcagcagataatatccgaatatcataaataacacagagtaagatcacactttacgtcaactaccaaaagttaaaggacaattagtccctagttatttagtaggttggggggagataggacacgtttaacacattttattcaaatatcttgatcgtgatataaacaactaacaacgttctatgagagtagcggggatacggttttataattatttgaatgtttcttgtttaccaccaaatggaacgagaaaatagaatgaaaacgtgtcccatcttcaccctgctgtcggttttaactttataataaaaactatttttccgaatgaccgaccgtcgcgcatctttactacgacgtcggttaaaaacttttttgctacatagttaacataactatcttcatgtatttcgagataatgagagataatatatcgccatagcttgttcatgaagtaacatgtagacaaacgagactactcaactcccgtaacctcacaaactgcaaacacttatgtcgtttaagtatgtaatcaattgtactgtgactttcggttaatgtaacgcttcgcctatagacctacagcggttaagtaatatctaatttcgggttaaatttgccgaatccaatatattcgtaaacgtgactcccagactgtcgcaagaagtcaatataaatatctcagtggccctgtaataattctaaatcgagttgaatcaatttggttcgcaaccttatcgcttcttattctggataatctaatcactggcatttgtgtacgctgtttgtgggcgaggaactcgctggcctttctataaaaacacaactcacaaagacctggcaagcggcgtcttttgttgctaggtcacattcagaccaccgcgcgcggtatacgtgttgcttttttctagtacagccaaagcgattctcttcaaaatacagtcgagctacacttcaattaaattcactttcaagcgaatctacatcgaaactaatgaattttcaacattaattagtagccaagtttgaaactaataggtcacacttgtattttggttagtttttatccgaacctttttgaagacttttttggctttcaacgataaataatatgccgagagacgttttataatgaatccaataaagaactcaggcttatattagactaccaaccctaacagttcatgaatcatttatataaggataacccgaatgtatacacacctatacgtatttaaaaacagctcaccactttttcagcgtaacatgtttaggcctagtttattacagctttgactaaatttgacttcacctttgattacatgtcttcaaattttggtcgttttcacagtaatagaatttcgataactccaaacgaatcagcaacataaatatgcgggcctcactcgtctggggcacaactcgtatgcaaagtttaaaactacgttcaaatgttaacttgcatccggattgtgacgtcagagaaaaacgtaagttacgttttttgcttgtctatatttgttaaagcgtcacgtttttattcaatttgatgaaaacaatgagcgattaggtttagaaaagcaaacacgtatagataaattttgtttttttgaccaagtcaacgaaagttaaaatagtatatacataaaaaattttgacacgctaaaaaggtataaaacttggatacagaacgtaggataaccgttatacaatgagaaattttattaatgtaacattattgggaaactgtggaaaaagagtttagacacatgactagtacaataatgggaactaccaaatttattttttttactaatgaaaaaatattatcacgaatgaattacgtaaacgagttttctccatacaattaatttcagatcaaaagcggaacggatagtcgtcattgcattaataattcatacacttggttgacttatacccgtcgtcaacatcaataattgatcttaactgtcgcggacataataaattcggttgctcgaattttttaaggtaattgactcccgtgttaaggtatgcgaatgttggctgtttagtgtctgctagcgtggaaataaattctttctcatgtttgctggttttcagtatagtaaattagggagattggaatttgttaaataagctgaatgtgtaaatttagtcagtaaatttaaataggatgtaatgtgtctaatacctaatacgttgtcgattattcagcgactttttaatgatgtaaatttcagacatttattaacacggacaaaaacttagtactga

>KH.C11.360 -1395/-3

gacgtcatcatggtacaaagacagtcaggcgttttatctgtgacaaggatcacacgtcatcagctcgactggttatacaattaaaatcgattcgcagcgtaatgacctacataaacccgcagtgataacacattagctatgaatggattcacgctgtgtttgaatacataaccctagttcaacgtacagacacggcgacggcgtacgattataattcaaactcgtaaaacgttccaacgcgtgtgcgctattccataaaatttctaacttataatcgcattataattgagcacatttcagtgcgccgttgtaactggtcgctaataacaagcgttgttcttatttaaggtcttggtatgtgttcgctactgtgaactctgctcgactgatacttctgactcttcacaacctgttatatacacgcaccacagtcagtgttaaatgtaacacatttcatccctagcgcctcgcgctcgtccaacttcttttcgctttttcatgtcggcgtttgaaagcgtcggcgctcgagttagcaaataaggtctttcttttcacactcggcaccagtccagggtaggagagatactacagaccaataatccgcacgagtatcatcaccttcgctaaacaagttctctaattcggggcatgagacgctggctacggttttagagcgtcttttgtttcaggtgcgccttgcttcgaaggttattgagataacgctgtgcactaatccacaaaacctttaaatctactaaacaaatgtaaatagtttagaatgtctcttttgcgttttggcttgcaacattattttattgttatcaaaacatatggactacaatcacaacaaacaccgtaagccgccgcttattaatccatttttgatgatcagttttcgttctaattcacgaactcaattaatataaatgtttaaaccaaataagtctacattttgcgttgttgtgtaaaatttattctcttcgcagtttcaattaatataatcgtcgtaaatcatatatgcgacacccaaaagaaaatttgtgtcaaccacaatatgaaaaaggacgcatataagaaccttatacacattgcaaatccattatggcctgttgtttttaaagttttaattgctacattatacgatacaaaacatacgcccaaatgcgtattgctacgtacatgccgacctattgccaatatatttagccaactaaatgcatatcgtaaagatttaataacatatcatacggatttgacgtatagtttcttttatatgttacgtaacccttagtacataagacttcactttcaaagactgctatgttgtgtggtttaattctagctccgtcatactgaaattttattttctaaactaaacgaagtttttcttgt

>KH.L141.36 -3401/-1

aaccactgcggttttcattatattaccaaacgcttatttcatacgatttgatcttaaatggaaaggaatgtttgaaaaatagagatttgctctgacgtcattaatttttttgacattttgtcagatttgcttgtgttggtgtattgatcttataatgtatcatggtggagggttagtactaaggataggaaatattaagtttataaataaatctgtaaacctttgaggtgtttctatttatactatatcgcgataaatatttacattttttcctttgtacagtggcccatgtatttaataccttgcaacaattcaagcgagacaaccaaggggcttatgcacgcattgcgtggtacacagttattgcgattacttgctgcaaaaatacaaggtcacagcatcaccacacaaaccactacaaaaaggaatatcttatgcaaatttaaattgttgtgtggctaattgttcaaaacacgattagaaaacagataatatggcatttaggggctaacggtatcccatcttaccccacagactgctgtgtaataatctgtacaaatatgacagaatagcgatttattattttaccaatatacatatttcatattattacaaagcactatagtcatacttagtaccgatgttttattccttgattctaatactcactgtaggggaaatccaggtaatttgttactgccacgaaacttaacacttacatactgacaacttggttcacacgcgtatctgcctatcgaagatctgtttcattattacacggaccatttgttcagcgacaccaaaaccctttgactttcctgagattacaggtttgaaacattgccgttaaatgatctccaccaaccgggtacaaggctcatgcttcgccgccatatagcacgcttcaactattacaacgcccggtgacgttggtcacgtgcctgtctcgcacttggcgcttcatagaggctaggcttcagtgagatactcgattgcgacccggggtatatggggtgcattgcgcgacttcgtgcgtacattcttcctcgttggcggcgaatgaaacttttgtcgtggccatatatgtttaaaatcaacactacagcaataactggtatacgataaaacaatataaatctttatttgttttgctttcaaacaaacgaagatcagattaattaaaacgaagggagggaaattaacagcaatacgaccattgtttaaaacaaggtacgaatatgttgtatacttcaatgaaagtgtcaaataaaacatgtcacagcatatattgtagcaaaaactgtgacgatcaaatatacagagaggaatgacttagataaagttcagttatatctgtaatatcaatgctaatcacataacaaagttgtacgttgtccaccattggtatactcacttacagacaatagttatctcccgaggtttttcgtgcaatttgactgcatttctttttattgttaatcaagtacgaattctgggcagaataacttgtagcaaatacggcatgttgtaatttcaagaaagccgttaaaagacagcctgccccagttctatttagaacgatacattttcgagaacctccttaaccgttaaatatttgtgtattaggctattgctaaaattattttcttacagttttgaaacgaataggttttttatcgtataaaaaatttaatgttaaattatttttcatatttttactgtttaaagttttagcgattattccatgagtcccaacaaacagcgatgatgcgactcgggacgcccacttgcggattgaagaaaccattctcttaaaaatagatatttgcaatgccccacggccgggtctgtcttttgtttctgataccagagaacccatgtcgatatattgctgatgacgtaacatggaatactgttagcacctatattccatattttctacttgtgttttggacaattgaaaacgttttatttacaacgctgttttagagcagtgggaataaggttatataattctgtaaatattcattgtttaacaagtgggacggtaaaagtaaatgaaaacatgtcccattttaccccaaccaatgacatcatttattatcaacactataattatatttactacatatagcgctgtataaagttgatgtatatatatctatatacatcaatataaattaaaacgtaattaagcaaagtataaaaatgttgtttcgtttttgcttttatcaattctacctcctgtgtttattactcagaaattcatgggagaaaattgacgcctctggtttttcccatttcgcacaaactacttctcagaaattatctcaatgagcatgtgactaaaaatctatgttcttataatggtatattttagttgaagtaccccttgggtttagatcactaaattacccgaaaaatttttttttaagtttgtccccaaaaaaactgcaaaattacaatgcggggcaaaatggcgaacggcatagaaaggttttgatatttaaaaccaattaaccctctttattttaattgctgcagataattattatatttaacattctgtcgattgaaaatatgcgggcgtgtacttaatggctataaattttaataaaaagagatatactgagatttggctctttcaattaggtgtttcacgcagactaatatttagttccgttcaattacaacaatattcacgttttatcgcaatcctaaatatgacattacgagacgtaatttaaattaagataatctgatcaaggagttcgaatagccggaggggggaatttactttaacgtaagcaagcctatatatattgttttgccaaactttcaaatttataattagtttagcggatataatgtgaaattggtatgtattaggttggggaaagatgggatcttcatgtttgggtagaacgggctatacgtgcccgctggggtaagacgggaaacttaaactggggtagaatgagacagctgggttatggtgagatgcgacgctgtatagaatgttcgattttcattcccttataatattttgtttgcttgtaaactaagaatatttacagaattatatgaacgtgtatcccaagactctaaaagtctccatctaaccctacaagactttagcgccgcgaatgtaaagacagtcgttaattgtttcagacacgaccaggaattatgggaacttgttgctcacaatatatcttatcccacagtatataggtataagttttaaaggaaccgtc

>C. intestinalis (Type B) C11.360 promoter (rosChr11:7821918..7823549)

ctgaggaaaaaaatacgaataatgacatcatcacggtacaaagacagtcaggcgttttatctgtgacgaggatcacacgtcatcagctcgactggttatacaattaaaatcgattcgcggcgtaatgacctacataaacccgcagtgataacacattagctatgaatggattcacgctgtgtttgaatacataaccctagttcaacgtacagacacggcgacggcgtacgattataattcaaactcgtaaaatgttccaacgcgtgtgcgctattgcataaaattcctaacttataatcgcattatgattgagcacatttcactgcgccgctaaaactggtcgctaacaagtattgttcttatttaagttcttggtatgtgttcgctactgtgagctgtgctcgactgatacttctgactttacacaacctgttatatacacgcaccatagtcagtgttaaatgtaacacatttcatccctggcgcctcgcgctcgtcgaacttcttttcgctttttcatgtcggcgtttgaaagcgtcggcgctcgagttagcaaataaggtctttcttttcacacccggcaccagtccagggtaggagagatactgcagaccaataatccgcacgagttacatcaccttcgctaaacaagttctctaattcggggcatgagacgctggccacggttttagagcgtcttttgtttcaggtgcgccttgcttcgaaggttaatgagataacgctgtgcactaatccacaaaatctttaaatctactaaacaaatgtaaataatttagaatgtctctttcgcgtttttgcttgcaacattattttattgttatcaaaacatatcgactacaatcacaacaaacatcgtaagccgccgtttattatttcattttcgatgatcagttttcgtcctcattcacgaacacaaataatataaatgtttaaaccaaataagtctacattttgcgctgttatgtaaaatttcgtctcttcgctatttccattaatgtaatcttcgtagatcatagagtatagggtggggaagatgggacacatttagcacataatatccaaatatacctaacgtcttttaaacaattaacaacggtgtataagagagtcgtgagaatatgcttttataattctttaaatgttctttgtttaccaaatgggacgagaaaatggattcaaggagtgtcccatctccccccaacctactatataagcgaagacccaaaatgattttgtgttaaaaaaataggtatatggaatttaaaaggacgcatgtattatgaacgtcacacacattacaaaaccattatagtatattgtttttaaagttttaattgctactttatacgatacacaacatacgcgcaaatgcgtattgctacgtacatactgacctattgcaattatagttagccaactaaatgcatatcgtaaagatttaataacatatcatacgtatttgacgtaaatagtttcttctatacgttacgttaccctcagtacataagacttcacattcaaagactgttgtgtagtgtaggttaccttaacttctcggtctgtcatactggaatcttattttctaaactaaacgaagtttttattttaaag

>C. intestinalis (Type B) L141.36 promoter (rosChr7: 1085551..1088515)

ccccactgtattaaacgctttaaaaccgctacggatttcattacattgttaaacgttaattttatatatacggtttgatcttaaatagaaaggaatgttttaaaaatagagaattgttctgacgtcattaatttttgtgacattttgtcagatttgcttgtgttggtatattgatctgctgatgtagttatatatcatgatagcggggtattactaaggatcggaaatattaagtttataaataaatctataaacctttgaggtgtttctatttatacaatatcgcgataaaacctttatattttcccctttgtacagggtccatgctttcaataccttgcaccaattcaagcgagacactcaacggggctaatgcacgcattgcgtggtacatatatattgcgattaattgctgcaaaaatactacatatcatcatcacacaaatcactacaataataatatcttgtgtaaatttaaatagctgggtggggtaagatggtccatgttttgattctctttcccaccgctttggtattgtaaacaaagaagatttacagaattttacaactgtatcctcacgactctaccagagtatggttatttgttgaaaacacgatcagaaaatatggcctttaggggcttacggtaacccatcttacctcgcagactgtataataatctgtacaaatatgatagaacaccgatttataattttaccaatatgcatattttatattattaagtacaaagcaatataatcatacctagtatccgatgttttattccttgattataatactcactgtaggggaaatccagttaatttgttactgccacgaaacttaacacttatacactgacaacttaattcacacacttatctgcctaacgaagatctgtttcattattacacggaccatttgttcagcgggaccgaaacctttgactcctgagattacaggtttgaaacattgcagtttaaacgaactccaccacccgggtatatggctcatgcttcaccgccatacagcacgcttcaactattacaacgcccggtgacgttggtcacgtgcctgtctcacttagcccttaataccggctacgcttctgcgagatacttgtttgcggcccgggtacattggatgtattgagcgagttcgtgtatacattcttcgttgttggcggcgaatgaaacttttgtcgtggccatatattatgtttaaatgtaaacactacagcaataactggtacggtaaaacataaaatcataatttttttatctcttcgaacaaacgaattgtttaaagttttagcacttattttatttagtcccgacatacagcgatgatgcgaatcgttacgcccacttgcggagttgaagaaaccattctcttaaaaatggatatttgcaaagccccacggccgggtctgccttttgtttctgacacccgagaacctatgtcgatatagcgctgatggcataagatgggatacctttagcacctatatcccatatttcctacatgtgtcttgaacaactaataaccattcgttttatttacaacgcgttttaaagtagtgaggataaggttatataattatgtaaatattcattgtttaacaagtgggacgataaaagcgaatgaaaacatgtcccatcttagcccaacctatgacgtcatttattatcattactatatatttactacatatagcgctgtataaagtttatacacacacatatataaaataaaacgtaattaagcaaagtataaaaatttttgttggttttctttccatcaattctacctcctgtgttcattacttggaaattcatcggagataattgacgccgctggttttttccatttcgcacaagccacttctcataaactatctccaagagtctgggactgaattttgaatttatttttcgtacattatcttgtttttatattggtatatttttttagaattatcctaggtgtagattattgaattacaaaaaaaaacatttcaaagtttgcccaaaaaaacctgcaaaattacaatgtgggccaaaatggcaaacggcattgaaagtttccatattaatattattctgggctttgccgcgctgtttagagtcacgagaatactgtggaaattaccgcaacaatttcatcaaatggacacacttaaactgttataaaatcatggcaaaaagcaattaactctctttattttaactaaataccatgctgctgataattattatatttaataatctgtcgattaccaatacgggtgtgtctttactcgctataactattaatataaaaagagatactgggatttagctctttcaattagggtgtttcacgccgactaatatttagttccgttcaattacaacgatcgttcgtattttatcgcaatccaaaatatgacatagcgagacgtaattttaattaagataatctgattaaggagttcgaatagccggaggggggaatttacttaaatgtaagcaagcctatatatattgttttgtcaaactttcaaatttataagtagtttagcggatataaagtgaaagtaatatgtattaggttggggtacgatgggatcttcatgtttgagcaggacgggctatgtgcaagttggactaagacgggaaacttaaattggggtagaatgagacagctgggttagggtaagatgcgacactgtacgatggtccattttcattcttatttatcatttcatttccttgtgaacaaagaatatttacagaattatataaccgtatccgcaaaactctaaaagttttcatctaaccctacggtattttaggcagcggccgcgactgtaaaagaccgttgttaattgtttcagacacgatcagaaaatatgggattttgtttctcacaatatatcttatcccacagtactataggtataagtttcaaagaaatcggtatg

>Gnrh1 (KH.S1051.1) -3787/+42 ATG start codon

(Based on Kusakabe et al. 2012)

gaaaacatgggtttcccaaattcgatcaactttgctaaagattttatgcgtactggtgttggatgagattctgtataaatatttaaatgtatcttattgttaaaaaagtcataactgtgtgaaatgacgtttagcaagtcgtgatgaaagcgctcgcaaccatagattataaacccacgggtaacatggttcgatgtttaaggctgctactaatgtctgcaattatcctagcgttgaaaatagacgtcattcagccactttttgaaacaattactttgtgtgcgtttgtgttgaaacgaccactttgtgcgtttatacacaatatatagtaggggggagaagacgacacatctttagcacatattatacaaatatcctgactggttttaaacaatcaaaaacggtctataggagtcgtgaggataagattttttgattctttgaatgttctttgtttatactaccaattaggcgaaaaaatagaatgaaaaagtgccctatcttcctgttttccccccaccctgctataaatatcttccgcatatattaaataccgtttttcttgtattcggccaggtgcaagaaaaaaacaacaaaatttaataatcacgtagagtcgtggacaattcattttcgtgaaaaacatattcatctcaaaatagttttctgacgtgttgaaattattgaatatggattatgacgtcatacagccatttgaaacgctcccctggtgcagattgatttcgaattttgcgtcactgaaaataattacctcccgagcctacgtcatgaacacgtgactagttgattaggggaatacgtcatgaagcaaacgaaataaatcccaatccttttctttattttcgtttgtcaaaattgaaacgaacgttgtttaaaaaaaagaaaatatattcacatagtgatttattatagaaattattatttttagttaatttaaaattgagttttttgtggaaaaagattttattcatttttagaaaattgtatgaactgaagaaatgggcaacataaaatatgtggttaattatgcttaggtattatctcatggaaatcttataattggaattgctaaaaatagaaatgttatacaccatactactagagaattatcgatcgcccaaaatcgatttctatgagtgctgctgctgtaaattggaccgttgaggttttttaaaaataatttagttaatatttttgcactgttaaaatagttgaaccgttgttttatttccaattggactgcttttaaaagtttttttttaacaaaaaacagcacaatgaaatataaagtagggtggggggagatgggacacttattcatttcttttttcgttccatttggtggtaaacaaaaaagatttaaaagtttataaaatcctattcttctgggctatcacactactcccatggaccgtggttaattgtttaaaacaagatcaagatatttggatatcatgtactataaaggtgccccagcgtcccccacagtaatatatatggtttggttttaaaaatcatttttgtgaatacgaaacaacttttcttaattggtcaaaaggaaataataaaatcttaattattaataattcctataatgattttacaccgtttttttgggggctattatacaataacagaaggcaagcgtatgtgtataattacttatttaaaatcatagtttccactaatctatactttattgtaaaaaagtcaaaaaatcttggatcaatttcactttaagatgatattccaattggcattattttttaatgcgactacgattgcaccaggttttagcctttccgtttactggttttggaagaaattctcggtgccaattacgaaagccagctgatccaattattggtttttagatcaatatcgcggaaaacgggatcctaattggatggctcttagaatacccttaacatgtgatttaattaaatgaaacacttgtggatgttgtttgccggcgttttaaattgtgacgtcattagcaggctcattctttgcggctgtgacgtcattaacaccacttgggagaaaacaacgataataatttcctatttgctatggtcgaatatatattacagcagttctacaagagaaaaaatatgtaaatctatttttcatttaaaaaagcattgctatagactaaaggtatatagcaggttgggaaaaatgggacatctttagcctaaaatatcctgatcgcgttttaaacaattaacaacggactatgaatgttgtaaggacacggttttataattccttaaatattcttcgtttaccaccaaattggacaaaaaaatatgtcccatccttccctaccgtacgaatatacatatttgtggtgtataggccgtaggataactctttttaagcgttggtcgtttacgaaataaacgaattacgtatatgtatataagaaataccctcgacaaaattaccattttaaaatctgtgtaaaataagtaaaatgaaaatggcgtgccactgtttttccttcacccatcatcgttactgcgacttgtcaaccatgtggttacgaaacccgaataaactttgaaaaatacgttttcgcatacgttaaaaatatcgtttcgttcacttgtttgcgtggtaaaagctaaactgtcataaaatatccttttcgcattcaatcggttgtcataaattgctaaattatgtaacggttattaaaaattgttaactcattaaaaacataaggtaacaccatatatacgttccgatagctattataactacagaatcaattattaatttaaagcaggtgcaatttaatccgttcgtcatttacgaaatccggcgtttcggctgcttacattaaaaattaattaaacggttacgatgttaccttttgttaaaataaattgtgctggtaaaaacggtagaacaattttcgtttttttctacggtaatgacagtgaactaacgtttatgatgaatatatacgtaggtgtatcagaattaaactacaaacgtatatcacttatacaatgcacaaaagtttatttatatgtcatacctaaaaataatcaacagaaatgtgtaatattttaatgaaataacaaactccttttatttttgatctaaataccggggacaaataaataccagattatgcatatataagctcgtacgaaataagtattggcgtgaacagaaaaaagactttttaaaataacgattgcttctcattttaccaaaaggaaatactttttgaggcatgtattccatcatgacccagttggttgtcgagtgcaattagtatcagctatttcgaagcggccaggcagaagaaaagtcctttcggcagcgttgttattcgcaggcaaatttgattagcttgacactgctgattcgtcttagagttgtgttgcattataaaaggcagcgagtttcacaaaccgacaccattcgtttgggtttcagcgagaacgacagaaacagaaaagaaaaaaaagcctaacgtctttttgtcacggcctgaattcagacattgggtgaatatttctagtcttatgttctgcagatttagtttaaaattttcagatttgtcagaaaggtaatgttggaattagtgtgctggccaagaaagatatactgttttaatattaaatcgtaaaatattactaaagcaaccagaaaagtgtagcgtcataactatttaaagattaaatatttaaatatttgtgcttaccaaattttttaactctagaaaacgataatcATGGGTTTTATTAAATACTTGTACCTATTCGCGTGTATAAGT

>KH.C14.116 -3252/-1

tgtggacaactctgcctctggtggcagctctgcagacagagatgagttaatttcgataactggagacagtgggaacgttttaatgctggatcaattaactagacaaactggagccactgtagctcctatgctttgcccgaacccgtgttaattttacgagattatgttaaagaattatacgttggtgataaacagcttattatgaaatgaataaaatttaagaaattatctgttcagtttctggaaaaatttatacaatgagaatattattgggataacactgtgcagattagtgagtgaatataaagacagtaaggaattttttgaaataattcatgatgaaacctttactatgagacacttagccgtaatggatatagctggatttattctgttaatgaaacggcatgcccgtttatttgactccttcttatcaaagaaattgaataaggcaaatttagatttctgtttatcacacgtacggcgttctatgtatagtaggatgggggaagatgggatacttttcattctatttttccgtcccagtattaaacagtgaacattcgaagaattctaaaactgtacctccacgactcccatagaccgttgtttattgtttaaaacacgatcaagatatttggacattaatgcgctaaaggtgtcccgttttccccacagtattatattttcttacacaggttgtatttttcttgtggtggtttgaaaatacggaatttctatttcccgcagataattttacccgacactaattgtagttattttattcgcgtataaatgtgattattataacctctactaacacgaagacagcacattaattgcaggtataatgctaacatgacacaacttgcatacgtgcaccaatacagcgcttatatgcacgttaactaatgcctttataaacgcacaaaagggcgatctttttatagtctagacgttcggaaatgcctgcaacaagagttagaaaatgtaaagaattacaatcacgtaaatttcccggtaatctgtatatgttgcataaacaaggtatatgttattaatccgtgccttacgtcacgataaaacgtcgaattttaagaatatttattattttttaaagaatactgcgtgactggtaaacgcgaattttgacagtttaaaaataaaccaccggaaaaagtttatctatctttgtctgccgggtaattcggaagcttcacccaaacatataaaaggacgggtgtaaaggtattatagacgtatttaaattgtcaagaactggtattcgtacaaaatgaaaggctcaatgttgatttgttttatcgttgccagcacttcatactttgaactaaccaaaggtttgtagttttattttaaaataaatatttttatgttcattttttccgtaatatacaggtttatcctgttggacttgtattaacgcaagaagcaatgcggagtgtcaggctacaggtcatctgcaacaatgccgatttacccaagtaagtttttcgttttgtaactcatacagaaataacaatggtataaacaatttgggtcaagatggttcatgttttcatacacttttatcggcacatttggtacaacagataatatttaccgaattattttgcggtagcattcgactcttttagaccctggttaattgttacaaatacggttacaaaaaaggagtttttttgtgctaaccgtatccatcttaccccacagtactatatatttacgattataatgtgcctacttattttttagagagcttgtcaaactcacatacgaaccgatccaatgggtattcgcatcacgaaggaatgtaaacaagtacaagcgtgcaccaacaactttttgcaggtaaattttaagggtgtgtaaaatgaatgcaagttataactaatatataaacaacattataaaaagtatgtgttgctgctacaatcatgtgttaaaaaattttttaataaaaagttaatatgataaagtatgaagtggtatttttctccgttaaataaacaatccgtttatattcaaattttttatttttaaaaatgttgtgaaaagttctgtactaatggcatattataaaataaaacaacagaatccgagaccagcctggtacccgagtcaatgcaacgataatgtacaaggatccgtatgtcgatgctgctgtgatttcgacaactgtaactttgagtctgtagcatgcccgggaagtgagttggttcaagttatatattgtaacgtgtatgtggaggttcggtatgaataaagaaatgaacgtaactgatttatattggttctctggagcaacgacggtcgttataacacagttgctttattccaagcaccttatattgtgcccgttaacgagttaccatacgtatttaacttatgggttttttttataaacagttgatattaatttaaaataagttatgcgtttagttgtgttatgcccaaacacaccaccataatgtcggcaacgtttcgaatacgtatccatcagttaaaaagcgagagctttaactacttcggcatgacgtcaacaactaaatcttaaattcaaatttttctacaggtcgcaccacaactactctagcaccaacaacgacaacaacaacccgtaagtagaagatataataaaacagttggcataggggtaaagttgtatacgagttttcactctattttacccccccccccaattcggaagtaaacaaagaaagttcaaagaattatatataactagtagagccccaaccgaatcaaaaaaggccttattgattttaataaaacacaatcaggaaatttgggaaattattatgctaaatgtaccatcctaccccatagttggtactatatgtttgtaaagtaactgcctactgtttcaatgcgtagtttgaagcacaagactacacctatataacgctccagggctcttagtaataaaacaacgcacatagttctctgtatctgtaagtccaaaagttaacaggaataatttgaaggattaccataaacataaagtaatccctaatcatataatgaccactcttattacaaaataaccttaattactaagcgattattacattattaaattatagaaccgctcacgcaattagtggacgtgttaaacgttggaccggaggagccaaaaacctgtgacaagataacgttacgaaacggattcgtcgcttgcactgatgagaataagcacgactcgct

>Emx intron 1

gctggtgagtgattgtgacgtaacaatattaattttggcgcaaactttattaaacttagatcataacatttgtttatatactccttaaacatgctgttgctgaagatttcttcgtgtggtttttggccgtttgaaatcacattcacgaaaactgccaaaacataccttttaacctttaatatcttgtttaaaatagattttttttgaaacctaaatctacattaaccgtttttataatatcgctgctaaaatgtttcgcctaaaagtttccgatgtttttgtcaaaacatcagccgccatttttaaatttacatcgggaaaatcggcgggaattaaacgcctcagcccccgcgttgggtttctaatttcccccaatgtctcacagcacaggatccacgtatttaatgcacttgttaataagtaggtaggttaactattctaaacttcattcccacgaaccagggtacttgaaccgccacattaactcgctatagtgaatgttaatccatgaatgaaataaatacaagttttgtaccaagacatgagtgctaaaactacattacgataaaacgttaaactattatctaacgttgcaaagcaggctttaaactgtaataaatccggtgaattaggctctgatgatctaatgttaatgaagttggacgattcccgtaaccagttaaagttagaatcaagcgctacgaaatacttagcccaacaaatattttcctataacagatatatttcacctaaatatttacaagaatttatctttcgataaaaacttgaactaaattttattaaaaccaacatttcgtctgccgtacgacggaaacttctttagaattatggtgaaaatggttgataaacggacagttgaagttcgtttcccaccaacttcggtcgttggtcagcgtttcaaaatcttcaatgtcgatcatcgatctatttattcgttagtttaaacgcgacaaatgtttctaaagcttgatgtatataattgtagcgcgacttggtataaatatcttatcaggataagatggtccaggttttcattttattttatcgtcctctttataaaccgtatccctgcgactttaaaacagcgttggtaccggtctatctatacactttaccgcaaagtactttattttaatagtgagttatgtatacatacactgtgtgttgtagatatttcataaatttaatctttttagaagcatttactttaaaatcaccagttcgaaatgttcaaaaaacaaaacaagcacgggatcagaatatgggatattatgttctaacaaagtaacagtatgcacagtattataatgttcgtggcaaaacacttcaactgaatgccgcaaccgttaacttctaaacacagatctcaccctttgatcttcgtctgcaaattaacgataatcgaccctgagataaacttaaaaacaaacataacctcgtatgtaacgatccattaatcaccccctaccctataacacggcaccgaatccttggtaatacgaactgctgataagaatcacaaccgacataacgcccgcccataaattacagtaattgatcataatttataaaacgttaaaataaataattttgttttcttaacaatacaaaacgttatattttgttcgttttataatgtttctatgttagagtgtaacgatggagatattttttgcaaataatatttacaataaataaagaagtaaaccgcgaaaaaagcgtgaagcgccgtttcaaagcggatgtgacgtcacaatgaatcattttaattctacaaacgttcaaggttaaaatcgcgatcgtttgaagattaacaccgggagaaaatatcatttcttctatctatattattgcgtcacaaaacgcattatcgcgtcataaacacatttgttgcggcgacaatcaacaaacccttaggctgtccggtgtgagagaacaaaccgcgtgtgtgacgtcacaatacccatggtttccgacacaaaggaaatgttctacaatacacgacaccatgttccatatgtttaattaattaattaaaaccgtttaattgttgggtaaattaacaatgaaacaaatttatcaatgctagcattgtgacgttggtttctaccattattacgtttgtataatcgggtggggcaagatgggacatgttttcataacatttttcgtcccatttggtagcaaacaaagattatttataaggatatataaccgtatccattttacaccaaactacgatcactttttgaaaattaaacatatttaaaatgtaaattctaacaccctatgttacgtcataaacgacctgaactaaccccct

>Ascl.a (KH.L9.13) -2402/-1[exon2] exons

gagcgactcaacatgaaccaggtgaggacaccttgtggttggcattcatagatatcttgttatcaatgtttttaagataaatatcaacaataataccaacaatttgttttaccttatatatatattcagatatattttatttttattcaaccaacattaacccccccccctccaggttgccttctctgacccaaccatcagcggcaaatctttcgcaaccaaatatttcaaaactccccatttccttacctacaattgctccacctagtgagtgaggtcagtggcaacacccccaccacgcccacttttattttcaactttttaagttttcactttttgtgtttgtgacatactgtgttacctattagcagaaaataaatttatttgaaaacatccatccgctagatgacgctgaagagtcgcttaaatattttatttgtttgaaaatttcctaccgctagatggtgctgcagggtaaaacaaaatggcggagtattggcgcgaaatcgcgattgtaaaacatggctattgtgacgtcacagtcgctatttttcgttaataatcaacttgattgttgacctggcgttttctccgctggggggcgacacagttatctcttgtatgtgacgtcatcactacgtcataatggcaattgtcatttaatatttaccaacaatacctagacaatccaactgttccatcatttatgacgtaacaatataccattgttgactcataatggacgtttatcgcgtcgataaacaatggaatcgaagccacgcccaagatttgttcttacacgcgctccacgcgactattatgacatcacaaacaagccattgtaaatcgcgttatcaggaaatgtttgctttcctaatttaaacatagaaatattataactacggtcacgtggtctaatatacgtcacaattacgattaacaaggtaacaatcgattgtttatgacgtcacatatggaatgttgtgtaacaatgccccttatcttatattgttcgtttattggacaaacaatggcgatgtctttggcgccgattttatcgaatccatgtttcgggacaattagctactgtgatgtcaccattacacttggtttgtgacgtcacataggattgtgacgtAATAGGTCCGTGTCGAATTGAAATTCAAGATTCTTAGGTATTGTTACGTCACATATGTCGAGAACAAGCGTATCATTTGATCACGTGGTAATCGAACCCTCTGTGACGTCACAAGTGATATTCAACGTGGGCGGTCAAATGTTCGAGAATCGAAATAGTATCAACTTTTAATCCGCTTGGAGGCCAGATTAGTGACGTTTGTTTTAGAACGCTGTAAACGCGATTGACGCGACGATGATCCTTTTACTCGGGATTGATGGAACAACCGCTAGCGTCGCTGCCGGGACGAATCGCTTTTACGCGAATTAATATTCGACAAAATATCGATTTGTTTAAGCGCGAACAATCGCTATTGTGACGAAGCACGAAAATCGCTGCGACCGCGGCAAATATTTCCCTCTGTGACGTCACATAGACCTCTATAAGCGAATTTTAGGCGAAAATAATTTTTGTTGAAATTGAATTAAAACCCGATGTCATGCGTGTGGATTGCACCTAAGGGCACCTGGCAGGCGTATGCGGTCACGTGGGTGACACGTCACGTGATTATGACGTTTAATATCACGATGAAATGATGTTTCGATGTTGTATATTCGACTTTCAACTgtgaatatgatatcataatgggggttgtgttgtggataagtggcgttagatcggatgcccccagggtcaatacattcaacattaaacttggcatggagatcgtataatccggtttgacaacaaacaacataaacatcttataaactccggaacttgaattgttttcgcgttcaaatgaatcccgtttgttacgtcacaaacatacggaacgatacaattaaaaacttaataaaaagtatttacgtcacaaataattattgtgtcatgctcgaaaccgttatgtgggtgttgtgacgtcataagcgcaaccggaagtgccctacgtcatcaagaatcggtgtttttttgcaaattgggaatcttccatcgaagttttgaggttgtgtggtaagatgaggagattaaataaaagagaataagaagagcaaagagtgagaagggcgcgtggccgaggtttctcattttgctcctgaccaatcagcgtcgaggagtggtttggtataaattgacggcacgcgacactcttgtgccagccccacttctcacgtcataatgaaagttattctcgcgttgtgacgtagggttttgtgatgtaatatttgagttgatgacgtcataataacttaaaattcttag

GTTTTGTTAGTTTTGAACAAAA

>Myt1 -3271/+39 (Based on Tolkin and Christiaen 2016) start codon

tttttcgtgtcgcaagactgttaaggagcaggaccgcgttcgttcaccagatgccgagtgttgattttgctgaactgtcaacaccgacataactcggagcaggatcaatagttcgacttttacggtaaaaagctgctcgccatgctatgtgcctttaaactatgaattataaacctcggtttcgcgtgtcgcctcagttacggagaaaccaaagaaacgctttaaagaagctgctacaacctggttaaaattttctttaacattcaataaaaatgaacaaaaatccatatatatgtgaatacagtgacgcagcaatctttcaaatttcctttaaagtaactgctacgaaattggtttttttaaatgggttattgtatgatggatggttttgcttcgcgtgtacttctgggtgattgtacacaaacatgtaataaatcacactctaattattctaggagggtatttcgccatcagcggttattaaagcagctgctgttaattaagatatagtttgccaaagctgaaatataactccagcaaagtgccatcatgcgcaatgccacctgcagacatctatagtcaggaaatgcagtttcgttttcttccaaatcgagagtataacgtaaagatatttttaacccaagtgtggggttaatataaaattaatatttttaatggaaaaacattaacatttttaattcttctttaagttgttgttgctttaagaaacggagtactttcaggtgatgttaagagcgatttgctgtcgagtacaagtgcattttcagcatctgttaccgggccacaggggtgcttcatctgcactaaagctgtagagatagtacagagttggtcgcacgatttagttgttgggggtagttggcaagcactttgctatctttagtccccatatcgattgacacatcgattacttttgcatggagataaacatgctagcaacatctgtttcgcctatcaagtgattctgtcgtcataacacccaacgattaaattataaacgagtcccaaaggaaagcaactttcaagaatttatttttgacaagcgttatgctggtcgtattataagcggaaaccttttgtttgtttgttttatattgaactgtatgcgcatcatgtgtaggctctagtgtatgaataagctacctaaaaaacaaaatacaccaaacaattgacgactgaataattaaaagacctcgctggttcgtggaatattgcattacaagtaacatgaagaaaaacgacaaaaaatcaatcatttctatgaaactcgttttaagcaatttccattactcatccgaattacaaaacatcgcctttcttcaacatgtgcatatgacattataagccgctgtcgttagtaaaaccatgatgtacacatgatccgactgacctgatttgcagttctatcgccatcgcagacatctgctgggacatctgcacttgcggacttgtttgttgtctgatggcaaatccatcactaaaatttgagtaactgcattaatgggacacaaactattttgttcacttaatccgtaatgaaggttaattattgcgctaagtaatcgtcgcaaaggaattgctgtggaaactaaatctttataatacgtctttccgcatatgaaaatcaaattgaaagaaatttaattattcgcgggcatgtgctggaaactacataacgtgttccgcatgataacaaaacgctgttcgatcgcctgtttaccgaacggtaaggatttgatcactggtttgtcaaattgactgtcataacgggcgaacgacttttaatcgctcgcccctgctggccgccattttaaattcgctaacaatggcgaccactactttcgctctatgtaatcaggaaaggtaaggcggaaacaaatcaacgacacgttcacctgtgcatttaatgcggattaaatttcggcggtaaagctccgatttccgacaactttgcactgttgatgaaaacaagtgcacccgtccagaagtattaagttcactgccaacagttctcgtaattgcttccgcaatgggcaaaggatttagtggcaacgggccttgcaagacaatgtcagaccacatggacctatttagtttctgattaacaagcggcctacagcgaattttcgctcgcctttcgttccctggcacctgcttctatcgttcgcttgcacgtgccgacaaattagtctatactagcctacttttttttaattgactgcgccatcatttgaaacattggcgaaaacttttggaagtccagttacacaactgccttttaaatttaaaattcaaaaaatagtttatatagagcttaattgttttacgatttatttttctgctaaataacattacacaaattggccgaaataattttttccaaagtacttaatgtgctgtaatttgggggcagttgtgtaactggacacaactgcccccaaatttaaaaaaagacatttaaaacccaaaaactttattaaccaaaactcgacaaaaaaagcatgtaaggtaaaaatcaaaatcttagataaaaaagtcttttcatccgggcgcgtggctatttgttgttttgtcataaatttaaagaatcagcgacaacgagaaaaagaaaaaggcgccgcagtttgccagacagatggcaggcataagaccgaagtagtgggacttcgagttgtcagaaaagtcgctggaaccgccaagagcagtcgcgccaagcaggtggtaggtagcgcgcttcgcagatgccgcaaagttttttgatcagcaaagacctatcagttgttgtggcgagaagttatattttacgagcacgggtttggtctgcatgtttcgtaaagtaagccatgctaaacttttgttttggacctaaaatgtaaggttttgtacaagttggaaggaaaatgcgaaagcgttttaaaacgtttgttaaaatatcgttaattggctaaacgaattttacgcggttcaagtttaagaatatatttttggagaatttatttgaattgcgtttagcaactacactggcagcgtagcatagatataagttgactccattttggtttagcttccaatctatgataccgatctcgaaaacaagatttaaatcgtagatagaaacttcagtgtggccgcgttcggaaccaaagtaaaatataatttttattggtaggttttaaggatagcactgtagaaacaatttgtaccaaacgaatgagaaaactttttgcttgaaaacgagataaaagcgacaatggatacagcggtccaccactctgtaggtgcacacg

>Astl-related (KH.C9.850) -1043/-3

tttataactgacatcgaatacaggaaaactgtccaaacagaaattataaacacaaaactactttgcaaatctttctttatcttcgtgtagcacacaatgcattattacattccccccatcaaacttttcgggaatgctttctctgggactttcttctccttcttcagctggcattttgaaaaaaaaaacaaaataaattaacactttgttaacgaaacgcaatggtaaaaatacatcagcaagaaaaagtgaaatgttttttttttgtaaatgtttatcgggacttcccctacgcgttcggttgctacgatccataaaaatgtaacacaaactttttattccccgatttcggtaagactaaaaataaaccctactttgtggcattatagctgtaagtgtatatttttaacctaatagtagtacaggccaaacaagataatgttttttatcggtttttattcactaaaattcctttatgagcgttaagtgcgaccagcgttcatcagagtccgtatatgtttatatagtagggtggggggagatgggacacctttagcacataatatccaaatatcctgatcgtattttaaacaattaacaacgggctacgggagtcgtgagcatacggtttaataattccttgaatgttctttatttactaccaaatgaaacgagaaaaaagtgtcccatcttctcccatcctactgtatacagactctggtttatacattctaattgcgacgattaggcagcgtatattagtacggcagactacatcaccacaccgtattttgtcattagacactgacaaagcactaattggcctattatctgctattaaataggttataccgacatgcagtaaatggattgcgatcggattaagtacgcacgaatatatacatatcttcctaccaggttgttatttcatagaccccgtatcagattcaaacggtaacttctgactacaagctgcatatttgtaccagtctatggtaaaacgaattttaggaaaaccagcaacagcacgctttcaac

>Villin (KH.C9.512) -721/-1

ggtttaaataagaggtgctgtttaaggtttttaacaaatgaaacgtcatcggtttttaacaaatgatacctcacgcaagatttaaatgtaaactggttaaacatgaggtttactatggtttgatgtaattttgtacttttcttattgttgaaactgtgcagctagtaacgtgcttattattaggatacctattaccagtaaaggttcggttttgtaatttttttaatgtaggtgttcaaagggtggcgctagagcccagtgtaacgtattgacataaaacctttttctcaactctcttgtcaagggttgttagaaaacgttcgagattagacagcgtaaaagcatattcagcgtagatagtttgacaatttctcgaattagtttcgtttcaaagaaactttaagtagttgtaagaaacatattaagtgctatataagtggtcaatcacatttagacagcatgtatatcactacagtgcgacgatatataagctattatgtaaacatcactttgtgaaagctgctaattcacgtcagtaaaacatatatcaacaatcatttaatcaggcgttaccgtacgatttaatagcagttgtaggtacatgtgaattaagtgtacctgataaccgattgcgttgataatttgtatgttttatgtttaacaggtttagacttacagattgttggttgttttatctgagtagaagtatcagaaaagagtc

>Sp6/7/8 coding sequence

atgagtgctcaaacgtcgtctttattgcaagccaatcctccaactggacctggatccgcactggggggcgccatattagcccatcatcacgccagtgcattcgtcagcaagatgcaacatgttcacgcaactccaactgaccaccccacatctcctataggaggaccagattccggttcgcctgctgagagtatttgttcaactacgtcatcaccataccacgtgacccctcctccgtcatcaccacctgtgatgacgtatccacattccatgacgtcatcaagctctgtgctcggatcttccgtaacgtcacgtggctttgttagcaacagcccctacatgcgatcctacgacccctggtaccccaagcccgtccctactgacccaattaacccaagttcgtcaccctacgtcactgctacgtcaccctgccccaccaacgtcatcgctacgtcaccagctacgtcaccaaactacgtcacaacaggaccaacctaccaccccaacccatgggaggcggctttgcattcaacaaacacttggttggaaatgcaatccttgcaacatcatcataaccagatggccgcttacacgcccgaatatcacccgaccttccccgcatccactttaaccccattagctacaaatacttcacttcttacatcggcagcagcccaacatcattccccctacgaaacatttaaaccagttctcccgagcgccgcgtacacagacccgacatcgagtctcccaccgatgttggcggcacctacactgcccccagccccgcggaattctcgccgatactccggcagatccaactgtaactgcccaaactgccaggaagctgagcgcataggaccagcagccgccgcattcaaacccaaggtccacagttgtcatatacctggctgtgggaaagtttacaataagacttcccatcttaaagcccatctcagatggcatacaggcgaaaagagattcgcttgcccggtttgcaacaaaagattccaacgaagcgaccaccttagcaaacacgttaaagcacatacctccaacggtgatacgtcatcgaatacgtcatcaatcataacgtcatcccagcaggacacaggagagatcgtcaaccagaagatgcacgtcacatccgctcgtgacgtcaaaccaaaaatagcaaagttggtcaagtga

>Islet coding sequence

atgaacgaatcattcgccattttcggtcaggacgagaaactgggagcaaagatggcggaccctcatggaggctgcgtgtcggcctcgcttaattgcgacccctgccgcatccctctatgcgtaggatgtggctcccctatacacgaccaatacatactacgtgtcgctcccaaccttgagtggcatgctggttgcttaaaatgcgctgattgtggtcaatacttggacgagacgtgcacgtgttttgtaagggatgggaaaacgtattgtaaacgggattacacaaggttattcggaacaaaatgcaacaagtgtggtctctgcttcagcaagaacgactttgtaatgcgagccagagacaagatatatcacatacaatgtttcaagtgtgtggcttgtagcaggcagttaataccaggggacgaattcgctctaagagacgacggtttgttctgcaaagctgaccatgaagttgcaacatccggggacatgatggttcacgacggccacatgattccgggaattccccaaaccccgaaccctcaaggggtcatcagccctcaaatggggggcgagagggtgatcagccatcggtcgggtgggcacagtggggggcaaaggcgaagcaaagatgcgaagacgacacgagttaggacagtgctcaacgagaaacaactccacacgcttagaacttgttacgcggccaactgtcgacctgacgctctaatgaaagaacaactcacagaaatgactggactatcttcgcgtgtaataagggtttggttccaaaacaaaagatgcaaagacaagaaacgttccatagctttgaaacaaatccaggagcagcaagcgaagcaacagcataataacgaacagggtaataacgtccagggtttatcagggatgaacggagtccccatggttgcatcagaacctgtaagaaacgacaactcagttagtgtagctcctgttgaagtgaggaactaccagcaaccagcttggaaagcactcagcgactttgctcttcaaagcgaaatagaacaaccggccttccaacaactgatgaacaacttttccgatcaaggtcaaggctcgatatccgattcgtccgaaatcagtagcattccttcagtttcctcagccagcatggattcgaccacatgttccaccccgcacacagtggagtcaactgtaccggcgtgtagctaa

>SUH-DBM (Originally based on Hudson and Yasuo 2006)

atgcgaccctcactagcactgtatcacccccaccacctaccagctcatggccaagttcagagtcatcagcatagagaagatgctgctgctacgtccagtagggatgttaatggtggattatctgtaacagaatctgccattgcttcatttaggtcgctccgagaaaaatatccgccgaaaaaattgacaagggatgcgatgcgacgatatttaaaagatccaaacgaccaaaccctgattgtgcttcacgcaaaagttgcgcagaaatcctacggaaacgaaaaacgtttcttctgcccaccaccatgtatgtacctgctgggcaacggatggaagaggaagcagcagatccttgaagaggaggagggatcctcggaggcaggacagttgcacgctttcatcgggatcggaagtagcgagcaagagatgcagcaacttcatctggacggaaagaacttctgcacggccaagacattgtacatatcggatacagataaaaggaagcatttcatgctcaacgtaaagatgtttttcggaggcggaggagcagatgttggccagttcagcagcaagaggataaaagtcatcagcaagccatccaagaaaaaacaatcactaaaaaacgcagacctttgcatagcatccggcaccaaagtagcgcttttcaacgaactcgaatcacagacagtgagcacagagtccctgcacgttgagaaaggaaactttcatgccagctcaattcagtgggggtgcttcgccattcacttattggacgacgacgagtcagaatccgaagagttttcagtggtggacggctacatacattacggacaaaccgtaaaactcgtttgttcgaacactggaatggcgctgccgaggctcataatacgaaaggtggacaagcaaacagcgattcttgacgcggacgacccagtatcacaactacacaagtgcgccttctacctaaaagacacagaacggatgtacctttgcctctcccaagaacgaatcattcaattccaggcaaccccgtgtccaaaagaaacaaacaaagaaatgatcaacgatggtgcttcatggaccatcatatcaactgataaagcggagtacaccttttgtgatggtatgggaccaacagctgacccagtaacacctgtaccgaatgtacatagcctccagttaaacggcggaggagacgtggccatgctagaggtgaatggtgaatgtttcacttccaacttgaaggtgtggttcggtgaaatcgaggccgacacgatgtttcggtgtgcagaaggtctgttatgtgtggtgcccgacatttctgctttccgggaaggctggaagtgggttaaggaatctgtacaggttcccatcaatcttgtgcgaaacgacggagtgatctaccccaccaacctcacattcacatttacccctgagcctggacccaggcaacactgtcccgcagctctcaacatactgcacggtagtaagaggccaacatccatgcccccgactccagtatccgggtctgaggacgacagtggtcgcggaaacgagtcggatcgcggcgatccgatcatgccaataaaacggcccgcgcttgatgtacacgggagacctgtcgcccctgaagcggcagccacgatgaacggtgcgaacatgcttcgtactgcatcgtga

>Cas9::CionaGeminin-Nterminus (from Song et al. 2022, which originally described it as having the Geminin sequence from human instead)

nls::Cas9::nls (described in Stolfi et al. 2014)

Ciona robusta Geminin N-terminus

atggctagccccaaaaagaagaggaaagtggacaagaagtattctatcggactggacatcgggactaatagcgtcgggtgggccgtgatcactgacgagtacaaggtgccctctaagaagttcaaggtgctcgggaacaccgaccggcattccatcaagaaaaatctgatcggagctctcctctttgattcaggggagaccgctgaagcaacccgcctcaagcggactgctagacggcggtacaccaggaggaagaaccggatttgttaccttcaagagatattctccaacgaaatggcaaaggtcgacgacagcttcttccataggctggaagaatcattcctcgtggaagaggataagaagcatgaacggcatcccatcttcggtaatatcgtcgacgaggtggcctatcacgagaaatacccaaccatctaccatcttcgcaaaaagctggtggactcaaccgacaaggcagacctccggcttatctacctggccctggcccacatgatcaagttcagaggccacttcctgatcgagggcgacctcaatcctgacaatagcgatgtggataaactgttcatccagctggtgcagacttacaaccagctctttgaagagaaccccatcaatgcaagcggagtcgatgccaaggccattctgtcagcccggctgtcaaagagccgcagacttgagaatcttatcgctcagctgccgggtgaaaagaaaaatggactgttcgggaacctgattgctctttcacttgggctgactcccaatttcaagtctaatttcgacctggcagaggatgccaagctgcaactgtccaaggacacctatgatgacgatctcgacaacctcctggcccagatcggtgaccaatacgccgaccttttccttgctgctaagaatctttctgacgccatcctgctgtctgacattctccgcgtgaacactgaaatcaccaaggcccctctttcagcttcaatgattaagcggtatgatgagcaccaccaggacctgaccctgcttaaggcactcgtccggcagcagcttccggagaagtacaaggaaatcttctttgaccagtcaaagaatggatacgccggctacatcgacggaggtgcctcccaagaggaattttataagtttatcaaacctatccttgagaagatggacggcaccgaagagctcctcgtgaaactgaatcgggaggatctgctgcggaagcagcgcactttcgacaatgggagcattccccaccagatccatcttggggagcttcacgccatccttcggcgccaagaggacttctacccctttcttaaggacaacagggagaagattgagaaaattctcactttccgcatcccctactacgtgggacccctcgccagaggaaatagccggtttgcttggatgaccagaaagtcagaagaaactatcactccctggaacttcgaagaggtggtggacaagggagccagcgctcagtcattcatcgaacggatgactaacttcgataagaacctccccaatgagaaggtcctgccgaaacattccctgctctacgagtactttaccgtgtacaacgagctgaccaaggtgaaatatgtcaccgaagggatgaggaagcccgcattcctgtcaggcgaacaaaagaaggcaattgtggaccttctgttcaagaccaatagaaaggtgaccgtgaagcagctgaaggaggactatttcaagaaaattgaatgcttcgactctgtggagattagcggggtcgaagatcggttcaacgcaagcctgggtacctaccatgatctgcttaagatcatcaaggacaaggattttctggacaatgaggagaacgaggacatccttgaggacattgtcctgactctcactctgttcgaggaccgggaaatgatcgaggagaggcttaagacctacgcccatctgttcgacgataaagtgatgaagcaacttaaacggagaagatataccggatggggacgccttagccgcaaactcatcaacggaatccgggacaaacagagcggaaagaccattcttgatttccttaagagcgacggattcgctaatcgcaacttcatgcaacttatccatgatgattccctgacctttaaggaggacatccagaaggcccaagtgtctggacaaggtgactcactgcacgagcatatcgcaaatctggctggttcacccgctattaagaagggtattctccagaccgtgaaagtcgtggacgagctggtcaaggtgatgggtcgccataaaccagagaacattgtcatcgagatggccagggaaaaccagactacccagaagggacagaagaacagcagggagcggatgaaaagaattgaggaagggattaaggagctcgggtcacagatccttaaagagcacccggtggaaaacacccagcttcagaatgagaagctctatctgtactaccttcaaaatggacgcgatatgtatgtggaccaagagcttgatatcaacaggctctcagactacgacgtggaccacatcgtccctcagagcttcctcaaagacgactcaattgacaataaggtgctgactcgctcagacaagaaccggggaaagtcagataacgtgccctcagaggaagtcgtgaaaaagatgaagaactattggcgccagcttctgaacgcaaagctgatcactcagcggaagttcgacaatctcactaaggctgagaggggcggactgagcgaactggacaaagcaggattcattaaacggcaacttgtggagactcggcagattactaaacatgtcgcccaaatccttgactcacgcatgaataccaagtacgacgaaaacgacaaacttatccgcgaggtgaaggtgattaccctgaagtccaagctggtcagcgatttcagaaaggactttcaattctacaaagtgcgggagatcaataactatcatcatgctcatgacgcatatctgaatgccgtggtgggaaccgccctgatcaagaagtacccaaagctggaaagcgagttcgtgtacggagactacaaggtctacgacgtgcgcaagatgattgccaaatctgagcaggagatcggaaaggccaccgcaaagtacttcttctacagcaacatcatgaatttcttcaagaccgaaatcacccttgcaaacggtgagatccggaagaggccgctcatcgagactaatggggagactggcgaaatcgtgtgggacaagggcagagatttcgctaccgtgcgcaaagtgctttctatgcctcaagtgaacatcgtgaagaaaaccgaggtgcaaaccggaggcttttctaaggaatcaatcctccccaagcgcaactccgacaagctcattgcaaggaagaaggattgggaccctaagaagtacggcggattcgattcaccaactgtggcttattctgtcctggtcgtggctaaggtggaaaaaggaaagtctaagaagctcaagagcgtgaaggaactgctgggtatcaccattatggagcgcagctccttcgagaagaacccaattgactttctcgaagccaaaggttacaaggaagtcaagaaggaccttatcatcaagctcccaaagtatagcctgttcgaactggagaatgggcggaagcggatgctcgcctccgctggcgaacttcagaagggtaatgagctggctctcccctccaagtacgtgaatttcctctaccttgcaagccattacgagaagctgaaggggagccccgaggacaacgagcaaaagcaactgtttgtggagcagcataagcattatctggacgagatcattgagcagatttccgagttttctaaacgcgtcattctcgctgatgccaacctcgataaagtccttagcgcatacaataagcacagagacaaaccaattcgggagcaggctgagaatatcatccacctgttcaccctcaccaatcttggtgcccctgccgcattcaagtacttcgacaccaccatcgaccggaaacgctatacctccaccaaagaagtgctggacgccaccctcatccaccagagcatcaccggactttacgaaactcggattgacctctcacagctcggaggggatgagggagctcccaagaaaaagcgcaaggtaatggccacgaaaaatattcttcaaaatataaatgcacaatggaaggagaatgacaacagatcaccaagtagaaagcgacggttagatgacgtcactgaagaatcacaattaccttccacgaccaaacgacgtcatcttcaaacaaatacaaacgttgtaaattccacaggattgaaacaaggcctgacaaatgtgaaaaattcaataaatccaaagaacaaatcaataaaaaatttcttttctgatattccacgtgtgtcatgtactaaatctgaaaagattcaaatttttaaagaagctaagaaaactccaaaaaagaatgcaaccactcagacaaggagtgaagctgaagaattggtctgcagtgatcaacccagtgaaaaatattgggaactcttagccgaggagcgaaggaaagggttgtaa

>KH.C14.116 *in situ* hybridization probe (coding sequence)

atgaaaggctcaatgttgatttgttttatcgttgccagcacttcatactttgaactaaccaaaggtttatcctgttggacttgtattaacgcaagaagcaatgcggagtgtcaggctacaggtcatctgcaacaatgccgatttacccaaagagcttgtcaaactcacatacgaaccgatccaatgggtattcgcatcacgaaggaatgtaaacaagtacaagcgtgcaccaacaactttttgcagaatccgagaccagcctggtacccgagtcaatgcaacgataatgtacaaggatccgtatgtcgatgctgctgtgatttcgacaactgtaactttgagtctgtagcatgcccgggaagtcgcaccacaactactctagcaccaacaacgacaacaacaacccaaccgctcacgcaattagtggacgtgttaaacgttggaccggaggagccaaaaacctgtgacaagataacgttacgaaacggattcgtcgcttgcactgatgagaataagcacgactcgctatgtatgttccaatgcgacgttacgagaggatacgaactactgcctttaacgttacacagacgacgtgcaatgcaactgtatggtggccgaccccgccttgctgccagcggccctgtcctccttatgcgaatgcggatattcttgtgattctggattcgtcatcgtcgattggacgttttaactgga

>KH.L96.43 *in situ* hybridization probe (coding sequence)

atgtatcgaattaaggtgtgtctggttttgtgctttatctgctgtatagtatcagcacaagataacgtagattcgtattggggagaatggcaagaatggtctgaatgctccaacgactgtggtgtcgggattcgggagagactcaggccatgcgagggcgctggagagtgcgactatacttgcgatggttgcgaagatttccaacaagaggattgtttgaatgaacagccttgtccagaatggtcaccatgggaagctgtggacgattgcaatgctacttgtggttgggggaaccaaacaatgacaagaacatgcgatgacgtaggttgtattggggaaacggagaagatagaagcttgctgggttgaatgcaaatcttgcgataatgagatgtcagaaacagattgcgaaacgtataaagtttggtgccagatttccccaacactaatgactcgttactgccccaaaacttgctcaatgtgcgatcggacatgttcttgtcaaactaaatggtcaagttggacaccatgcactttaacttgtggcggtggtgtcaaatacagacaacgggattgctccaatgtagaggatagatgcggcctctcatccaacatggaggtcccatgcaacacacaggaatgtagaaattggagttcatgttcccgtacgtgcggttggggtacacggcataggagacgggtttgcccaccgaataacccgcttatatgcatcaagacacagatggaatatgaagactgcaagaatatggactgcccaccaaagagttgtccacgcggttcattcccaaggtatggaaccaacacaaaatgttgcacacctgataatgaaaactgcggtagtttcttcgggaatgacaacgatatagtgggaggggtggcagcggataaaaggtcatggccgtggatgttaagattaagattgggaaacccgagaggtggatcgtttgcttcgtgtggagcaagtttaattcacagtcgttgggcgatcactgccactcattgtattctaacattcaccggtccacggcaaatccttgcgattgcttcgcagacaagatacgtgcataaggtttacaaacacgaaggcttctctagctggttggagaatgatatttcattgcttgagtttaaggaaccattcgagttcaacccggagacaaacattgctccggtctgtcttccctggaacgaagagacaccggacgatacgcagtgtttcgctgcagggtggggagcaactgatccagcggatttacgtccgagcaatacgctacagcaagtgagacttcgtaatttgaacatggacgtatgccgctatgcgtatgaccaagcacctggcgacccattccgtattccaaacgctgataaactgaacgaaactacgatgatatgcgctggtcggttaagagggggtgcagatacttgtgcgggcgacagcggaggacctttaatgtgtcaaaggtgttcaacatgttcgtggtacattgctggtataacttcctacggtactgctaactgtggggcttctggtcggcctggtgtatacactaaagtcctggcatatgaaccatggatacgagagaccacacgaggagaaatacaaaggattcgagactttagaggaagttgttcaatacaaaacacactgcagtaaaacttgaatctcgttgtaatggcc

>KH.C2.1013 *in situ* hybridization probe (partial coding sequence)

tgctcatttacttgcgatgctggatttgctttggtcggtgcggaagtttcaacatgcttggatgattttgatggaaacttgctgggcgcttggagttctattgccccgacttgccagccaattgtgtgtctaccgccgcacatcaatccagccgacggtgcagtttcttgcacgaattccaatttcatggcgagccagtgttcttttagctgctctgtcggattcgcattggtcggtccaccagtttcaacttgcaacgatgactttaacggagatacgttaggcgcgtggaacaacatcgcacccacttgccaacccattacctgtattccgccacacacctcaccagagaacggcgcagtcgtgtgtagtaatgctaacttctacgccagcgaatgtgtgtttgaatgcgacaccgactatgttttaactggagaatcagcttcaagctgtttggaaggagtgcccggagatactgtcggcgcgtggagctctcttgcgccaacctgtgaacgaaagcaatgcgcagcacccccaagccccccgactaacggcttcagagtgtgcacagacggcaactttatcggctcccagtgcgaatacggatgtgacgcataccacacccgggttggtccactttactctacatgcgtagaagggtcggatggaaaccttgtttttgacaaccccgcacccgagtgtcaacctaaccagtgtccagagcaagggccacttagacacggtcgtatgacgtgttcggatgcgaacaacgccccatcaacatgcttgtttcaatgtgtggatcctggttaccaactctacccagcagatatgacacaaaacacttgcagaaacgactcacagtggaatttaccgaaaccttgttgcgctcgaccttgtccaccattcgctttaatggatgctgttttcatacttgattcgtcgtcgtccattggaaccgcaaactgggtcactatgaaaactttcgtccgcaacgtactcgggtcattcgtgctggctccagacgcagctcgtttctcagtttttagatacaaccgtcatgttgacaacaccactcaaattcttctcaacgagttcgaaaatgacatcgacctctttttaaacaagtttgatgacatcccatatgatggcagtggaacctggacaggtcaagcactgagacacgccaaagacaccattctgctcccgggtaatggaaaccgacccggagtcaaagacgttgtccttattattacagatggtcgctcccaagatgacgtcagagaagtttcccagc

>KH.C11.360 *in situ* hybridization probe (coding sequence)

atggcgtggcctattaacttagttaagaacattattgtagtaattgtgttgctatgttatgtcagctccacgttgggttgtcacatcccaggccacacaaataagcgagccataagcctagccgcatcggtagaaatgaccgacgtccaaaaaattcgaaactttttcatcccctgcgttaatgacgtgtttggtgacgccagcttggatttttctggaatcgttaagtttataacagaccactcgctatggtatgacgtcacacagcgtctggaatgcggggagaaaaacgatactgtcgaatgttggaacgaacgtcttactgatgttttgacccgaggttcgaaagaacttgctgtggcacaggagtgtgtggagcgatataaagactttgttaataatggaaaatacaatcctaccccgttggtttttaataaattcgaccgagtgaggtccgaagatctgttgaactgtcccactcgaacgccacgaaaattcgatctttgcgtgaataaccaattcgccaaagatgaggtcgagaatccaatcggaaattttgattctttgattggatttgttcaaaggaagatgaaatgcagtgactgtgattgtggcgaggttctacaagccgacagtgacttcgcatcgttcttcgtgggcgagggactacaatgcgactcttgttggtcaactagactggcagcattggtcaaaagtcaatcgactttaagtgaaacaaccaaaaagtgcgctgcaactaccaacgacgaagtttttttgaaagaggaaacaatgagagtatactaggtcatactctcgctgtatccgtaggag

>KH.C11.249 *in situ* hybridization probe (partial coding sequence)

atggacaagcggattctcttcgtatcattgtgcgcatgcgcagtagcctctggactgtacattcaagaccccgaacccaatgttgattgtcaattatctgaatggggggaatacgggccgtgcacgaatgttggtgggtcacaacgacctgtcttaattcgtataagacgccgagagattctggttcgtccctcgggaaatggcaacgactgtactccgcgggaggaagtgtcaggttgtggggccgaccctactactgctgagccacaaactggatacccaaacaatctcccaacatcggttccacctaagaagaaaaagaagtgtagaaagggaaagaagggacgtaaaggaaaaaaaggtccccgaagatcaagcgggaagtccggatgctccgactccgataatgacgatgatgataagaagagaagaagacaaaataagaggaaaggaaagaaacaaggatccggatctggaagtgatagtgatggttattctgggaaccaatcagattaaagacacattgtgacgtgtagtagttaagtgtgacgtataatggtgggaatttatgttagattggtcatgtagtgattaaagaagtcaaaaaattgtggcgtagttgcagaagaaatcagaacaacgctttttgttttaaactcacttattatgacgtcattaaggcgtctttccagaagcgcacgaaacaatcaacaaaatgtgcttgcg

>TGFB (KH.C3.724) *in situ* hybridization probe (coding sequence)

atgtggaaaaggaacacaagggcgtctgtgacgtcatggtcgtcattattaactctattttcaatctcgtttatgtttcttgcaccacgaacaatggcggggtgtggggttaattttgataagatgaggaagcgcaggatcgaggctgtgcgaggccagatattaagcaagcttggccttacagaactccccagcgccgccgcaacgccgcgcgacgttcctagggaggtcgaggctttgtacaaccgcacgcgggactttgtgctcgagcaagcccgccaacagaggcaagaatgtctcgatcctgaggagacctactacgcagaagatgtattgaccgtgtatatgaagaacgtacaaccacagtcggcgctccaaccagatcgctacaagggttacaaactatcaaggggactttcggagttttacgacttcgatctcacagcaacacaagtcaacccagactcaatagtgtcggcaacattgcgattatatcaagtacagaaccctggtagtaggggggaaaggaaccaggttgaactgtatcaactacaacctccagaaaaggaagggttgacgcctgtcaagaggtttctcgatatgaaactaatggacacgggtgtggaagcatggcagtcgtttgacgtaacttcaactgtgagagaatgggttcaattccctcacttaaacgacggtcttgaactcacgataccgtgtctcgacgaagacaacttttcagggacccaaaaacgacgaagtttgggacccaggaaaaccttggatgtcattatagctggcccgggtgggactcgaaccatgagcagtcgaggtgaccaggaccccgaaaacttcaccccggaccttgggagggagttttatccccacctagtgatcatggtgaggacccctgctaaccaggggagctctcgtaccaccaccagcagccgcaggaggcgcaagagagcattagatgcggattactgcttcaaccgaaatccgtcagagacaaattgttgtttgagggaattatacattgatttccgcagagatttggaatggaattgggtgcgagcacctgttggttacaaagctaacttttgcgctggtgcttgtccgtatttatggagtatggacacccaacatgctactatacttggtctgtataaaagtatgaacccgcacgcttcttcagcaccctgctgtacacccaaagaactggaccctcttatattgatgtattatgccaacaatgagtttaaattcacgaaaatgtcggacatggttctcctatcttgcaaatgcagctga

>KH.S908.2 *in situ* hybridization probe (partial coding sequence)

cttccatccaacatcacacggtcgcaccagtgtcattaacatcatggtggaaccgaactcctatttccaaaagagtggatcttcatacaatggcagggttcgatccagcatcacatacattgatcctcgagatgtttcatcttatctagaggcaccaagtgatctccgaacagaagatggaagagtgatgaggtcatatgggatgtttgaacttgattttagaaacttccagaattacgaactcgatgtcagaaatattcgaatttctcttcaagctagtgaggtgggtattctttctgaacaacgacaaatgaggatgtggtctcttaatccagtcaacggactgtggtatgatgaaggagaaatggctctgatatcatctgcgcaaggaatgcaattaactgctaacattgctttggttcaaactcgaagattattaaacatcgacgttgcttcgccaagaacttgctatgctaaagtgcgtgcatatggatctgatgcattcaatatctgggatcaagtttcctacatcttggtggatgctgtcttccgagaagccactgatagtagatggaggaactttggatcatcactcacgcttcctaactcaggagcatgcttaagagtcttttgtgatgttgctgacagcaatgctttcactgcaacatttactgcctcactaaatgctgagaaaagcttggatgcagcttcatctattaatcaagattccgctatcattggggtgtcagatgaaattctaccactcatcaattacaggaggtctggtcatggaggaagtcaccaagacacaacgtcgttcaccatcaatttaccacaaacttccaatgtaacacatttggatggacctttctacagagaccaatttgattgtttaactgcaccagttactcatgatcatttcagattctcaatgaaaccaagtgcactatttgagtttaacacaatacccattgatgagaataacattttctttacttggcaaaatacacctctgatttggtggccaagaccaaatcaatttagatcatgctatatgaaggtgaaaattattggaacggcaaacgcagttgtagtggagagcagaaatactgttggttcgagacaagacgttcttcaccagttgtatggaattcgggaagatagtagcgttagtaatggagtggaatctacagcttgtattgaatttaagtgcagtggagtttcgctggaccagcaaacagatcaaacggtcgtggatataacaccacgtggctcatgttctcgtacaaatgtcaacgtgctcttaacgaactacctcaccgaacgtaacagcctggctgaccaagtctctatcgcaagcttcaagatgtttgcacctaccgatccactcggacataactacggtatctatacagtcaccaccagcgatcctatcacatcgaagcagttgtcaaagggcagatgtatggctggccaatcaagtatcacttcttccgtaatgcaagttgataataacgtcgcggtgactttcaactgcggatcccattaatttatacctaataccaaaagcatattaaccacgcgg

>Astl-related(KH.C9.850) *in situ* probe template (coding sequence)

atgctctttacgaaagaattgtcgtttgcaatggttttgttggtgcacttggttcaggcagtacctgttgtaaacttagtgaacatgccggacaccgaactgtgtgctttaatcaggaacggcacaaacgttgcatttgcgatggaagaggcagaaaaatgcaatgatacgcaagacgatttgggcgccggatgtccttctgtagaatgcgacaacagtttactttgctctaacggctacttattagatgcagaaggctgcgagacatgcgagtgcatcgaatgggcacgagatatggaaatggacattaccggagactttgacgacattgatgtgtggaaagattacactgatgagctcggagaatttgtcataacatatgaaacatctgaatacctaagttcagccgagaagaaagcaattacgcaagccattgacgaaatcaacgagaacacatgccttcaatttttaccccgaaagatcaatgactcaaaggaaaattacataaggtttgaatgggatgaagtctgttcatctgcagttgggaagactggtggagcacaaggagttattctaagtgaagagtgctccacgaaaggcggtgttctgcacgagttgatgcatacactcggtttcctccatgagcacaaccgacccgatcgagatcagtatattaatattgaatgggagaacattatacctgaaaaacgaagcatgttcaacaagtggcaggtttcagaagacggtgtattgaatgagttcccgtatgacgtaagctcggtcacgcatcttgattcctttgccttctccaacaatggtgaacccaccattacagggctggatggctttccacttccaacacaaaacactcatctgtctgtgctagatattgacaaaattaaccacgtgtatggttgccctgagttcatgcctttgccgcccatgatgccagacaaatgtaaagatgatttatttgcttgtcccgaattggcgaacacttgcgcatgcttattgtcacccgcattcatgatgaatcactgctgcagctcttgcgagatggaggcgacacagatcgttgaccattttcccttctgcaacttatgggcagcaaactgtggtaaaagtagatatttcgacaacctttgccgcaagacgtgtccgaagtgtgttctggctgggtag

>Villin (KH.C9.512) *in situ* probe template

aacaaggtgctgccgctatgctggctacacaacttgacgactatcttggtggggacccagttcaatacagagaaacacagggcaacgagtcgacgatgtttaaggcctacttcaaaagtggtatcgtctactgtaagggtggagtagcatctgggtttaaacacgtcgaaaccaaccagtatgatgtccgacgtctattgcgcgtaaaaggaagaaaaactgtgaatgcgaccgagcaggactttgcttggacatcatttaacctgggcgatgtatttcttgttgatttgggcaagattatcattcaatggaacggacctgagagtaatagaatggaacgattgaaggctaccatcctggccaaagatattcgtgatcgtgagcgtggtggacgaggtcaagtccttatcgtggatggggagaacgagaaaacaagtgacaaggcgtatggggccatgctgaaattgttgggcgacaaaccaaaactgaatccagctataccagatgaaattgcaagcaggaacaaactcagtcagttgaagcttttccat

>Foxg *in situ* probe template (coding sequence)

atgacgaacgacgcggccgagtctggtcattcaaagcgagaaaacttcatagagatgtcgcccgagtattcaacgctgatcgccgccgaagatcgaagctcagttccccgcagagaggacgcgcaagtgccgaggtttgagaatatacaaaatggtggtgccgatatagaaaacggtgaagtgtctccaatacaaaaagatattgcaaaccaagagctcaatgacgtagcaattaatatgacgtcacctcagaaacaaacaaacaacgaaaatttagaagaaaaatgtccaaaagaccagaaaccgtcaacaagtccaccgagtaacaagtacggtaagaaaccgccatattcatacaacgccttgataatgatggccatcaaaaagagcccacgaaagcgacttacactaagtcagatctaccaatacataacaactacattcccatactacaaagaaaataaacaggcgtggcagaattctatcagacacaacttatcgttgaacaaatgctttgtgaaagtaccgaggcactacgacgaccccgggaagggtaactattggatgctagacccgtctagtgatgatgtatacattggtagtagcaccggtaaactgagacgaagaagttcaagcagtcaagcaaggggtcgtttagcattacgacgaagaacattcgcccaagtatttggatcgccgcaagatattttacaacacgaccccccacaacagataataagaacagacgtgacgtcacgagcggccatgttacgtcatcatgacgcagcacgaacaatgctgggttctggaatcccacagcgggcagatccctaccctatgtttcaacacacgcggttccctatgactacagaaactgattcaagatacaggcaattgtatagagcgagattggaacaatactacgcgcatctcgcgtcctctgcactgttcgggcacatgcaagctactgcattaaacgcccagccacgtattgttaaaagcaaccccgcctctcctgagcctgtgcattcccattacgaacaaactacgtcaccaaacaccgcacctcctcgctctgacacctccacaccaccgcgagggatcgaacgaccgtgggcatcaccgccgcgtaggaggtgtgacgtcacaaagcatgaaacaattgaccgcgacgtcgtgtcgcgatgctcctccgaatcttctaatgaatcatcaagaaacaagagagaagtcaccgaaaactctacgaattcaccgatacagcaacttagtacaagaggagggttaccgttctaccttacaccaaccccgaacccttgccctaatttcttaatgcccaacactgccgggttagttccaaatccaagttaccccttctttttcccccaaccattccacccagccttggcgtttctctctcggccacaaactgttgcgtcatcatcgcaattgtga

>Ascl.a (KH.L9.13) *in situ* hybridization probe (coding sequence)

atggcgaccggaagtgacgaaccgcggtcgaacgcgataatatcgatgccgttcaacaacagatggaaggagggaagtttgggggaaaataacccaaacttccgagttgagattcggagtggggctggggggaaaaaccccaccagtgtagcgaggcgaaatgcacgggaacgacgaaggattaaaaacgtgaattcagcattcgacgaattgagacaacatgtgcctaatggtgaaagaaatcggaagaagattagtaaggtggacacgttacaatctgctatcgaatatattaaagcattagaagaactcgtgcgtaaccggaagtccaaaagtgacgtcatcaataaagagaacgctacaacgtcatcgaacgctatgacgtcacaaaacgatgacatcatgtttgtaaaggaagctgaagtgacgtcacagaagaaaaaagattcgaaatctccggtagcgttaacggagtctatgctaaaggcgtttgacgtcatgctgcaaaaatgtacgtcacaatcgaagacaaaagaagatgatgacgtcataaggatggattcgacgagcgacagcggtttctccgagatcctctgtgatgtcacaagcggcggggaatcgatgacgaatttgccgccaaatattccagaatccccgatcttacaatcgaacaattgtgacgtcagttttgaatcgttgcaggggtttcccccaagttacaacccccaacgttttaccccttaccccacgtacatgggtgagtggtcccacatgccccctaccccaggagaattccccccagtagaccaaattaccccaacccttcaacttccggttggattttcaaacgattttcccgccaatgacgctgattggttaaaccacaatcatttctgatcaccacgtgaccttaaccga

Pou4 *in situ* probe template: Used clone citb034g05 from Nori Satoh gene collection (R1CiGC32g05).

**sgRNAs used in this study:**

Islet.2 (from Gandhi et al. 2017)

**GGATGCGGCAGGGGTCTGTA (G+N19)**

Islet.1

**GAAGATGGCGGACCCTCATGG (G+N20)**

Sp6/7/8.3.29

**GACCCCACATCTCCTATAGG (G+N19)**

Sp6/7/8.7.117

**GTACACTGCCCCCAGCCCCG (G+N19)**

Pou4.3.21

**GCTGAGTGGTGGAAAGCGGG (G+N19)**

Pou4.4.106

**GAGGATGGAAATGATTCGGG (G+N19)**

Foxg.1.116

**GAGCTCAGTTCCCCGCAGAG (G+N19)**

Foxg.5.419

**GTGGAGGTGTCAGAGCGAGG (G+N19)**

Villin.4.74

**GTTCAATACAGAGAAACACA (G+N19)**

Villin.5.105

**GGTCCAAGCAAAGTCCTGCT (G+N19)**

Villin.18.72

**GGATCCGATTGTTTCCAAGC (G+N19)**

Control (from Stolfi et al. 2014)

**GCTTTGCTACGATCTACATT (G+N19)**

*PCR primers for amplicon sequencing by NGS*

| **Gene + exon** | **Forward primer** | **Reverse primer** |
| --- | --- | --- |
| Sp6/7/8 exon 4 | ACTTTTGTGCGACTATTTTCTG | TAAAATCGCTAATACGGTACAC |
| Sp6/7/8 exon 8 | CCCCAGCCCAACATCATT | GTCCGCCAAAATGTTTAGC |
| Foxg exon 1 | GCCGAGTCTGGTCATTCAA | TCATTGAGCTCTTGGTTTGC |
| Foxg exon 5 | TACTGCATTAAACGCCCAG | TTAGAAGATTCGGAGGAGCAT |
| Pou4 exons 3+4 | CCGTACCACCACCAGGTA | TGCAGAGATGGGCGTGAC |
| Islet exon 1 | CTACAAGCGGAGAATTTCGAA | CGGGATCCCTGGATTTTC |
| Villin exon 4 | CCCATTTCGCTTTAGCACA | GTATCAACTCTTACATAGCACCG |
| Villin exon 5 | CTGTCACGACGTTTTAAATGC | ACATGTTTGTCATATACCAGAACAG |
| Villin exon 18 | CAGCAACATTTAAGCGACAGT | CGAAAGTGGCCAAATAATCAG |

**Electroporation mixes for perturbation experiments**

**(per 700 ul of total cuvette volume)**

Islet CRISPR to look at ACCs (CryBG reporter):

40 ug Foxc>Cas9

10 ug Foxc>H2B::mCherry

50 ug U6>Islet.2

35 ug CryBG>Unc-76::GFP

Negative control for the Islet CRISPR/ACC experiment above:

40 ug Foxc>Cas9

10 ug Foxc>H2B::mCherry

50 ug U6>Control sgRNA

35 ug CryBG>Unc-76::GFP

Islet CRISPR to look at ICs (C11.360 reporter):

40 ug Foxc>Cas9

10 ug Foxc>H2B::mCherry

50 ug U6>Islet.2

70 ug C11.360>Unc-76::GFP

Negative control for the Islet CRISPR/IC experiment above:

40 ug Foxc>Cas9

10 ug Foxc>H2B::mCherry

50 ug U6>Control sgRNA

70 ug C11.360>Unc-76::GFP

Islet CRISPR to look at OCs (L141.36 reporter):

40 ug Foxc>Cas9

10 ug Foxc>H2B::mCherry

50 ug U6>Islet.2

70 ug L141.36>Unc-76::GFP

Negative control CRISPR to look at OCs (L141.36 reporter) as above:

40 ug Foxc>Cas9

10 ug Foxc>H2B::mCherry

50 ug U6>Control sgRNA

70 ug L141.36>Unc-76::GFP

Islet CRISPR to look at PNs (TGFB reporter):

40 ug Foxc>Cas9

10 ug Foxc>H2B::mCherry

50 ug U6>Islet.2

70 ug TGFB>Unc-76::GFP

Negative control for the Islet CRISPR/PN experiment above:

40 ug Foxc>Cas9

10 ug Foxc>H2B::mCherry

50 ug U6>Control sgRNA

70 ug TGFB>Unc-76::GFP

Islet>Islet to look at ICs (C11.360 reporter):

70 ug Islet [intron 1, 0 to 2014]+[-473/-9]>Islet

35 ug Islet [intron 1, 0 to 2014]+[-473/-9]>H2B::mCherry

70 ug C11.360>Unc-76:GFP

Islet>Sp6/7/8 to look at ICs (C11.360 reporter):

70 ug Islet [intron 1, 0 to 2014]+[-473/-9]>Sp6/7/8

35 ug Islet [intron 1, 0 to 2014]+[-473/-9]>H2B::mCherry

70 ug C11.360>Unc-76:GFP

Islet>Islet to look at ACCs (CryBG reporter):

70 ug Islet [intron 1, 0 to 2014]+[-473/-9]>Islet

35 ug Islet [intron 1, 0 to 2014]+[-473/-9]>H2B::mCherry

35 ug CryBG>Unc-76:GFP

Islet>Sp6/7/8 to look at ACCs (CryBG reporter):

70 ug Islet [intron 1, 0 to 2014]+[-473/-9]>Sp6/7/8

35 ug Islet [intron 1, 0 to 2014]+[-473/-9]>H2B::mCherry

35 ug CryBG>Unc-76:GFP

Foxc>Islet to look at ICs (C11.360 reporter):

50 ug Foxc>Islet

50 ug Foxc>lacZ

10 ug Foxc>H2B::mCherry

70 ug C11.360>Unc-76::GFP

Foxc>Sp6/7/8 to look at ICs (C11.360 reporter):

50 ug Foxc>lacZ

50 ug Foxc>Sp6/7/8

10 ug Foxc>H2B::mCherry

70 ug C11.360>Unc-76::GFP

Foxc>Islet+Sp6/7/8 to look at ICs (C11.360 reporter):

50 ug Foxc>Islet

50 ug Foxc>Sp6/7/8

10 ug Foxc>H2B::mCherry

70 ug C11.360>Unc-76::GFP

Foxc>Islet to look at ACCs (CryBG reporter):

50 ug Foxc>Islet

50 ug Foxc>lacZ

10 ug Foxc>H2B::mCherry

35 ug CryBG>Unc-76::GFP

Foxc>Sp6/7/8 to look at ACCs (CryBG reporter):

50 ug Foxc>lacZ

50 ug Foxc>Sp6/7/8

10 ug Foxc>H2B::mCherry

35 ug CryBG>Unc-76::GFP

Foxc>Islet+Sp6/7/8 to look at ACCs (CryBG reporter):

50 ug Foxc>Islet

50 ug Foxc>Sp6/7/8

10 ug Foxc>H2B::mCherry

35 ug CryBG>Unc-76::GFP

Foxc>Islet to look at OCs (L141.36 reporter):

50 ug Foxc>Islet

50 ug Foxc>lacZ

10 ug Foxc>H2B::mCherry

70 ug L141.36>Unc-76::GFP

Foxc>Sp6/7/8 to look at OCs (L141.36 reporter):

50 ug Foxc>lacZ

50 ug Foxc>SP6/7/8

10 ug Foxc>H2B::mCherry

70 ug L141.36>Unc-76::GFP

Foxc>Islet+Sp6/7/8 to look at OCs (L141.36 reporter):

50 ug Foxc>Islet

50 ug Foxc>SP6/7/8

10 ug Foxc>H2B::mCherry

70 ug L141.36>Unc-76::GFP

Foxc>lacZ (negative control) to look at OCs (L141.36 reporter):

100 ug Foxc>lacZ

10 ug Foxc>H2B::mCherry

70 ug L141.36>Unc-76::GFP

Sp6/7/8 CRISPR to look at ICs (C11.360 reporter):

40 ug Foxc>Cas9

10 ug Foxc>H2B::mCherry

25 ug U6>Sp6/7/8.4.29 sgRNA

25 ug U6>Sp6/7/8.8.117 sgRNA

70 ug C11.360>Unc-76::GFP

Negative control CRISPR to compare to Sp6/7/8 CRISPR/ICs above:

40 ug Foxc>Cas9

10 ug Foxc>H2B::mCherry

50 ug U6>Control sgRNA

70 ug C11.360>Unc-76::GFP

Foxg CRISPR to look at OCs (L141.36 reporter):

40 ug Foxc>Cas9

10 ug Foxc>H2B::mCherry

25 ug U6>Foxg.1.116 sgRNA

25 ug U6>Foxg.5.419 sgRNA

70 ug L141.36>Unc-76::GFP

Sp6/7/8 CRISPR to look at OCs (L141.36 reporter):

40 ug Foxc>Cas9

10 ug Foxc>H2B::mCherry

25 ug U6>Sp6/7/8.4.29 sgRNA

25 ug U6>Sp6/7/8.8.117 sgRNA

70 ug L141.36>Unc-76::GFP

Pou4 CRISPR to look at OCs (L141.36 reporter):

40 ug Foxc>Cas9

10 ug Foxc>H2B::mCherry

25 ug U6>Pou4.3.21 sgRNA

25 ug U6>Pou4.4.106 sgRNA

70 ug L141.36>Unc-76::GFP

L141.36>Unc-76::GFP to check for colocalization with PNA staining:

50 ug L141.36>Unc-76::GFP

Foxc>Sp6/7/8 to look at ICs and PNA staining:

50 ug C11.360>Unc-76::GFP

50 ug Foxc>Sp6/7/8

Foxc>Sp6/7/8 + Foxc>Islet to look at ICs and PNA staining:

50 ug C11.360>Unc-76::GFP

50 ug Foxc>Islet

50 ug Foxc>Sp6/7/8

U0126 vs. DMSO to look at PNs:

50 ug C4.78>Unc-76::GFP

20 ug Islet [intron 1, 0 to 2014]+bpFOG>H2B::mCherry

U0126 vs. DMSO to look at OCs:

50 ug L141.36>Unc-76::GFP

10 ug Islet [intron 1, 0 to 2014]+bpFOG>H2B::mCherry

Pou4 CRISPR to look at C4.78 reporter expression:

50 ug C4.78>Unc-76::GFP

40 ug Foxc>Cas9::Geminin-Nterminus

10 ug Foxc>H2B::mCherry

40 ug U6>Pou4.3.21 sgRNA

40 ug U6>Pou4.4.106 sgRNA

Control CRISPR for Pou4 CRISPR above:

50 ug C4.78>Unc-76::GFP

40 ug Foxc>Cas9::Geminin-Nterminus

10 ug Foxc>H2B::mCherry

80 ug U6>Control sgRNA

Foxc>SUH-DBM to look at PN specification:

10 ug Foxc>H2B::mCherry

70 ug Foxc>SUH-DBM

70 ug C4.78>Unc-76::GFP

Control for SUH-DBM PN experiment above:

10 ug Foxc>H2B::mCherry

70 ug Foxc>lacZ

70 ug C4.78>Unc-76::GFP

Foxc>SUH-DBM to look at OC specification:

10 ug Foxc>H2B::mCherry

70 ug Foxc>SUH-DBM

50 ug L141.36>Unc-76::GFP

Control for SUH-DBM OC experiment above:

10 ug Foxc>H2B::mCherry

70 ug Foxc>lacZ

50 ug L141.36>Unc-76::GFP

Foxc>Sp6/7/8 to look at PNA staining:

50 ug Foxc>Sp6/7/8

50 ug Foxc>lacZ

40 ug C11.360>Unc-76::GFP

Foxc>Islet + Sp6/7/8 to look at PNA staining:

50 ug Foxc>Islet

50 ug Foxc>Sp6/7/8

50 ug Foxc>lacZ

40 ug C11.360>Unc-76::GFP

Islet CRISPR to look at protrusions:

35 ug Foxc>Cas9

70 ug Islet [intron 1, 0 to 2014]+[-473/-9]>Unc-76::GFP

50 ug U6>Islet.2 sgRNA

Control for Islet CRISPR to look at protrusions:

35 ug Foxc>Cas9

70 ug Islet [intron 1, 0 to 2014]+[-473/-9]>Unc-76::GFP

50 ug U6>Control sgRNA

Islet overexpression for bulk RNAseq:

50 ug Foxc>Islet

Islet CRISPR for bulk RNAseq:

40 ug Foxc>Cas9

30 ug U6>Islet.1 sgRNA
30 ug U6>Islet.2 sgRNA

Negative control for Islet bulk RNAseq:

40 ug Foxc>Cas9

60 ug U6>Control sgRNA

Islet CRISPR to look at Villin reporter expression:

50 ug Villin -721/-1>Unc-76::GFP

10 ug Foxc>H2B::mCherry

40 ug Foxc>Cas9

30 ug U6>Islet.1 sgRNA

30 ug U6>Islet.2 sgRNA

Islet overexpression to look at Villin reporter expression:

50 ug Villin -721/-1>Unc-76::GFP

50 ug Foxc>Islet

10 ug Foxc>H2B::mCherry

Control for above Islet manipulations with Villin reporter:

50 ug Villin -721/-1>Unc-76::GFP

10 ug Foxc>H2B::mCherry

40 ug Foxc>Cas9

60 ug U6>Control sgRNA

Villin CRISPR to look at ACC length:

40 ug Foxc>Cas9

10 ug Foxc>H2B::mCherry

35 ug CryBG>Unc-76::GFP

35 ug U6>Villin.4.74 sgRNA

35 ug U6>Villin.5.105 sgRNA

35 ug U6>Villin.18.72 sgRNA

Negative control to compare to Villin CRISPR above:

10 ug Foxc>H2B::mCherry

35 ug CryBG>Unc-76::GFP

Pou4 CRISPR to look at metamorphosis:

40 ug Foxc>Cas9

25 ug U6>Pou4.3.21 sgRNA

25 ug U6>Pou4.4.106 sgRNA

Islet CRISPR to look at metamorphosis:

40 ug Foxc>Cas9

50 ug U6>Islet.2

Foxg CRISPR to look at metamorphosis:

40 ug Foxc>Cas9

25 ug U6>Foxg.1.116 sgRNA

25 ug U6>Foxg.5.419 sgRNA

Sp6/7/8 CRISPR to look at metamorphosis:

40 ug Foxc>Cas9

25 ug U6>Sp6/7/8.4.29 sgRNA

25 ug U6>Sp6/7/8.8.117 sgRNA

Negative control CRISPR for metamorphosis CRISPRs above:

40 ug Foxc>Cas9

50 ug U6>Control sgRNA

Foxg CRISPR replication to look at metamorphosis:

40 ug Foxc>Cas9

10 ug Foxc>H2B::mCherry

40 ug U6>Foxg.1.116 sgRNA

40 ug U6>Foxg.5.419 sgRNA

Negative control to compare to Foxg CRISPR replication above:

40 ug Foxc>Cas9

10 ug Foxc>H2B::mCherry

80 ug U6>Control sgRNA

Islet>Sp6/7/8 to look at metamorphosis in absence of ACCs:

70 ug Islet [intron 1, 0 to 2014]+[-473/-9]>Sp6/7/8

35 ug Islet [intron 1, 0 to 2014]+[-473/-9]>H2B::mCherry

35 ug CryBG>Unc-76:GFP

Negative control to compare to Islet>Sp6/7/8 experiment above:

35 ug Islet [intron 1, 0 to 2014]+[-473/-9]>H2B::mCherry

35 ug CryBG>Unc-76:GFP

*Ciona robusta* gene model ID table

| **KH gene model ID** | **KY21 gene model ID** | **ANISEED gene ID** |
| --- | --- | --- |
| KH.L96.43 | KY21.Chr4.999 | Cirobu.g00013387 |
| KH.C4.78 | KY21.Chr4.267 | Cirobu.g00006902 |
| KH.C11.360 | KY21.Chr11.1038 | Cirobu.g00002187 |
| KH.L141.36 | KY21.Chr7.130 | Cirobu.g00011190 |
| KH.C14.116 | KY21.Chr14.663 | Cirobu.g00003524 |
| KH.C2.1013 | KY21.Chr2.620 | Cirobu.g00004128 |
| KH.C11.249 | KY21.Chr11.375 | Cirobu.g00002066 |
| Gnrh1 (KH.S1051.1) | KY21.Chr1.1655 | Cirobu.g00013539 |
| Astl-rel (KH.C9.850) | KY21.Chr9.1097 | Cirobu.g00010438 |
| TGFB (KH.C3.724) | KY21.Chr3.1474 | Cirobu.g00005927 |
| CryBG (KH.S605.3) | KY21.Chr1.2268 | Cirobu.g00014792 |
| Islet (KH.L152.2) | KY21.Chr4.1164 | Cirobu.g00011396 |
| Foxc (KH.L57.25) | KY21.Chr12.158 | Cirobu.g00012813 |
| Foxg (KH.C8.774) | KY21.Chr8.693 | Cirobu.g00009441 |
| Sp6/7/8 (KH.C13.22) | KY21.Chr13.415 | Cirobu.g00003422 |
| Pou4 (KH.C2.42) | KY21.Chr2.456 | Cirobu.g00004616 |
| Emx (KH.L142.14) | KY21.Chr8.1320 | Cirobu.g00011241 |
| Ascl.a (KH.L9.13) | KY21.Chr2.1499 | Cirobu.g00013239 |
| MyT1 (KH.C1.274) | KY21.Chr1.1192 | Cirobu.g00000497 |
| Villin (KH.C9.512) | KY21.Chr9.368 | Cirobu.g00010065 |
